# *De novo* design of drug-binding proteins with predictable binding energy and specificity

**DOI:** 10.1101/2023.12.23.573178

**Authors:** Lei Lu, Xuxu Gou, Sophia K Tan, Samuel I. Mann, Hyunjun Yang, Xiaofang Zhong, Dimitrios Gazgalis, Jesús Valdiviezo, Hyunil Jo, Yibing Wu, Morgan E. Diolaiti, Alan Ashworth, Nicholas F. Polizzi, William F. DeGrado

## Abstract

The de novo design of small-molecule-binding proteins has seen exciting recent progress; however, the ability to achieve exquisite affinity for binding small molecules while tuning specificity has not yet been demonstrated directly from computation. Here, we develop a computational procedure that results in the highest affinity binders to date with predetermined relative affinities, targeting a series of PARP1 inhibitors. Two of four designed proteins bound with affinities ranging from < 5 nM to low μM, in a predictable manner. X-ray crystal structures confirmed the accuracy of the designed protein-drug interactions. Molecular dynamics simulations informed the role of water in binding. Binding free-energy calculations performed directly on the designed models are in excellent agreement with the experimentally measured affinities, suggesting that the de novo design of small-molecule-binding proteins with tuned interaction energies is now feasible entirely from computation. We expect these methods to open many opportunities in biomedicine, including rapid sensor development, antidote design, and drug delivery vehicles.

**One Sentence Summary:** We use informatic sampling to design low nM drug-binding proteins, and physics-based calculations to accurately predict affinities.

## Main Text

Molecular recognition underlies the binding and catalysis of small molecules by protein receptors and enzymes (*1*–*5*). While we have an advanced understanding of both protein design and molecular interactions, the rational design of *de novo* proteins that specifically bind small molecules with low nM to pM affinity is a major challenge that has not been achieved in *de novo* proteins without experimental screening of large libraries of variants (*4*–*12*). The few successful examples of micromolar binders have relied on structural informatics to guide sampling of protein structure, sequence, and interactions, and scoring functions that rely on a mix of statistical and physical terms without explicit representation of dynamics, conformational entropy or water (*6*– *11*). From a practical perspective, the reliance on imperfect empirical and statistical information has limited the success of the process, and more fundamentally leaves open the question of whether our understanding is grounded in physical forces or limited exclusively to advanced pattern recognition (*13*–*17*). Here, we show how fully physics-based methods can be used to evaluate small molecule binders. We also introduce enhancements on existing sampling methods to better account for molecular complementarity between the ligand and protein. Finally, we show that all-atom molecular dynamics simulations are sufficient to describe the conformations, dynamics, binding interactions and free energies of association of a *de novo* protein designed to bind a series of poly(ADP-ribose) polymerase (PARP) inhibitors (PARPi) (*18*). These studies show the feasibility of combining informatics-based approaches for sampling and physics-based approaches for evaluation of top-scoring designs.

Inhibitors of PARP are a recently developed class of clinically useful anticancer drugs. De novo designed binders of PARPi drugs might serve as components in detectors, delivery agents, or detoxification agents for these cytotoxic drugs. The predominant class of PARPi drugs share a tripartite pharmacophore consisting of a fused 5,6-bicyclic core, an amide and a phenyl group bearing a positively charged alkylamine (Fig. 1A). We chose to target rucaparib, the most structurally complex of several related drugs, as our primary target (Fig.1A), as well as a series PARPi analogues. By considering a series of drugs, we at once provide reagents that might be widely useful, while simultaneously testing our understanding of the essential features required for binding.

**Figure 1.**
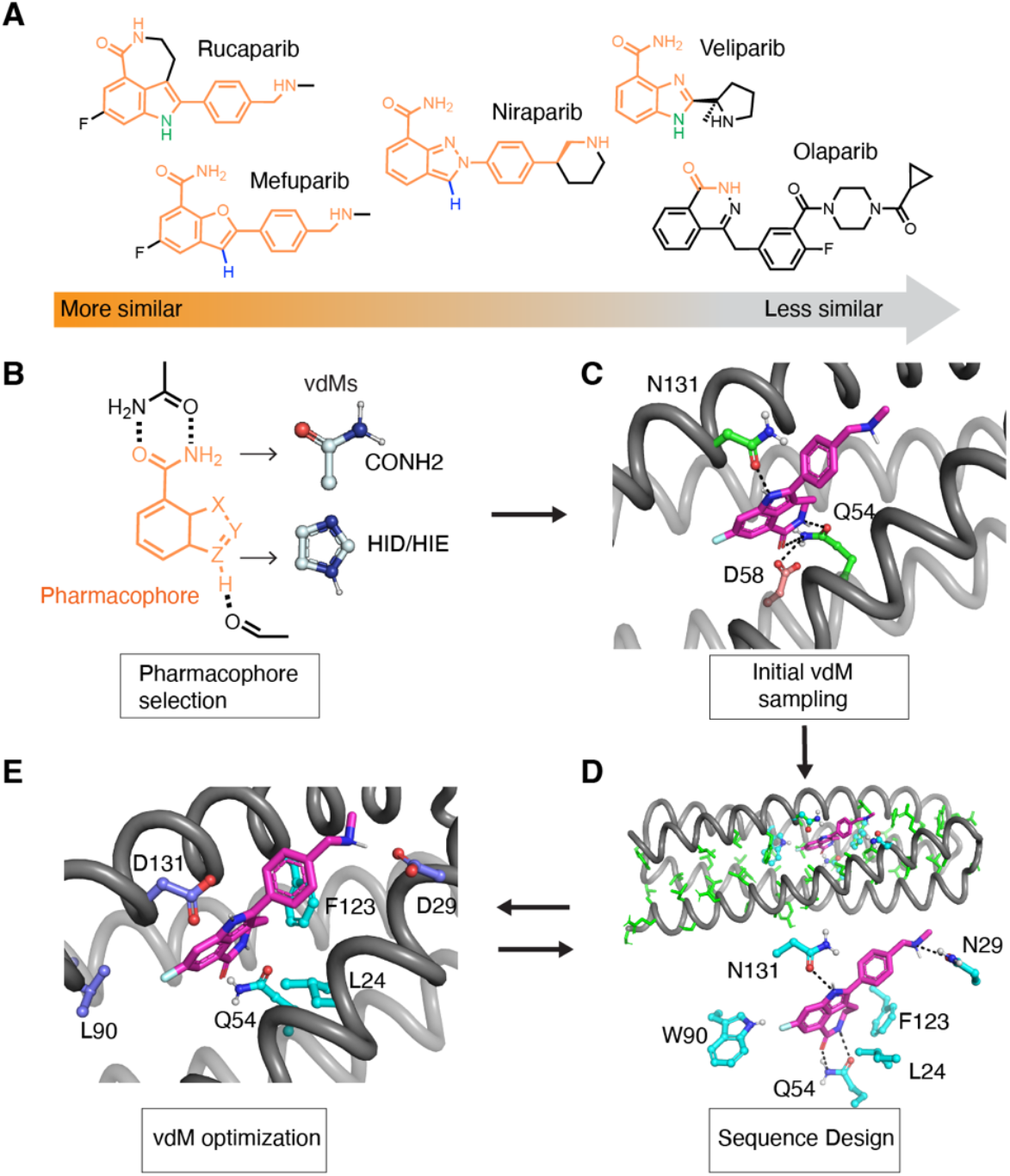
The computational design of poly(ADP-ribose) polymerase inhibitors (PARPi) protein binders. (A) The PARPi analogues. The shared chemical features are marked in orange. Olaparib is used as a negative control in the design and binding assay. (B – E) The overall design strategy. (B, C) We first define the pharmacophore and use COMBS to sample vdMs on the selected protein backbones. We initially targeted the indole and carboxamide of the drug, and used COMBS to discover sidechains that would form first and second-shell hydrogen bonds to both of these chemical groups. We discovered a solution in which the carboxamide formed bidentate hydrogen bonds with sidechain of Gln54, and the drug’s indole NH interacted with the Asn131 (C, carbon atoms of protein green, those of rucaparib are purple). A second-shell interaction to Q54 that was discovered by COMBS was Asp58 (carbons brown). (D) We applied flexible backbone sequence design with a custom Rosetta script while fixing the interactions selected from COMBS. (E) Then we search vdM again based on the design output from the previous sequence step. The slightly different (∼ 1 A Ca RMSD) backbone now preferred different vdMs at some locations (higher cluster scores) and these mutations were made. Three residues at 29, 90, 131 (deep blue) were changed based on COMBS results.

### *De novo* design of high-affinity drug-binding proteins

We used a new recursive version of the COMBS algorithm to design rucaparib-binding sites in a family of mathematically generated four-helix bundle proteins into which key binding residues were introduced using van der Mers (vdMs) (*10, 19, 20*). A vdM is an element of protein structure that identifies the preferred positions of interacting chemical groups relative to a residue’s backbone atoms (*10*). The COMBS algorithm finds positions on a given protein backbone that can simultaneously form favorable van der Waals, aromatic, and/or hydrogen-bonded interactions with the chemical groups of a target small molecule (Fig. 1B, 1C). To initiate the design process, we used COMBS to identify sidechains to bind rucaparib’s key polar groups, which included H-bonded interactions involving the indole NH, and the carboxamide’s C=O and NH_2_ groups (Fig. 1C). Additionally, COMBS identified Asp131 as a second-shell interaction to the carboxamide of rucaparib (Fig. 1C). It is important to design binding interactions with these groups with sub-Å accuracy to engender specificity and a favorable free energy of association. Next, the remainder of the sequence was designed using flexible backbone design (Fig. 1D) (*10, 21, 22*) while retaining the identity of the keystone residues (identified in the COMBS step). The mainchain moved 1 Å rmsd during this step (Fig. S1), so a second round of vdM sampling was performed on the relaxed backbone. This procedure identified three mutants involving drug-contacting residues, including N29D, W90L and N131D (Fig.1E). A second round of flexible backbone sequence design using this backbone and the newly fixed vdMs resulted in converged sequence/structure combinations (Fig. 1E, 2A-B), as a third round of COMBS showed the vdMs were now optimal. The final designs include numerous CH-π and hydrophobic interactions interspersed with specific polar interactions, including four H-bonds (an H-bond donor to the drug’s carboxamide oxygen as well as three H-bond acceptors to the drug’s carboxamide NH, indole NH and charged ammonium group), as well as second shell interactions. (Fig. 2C, Fig. S2).

**Figure 2.**
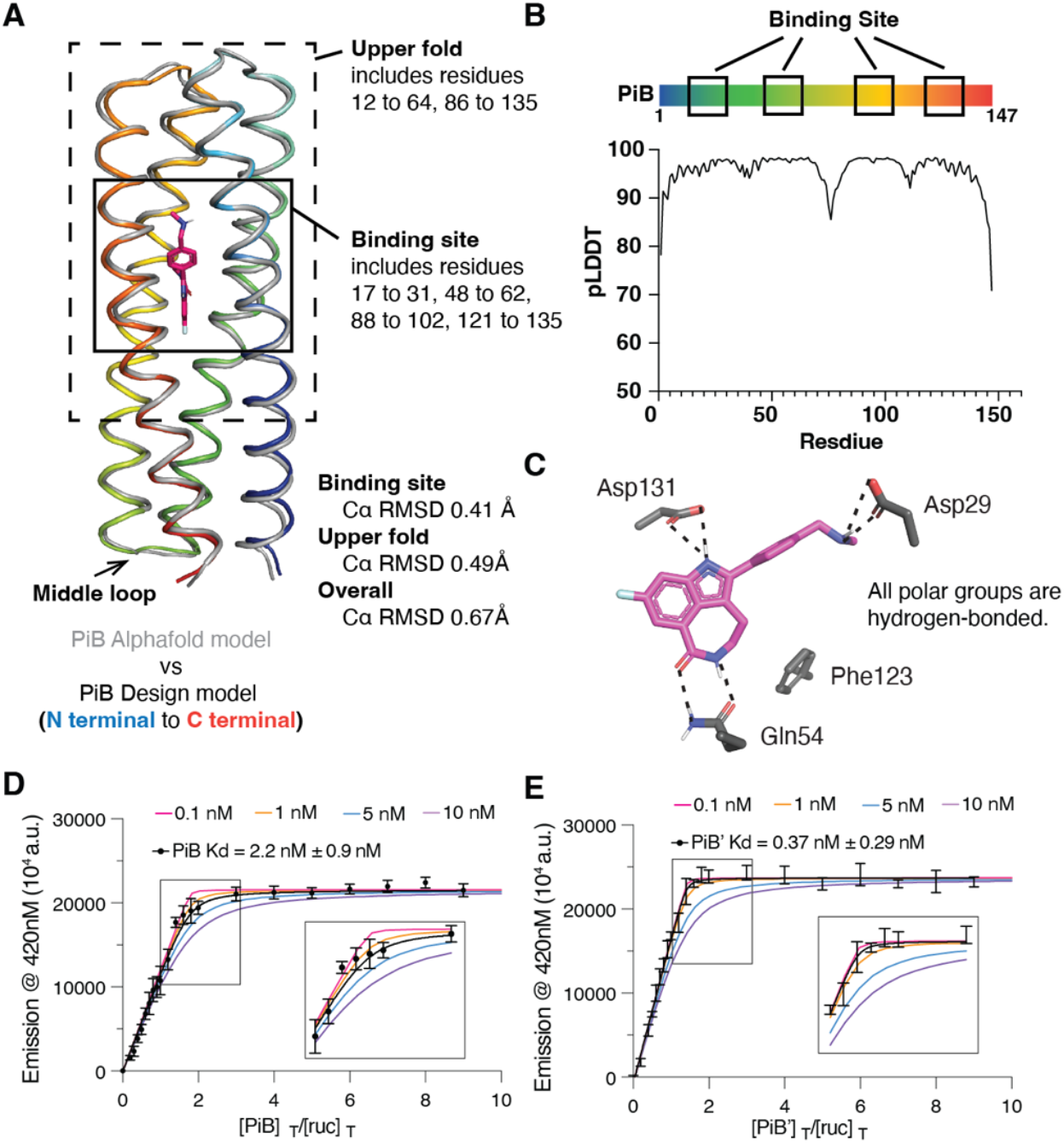
Assessing the computational model and experimental binding of PiB to rucaparib. (A) The AlphaFold2 model agrees with the designed PiB very well, with the binding site Cα RMSD of 0.41 Å, the upper fold Cα RMSD of 0.49 Å and overall Cα RMSD of 0.67 Å. (B) The predicted local distance difference test scores (pLDDTs) concur with the trend of RMSD difference of the design model. For example, the N-terminal, C-terminal and the middle loop with low pLDDTs (<90) showed higher Cα RMSD. (C) The design model showing the polar groups of rucaparib are all hydrogen-bonded. (D) (E) A fluorescence titration shows that PiB and PiB’ bind rucaparib with K_D_ < 5 nM. The fluorescence emission intensity at 420 nm of rucaparib (excitation wavelength 355 nm) was measured after titrating aliquots of PiB (D) or PiB’ (E) to a final concentration indicated in the abscissa. The data are well described by a single-site protein-ligand binding model, and a non-linear least squares fit to the data returned values of K_D_ of 2.2 (± 0.9) nM for PIB, and 0.37 (± 0.29) nM for PIB’. Although the fitting error was relatively small, a sensitivity analysis, in which the value of K_D_ was held constant at various values, showed that the data for both proteins were fit within experimental error so long as the K_D_ is less than 5 nM. Therefore, while the most probable binding constants were 2 and 0.4 nM, respectively, we can confidently conclude that the values for PiB and PiB’ are less than 5 nM. The titration was carried out in in buffer containing 50 mM Tris, 100 mM NaCl (pH 7.4).

Throughout the design process, we ensured that the designs would also retain favorable interactions with most of the common pharmacophores of the three other drugs (see supplementary methods for details). However, we predicted that the protein would have lower affinity for niraparib and mefuparib, because they lack the H-bonding group indole NH of rucaparib. Also, we expected veliparib to bind weakly, because it lacks a hydrophobic phenyl group and the position of its charged ammonium group differs significantly from that found in the other three drugs.

The final models were chosen based on multiple criteria: 1) low Rosetta energy (lowest 50 of the 1000 total designs); 2) favorable vdMs (highest total vdM cluster scores); 3) satisfaction of all buried H-bond donors in the protein and ligand; and 4) avoidance of clashes with the three other PARP inhibitors, which show significant structural variability near the amine end of the molecule. Four designs were selected for expression: two designated as PiB (PARPi binder) and PiB’, respectively, are closely related (Table. S1, Fig. S3). PiB’ differs from PiB only by the substitution of five solvent-exposed charged residues with Ala to enhance the crystallinity of the protein for X-ray crystallography. The other two (PiB-1 and PiB-2) were less closely related to PiB in structure and sequence (Fig. S4). Circular dichroism spectroscopy showed all four had substantial alpha-helical character (Fig. S5). However, PiB-1 and PiB-2 failed to induce large changes in the fluorescence emission spectrum of rucaparib (Fig. S6). Therefore, we focused our efforts on PiB and PiB’ (Fig. S7 - S11).

Spectral titrations showed that PiB and PiB’ bound the PARPi drugs with high affinity. Incubation of PiB with equimolar concentrations of rucaparib led to a marked blue shift and an increase in intensity of its fluorescence spectrum, as expected if its indole core were bound in a rigid, solvent-inaccessible site (Fig. S6, S7). NMR spectroscopy of PiB showed that it folded into a well-defined structure, and the addition of a single equivalent of rucaparib led to a new set of peaks, consistent with a stoichiometric, specific complex (Fig. S10, S11). Fluorescently monitored titrations of protein into a solution of rucaparib showed that PiB and PiB’ bound with very low to sub-nM affinity (Fig. 2D, 2E, 3A). Even at the lowest experimentally feasible rucaparib concentration, the binding isotherms show a linear increase in intensity with respect to protein concentration until a single equivalent is added, followed by an abrupt leveling at higher protein concentrations. This behavior is indicative of a dissociation constant that is much lower than the total rucaparib concentration. A non-linear least-squares fit to the data returned a K_D_ of 2.2 nM for PiB and 0.37 nM For PiB’, and a sensitivity analysis showed that the K_D_ was less than 5 nM for both proteins (Fig. 2D, 2E) (*10*). Thus, PIB and PiB’ are the first reported *de novo* protein that bind a small molecule with very low, single digit nM to pM binding affinity prior to extensive experimental optimization.

UV/visible absorption titrations showed that PiB and PiB’ also bound to the remaining ligands with affinities that grew increasingly weaker as the drugs’ structures diverged from rucaparib (Fig. 3A, Fig. S12). PiB retained sub-μM affinity for mefuparib (K_D_ = 190 nM and 350 nM for PiB and PiB’, respectively) and niraparib (600 and 550 nM). The corresponding K_D_ values were 14 μM and 24 μM, respectively for the structurally divergent drug veliparib, and no binding was detected for the most divergent drug, Olaparib (Fig. S13). This observed trend in binding affinity matches our hypothesized order.

**Figure 3.**
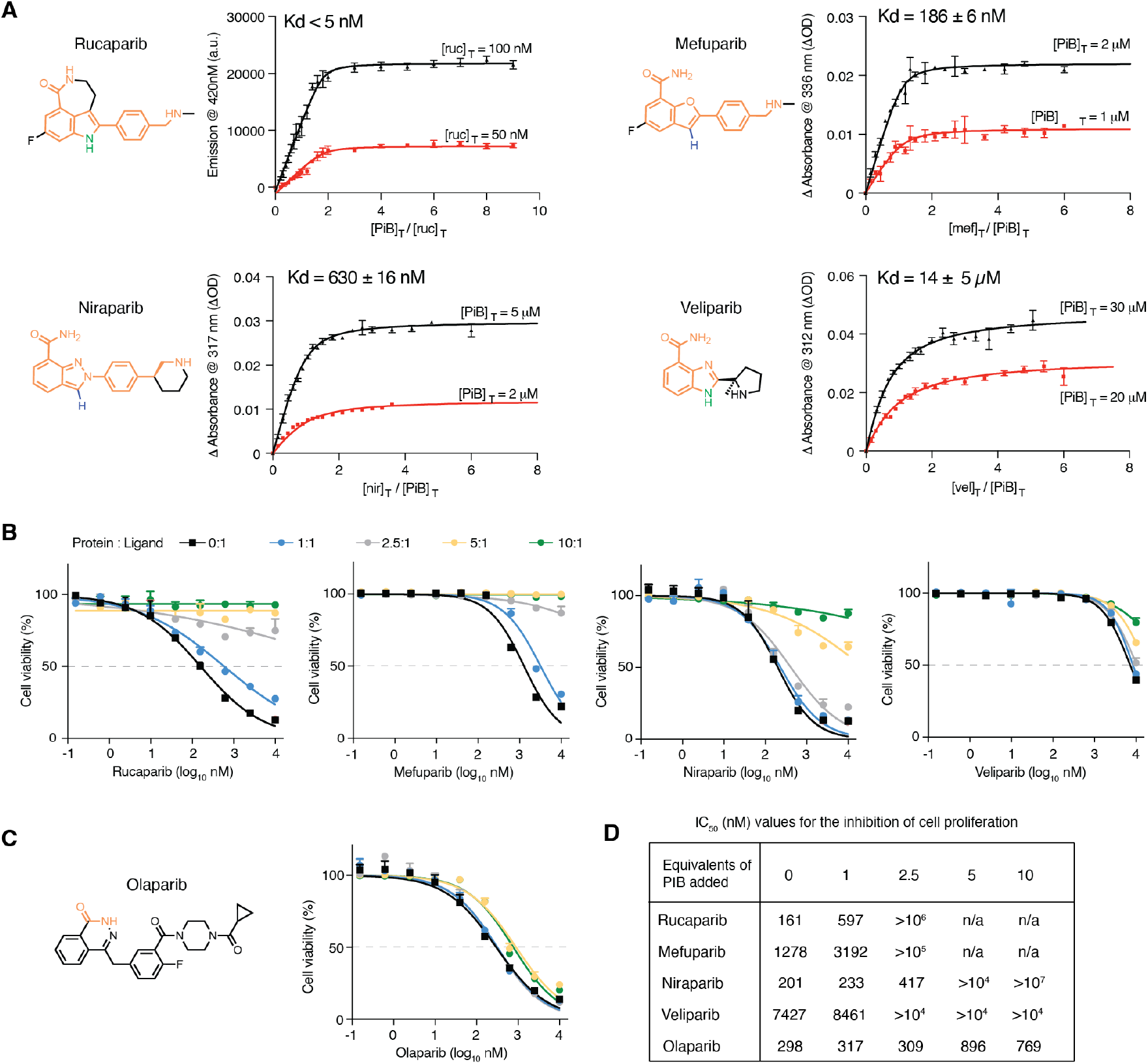
Spectral titrations and cell viability assay of PiB with PARPi. (A) The values of K_D_ of various drugs for PiB as obtained from global fit of a single-site binding model to the fluorescence changes (A, from Fig. 3) or absorbance changes as a function of the concentration of PiB. Indicated wavelengths for the titration were chosen to maximize the difference in absorption for the free versus bound drug. (B) Seven-day growth assays in DLD-1 *BRCA2* mutated cells show that PiB alleviates the effects of rucaparib, mefuparib, niraparib and veliparib toxicity in a dose-dependent manner. The PARP inhibitors were pre-incubated with PiB in media at room temperature for 5 minutes at multiple concentration ratios (ligand : protein) of 1:0, 1:0.2, 1:1, 1:2.5, 1:5 and 1:10. (C) Cell viability assay as in Figure 4B showing that PiB had no effect on the olaparib dose response. (D) Table showing IC_50_ values for the inhibition of cell proliferation by PARPi drugs in the presence of increasing mole ratios of added PiB protein.

We next examined the *in vitro* stability and potency of PiB and PiB’ in serum and cellular assays. PiB and PiB’ were highly stable in human serum, as are other *de novo* proteins designed for medical applications (Fig. S14, S15) (*10, 23, 24*). PARP inhibitors potently inhibit the viability of cells with certain DNA repair deficiencies, including loss-of-function mutations in *BRCA2*. To determine whether PiB and PiB’ could attenuate the lethal effects of PARPi drugs, we measured their effects on the growth of *BRCA2* mutated DLD-1 cells and SUM149 cells (*25*) over after an 8-day incubation. Dose-response curves were first established in the absence of PiB, then the titration was repeated with PiB or PiB’, at varying [protein]/[drug] ratios for each PARPi drug concentration. Addition of a single equivalent of PiB or PiB’ resulted in a 4-fold increase in the half maximal inhibitory concentration (IC)_50_ value for rucaparib. Thus, PiB competes effectively for binding of rucaparib to human PARP1, an enzyme reported to bind rucaparib with a dissociation constant of 0.1 to 1 nM in biochemical assays (*26, 27*) (Fig. 3A, 3B, 3D, Fig S16, S17, S18). The potency of PiB and PiB’ in the cell viability assay generally tracks with the spectroscopic assays, with the protein showing effects on mefuparib and niraparib intermediate between that for rucaparib and veliparib (*26*) (Fig. 3B, 3D, Fig. S16, S17, S18). Moreover, PiB and PiB’ did not significantly change the cellular response to olaparib (Fig. 3C, 3D, Fig. S16, S17, S18) in line with spectroscopic data that indicated that PiB and PiB’ does not bind this drug.

### High-resolution crystal structures of complexes are in remarkable agreement with the designs

The crystallographic structures of PiB’ were solved in the absence and presence of the four active compounds at 1.3-1.6 Å resolution (Table S2). The protein’s conformation is in excellent agreement with the predicted AlphaFold2 model (Table. S3), particularly near the binding site (Cα RMSD of the 60 surrounding residues was 0.2-0.5 Å, Fig. 2A, 4A-B, S19, Table. S3). The binding pocket of PiB’ is very well pre-organized, and the binding of the drugs leads to only 0.2 – 0.5 Å Cα RMSD changes in the binding-site residues (Fig. 4A-4C, S20, Table. S3). The conformations of the sidechains in the unbound structure are almost entirely as in the design, and they interact precisely as predicted in the design of the rucaparib complex (Fig. 4B, S20): Asp29 makes a direct H-bonded salt bridge to the drug’s charged ammonium group. Rucaparib’s carboxamide forms a two-coordinate hydrogen bond with Gln54, which in turn is stabilized by a second-shell network of H-bonds predicted in the design; Asp131 formed a solvent mediated H-bond to rucaparib’s indole NH group (Fig. 4B, S20). Interestingly, a search of water-mediated Asp sidechains with related indole and imidazole sidechains showed this bridging interaction is frequently found in the PDB (Fig. S21).

**Figure 4.**
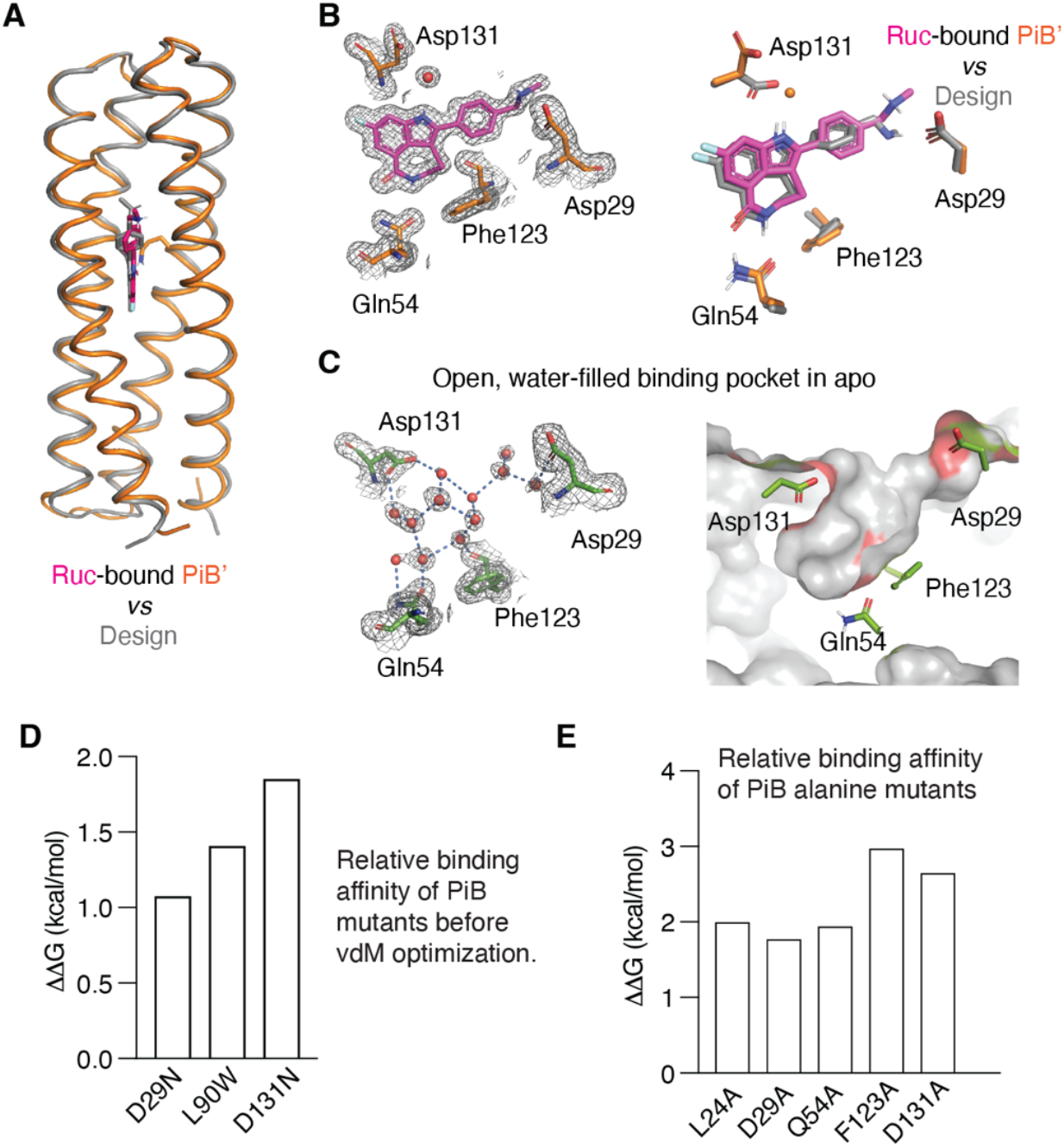
The structure of drug-bound PiB’ agrees with the design. (A) The design model agrees well with the rucaparib-bound PiB’ crystal structure, with binding site (Fig. 3A) Cα RMSD range between 0.38-0.46 Å for the three monomers in the asymmetric unit. (B) The binding site of PiB’. A 2mFo-DFc composite omit map contoured at 1.6 σ. The map was generated from a model that omitted coordinates of rucaparib. Overlay of the design (gray) and the structure (protein in orange, rucaparib in pink). The sidechains of the binding pocket in rucaparib-bound PiB’ agrees with the design. Asp131 interacts with the indole NH via a bridging water as in MD simulations. (C) The structure of apo-PiB’ shows a preorganized open pocket filled with multiple waters, which are displaced in holo structure. (D) Reversal of the three designed substitutions from the vdM optimization procedure led to lower binding affinity (higher K_D_) for rucaparib by fluorescence emission titrations. (E) Alanine mutations of the direct binding residues decreased binding affinities confirmed by fluorescence emission titrations.

The structures of mefuparib and niraprib bound to PiB’ show a similar set of interactions as rucaparib (Fig. S22). However, their aromatic 5-membered azole ring lacks a H-bonding group to interact with Asp131, explaining their decreased affinity for the protein. As expected from its divergent structure, veliparib has a less favorable fit with PiB’s binding site, and it lacks a salt bridge to its ammonium group as in other complexes (Fig. S22, S23). In summary, the structures are in excellent agreement with the design and confirm our structural prediction of the affinity differences between drugs.

Three residues were changed to improve binding during the second round of COMBS design of PiB. To determine whether these substitutions indeed increased affinity, we evaluated mutants with the second-round substitutions reverted to their identities in the first round of design. These changes each led to one to two orders of magnitude weaker binding affinity for rucaparib Asp29Asn (K_D_ = 13 nM), Leu90Trp (K_D_ = 24 nM) and Asp131Asn (K_D_ = 50 nM) (Fig. 4D, S24A). Thus, iterative vdM selection successfully identified interactions with improved binding affinity. We suggest that vdM guided amino acid optimization might provide a useful alternative to other methods of affinity optimization. We also conducted an alanine scan to probe the energetic contribution of each of the residues that lined the pocket in the rucaparib complex (Fig. 4E, S24B). Each mutation was unfavorable with values of ΔΔG ranging from 1.7 to 3.0 kcal/mol. These values are within the range observed for substitutions of critical binding residues in natural protein binding sites (*28*).

### Molecular dynamics and free energy calculations confirm mode of binding and accurately predict binding thermodynamics

We performed 2.0-microsecond all-atom molecular dynamics (MD) simulations to compare the structural stability of PiB, PiB’, PiB1, and PiB2 in complex with rucaparib. The simulations were performed on the designed models (instead of the crystallographic structures) to assess the usefulness of incorporating MD into a design pipeline. The protein backbone conformations were very stable for all three complexes. However, rucaparib’s designed binding pose was stable only in PiB and PiB’ (Fig. S23) (as PiB and PiB’ behave similar in MD, we only use PiB to illustrate later): it retained its bivalent hydrogen bonding interaction to Gln54 (Fig 5A), and Asp29 and Asp131 showed stable interactions with rucaparib’s indole NH and ammonium groups through direct and water-mediated hydrogen bonds, respectively (Fig. 5A). By contrast, PiB-1 and PiB-2 simulations exhibited significant deviation from rucaparib’s designed pose, and their key buried H-bonds to Gln54 were broken within 50 nanoseconds in each of three independent calculations (Fig. S25). Moreover, PiB shielded the apolar atoms in rucaparib more efficiently in PiB than PiB1 and PiB2, as determined from solvent-accessible surface area calculations (*29*) within individual MD trajectories (Fig. S26). Furthermore, MD simulations of PiB in complex with niraparib, veliparib, and mefuparib show similar binding-site stability as PiB : rucaparib over 2.0 microseconds (Fig. 5A, S27). Thus, MD appears to be a useful tool in assessing the stability of interactions in designed complexes.

**Figure 5.**
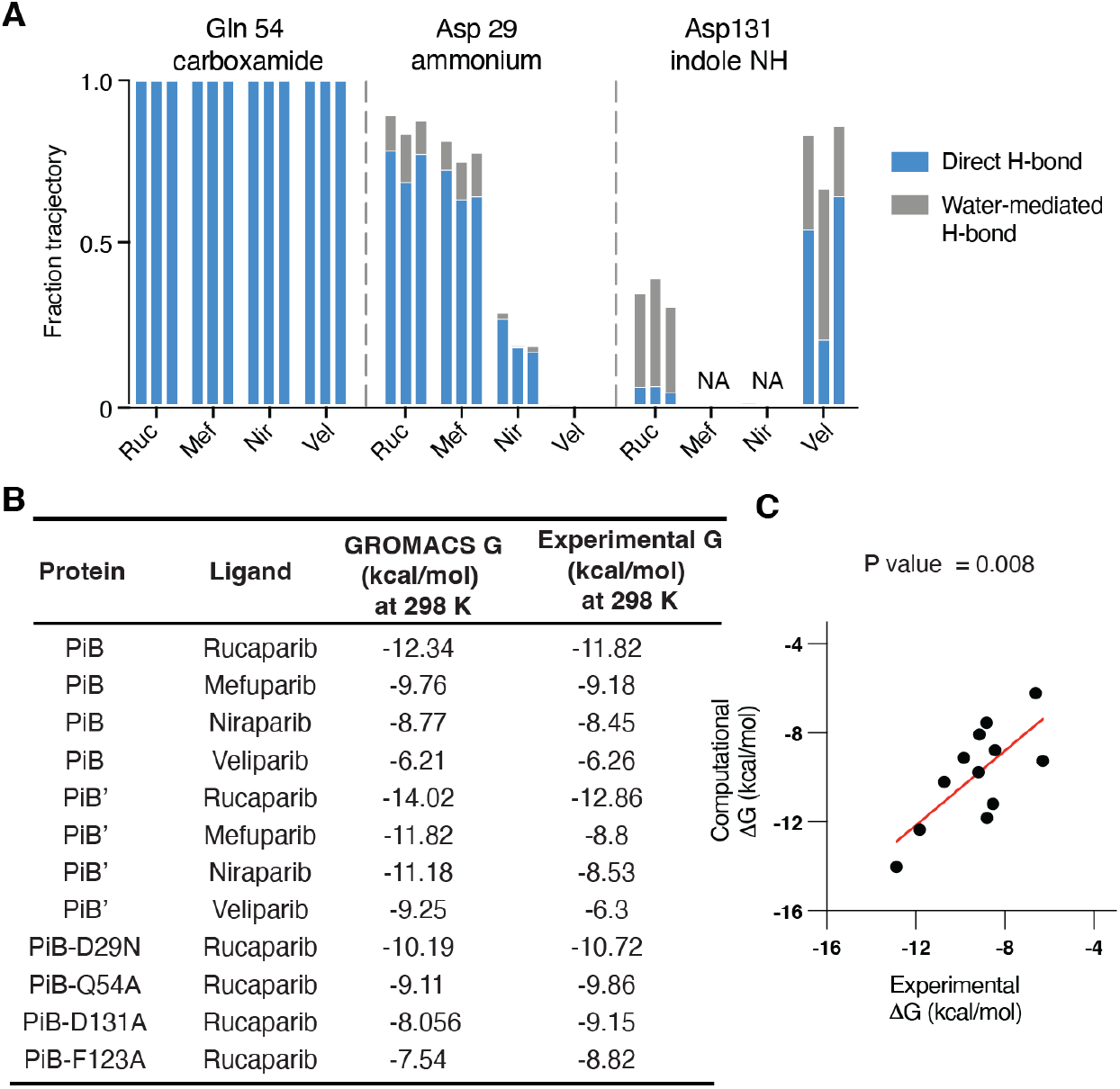
The MD simulations of PiB, PiB’, and mutants. (A) Using unbiased molecular dynamics simulations in Amber, we calculated (in triplicate) the frequency with which the intermolecular hydrogen bonds formed between the protein scaffold and the bound drug molecule. PiB was found to form a hydrogen bond between Gln54 and the targeted drug carboxamide in 100% of all simulations for each drug complex. The charged ammonium groups of rucaparib and mefuparib interacted with Asp29 through a combination of direct and water-mediated hydrogen bonds, totaling to more than half of the full simulation time, which contrasts niraparib and veliparib’s inabilities to form equivalent hydrogen bonds (due to changes in chemical structure around the ammonium tail of the ligand). In a small fraction of each rucaparib and veliparib trajectory, Asp131 engaged in water-mediated hydrogen bonds to the drugs. (B) Using biased simulations in GROMACS, we calculated binding free energies for each ligand and found that ranked affinity for each drug is consistent with experimental results. (C) Comparison of ΔG binding from the GROMACS calculation with the experimental value from spectral titrations.

We next turned to alchemical and physical-pathway methods to calculate absolute binding free energies to determine whether the dissociation constants for binding of the drugs to PiB and PiB’ might be predicted directly from molecular dynamics simulations. The alchemical transfer method (*29*–*31*) was carried out starting with the design models by two of the authors, who had no knowledge of the experimental results. This method has been shown to be comparable to other alchemical methodologies such as Schrodinger’s FEP+ (*30*) or Amber’s thermodynamic integration (*31*) given comparable sampling of the configurational space. An initial absolute binding free energy calculation was used to evaluate the energetic contributions of the fused-ring cores of the drugs and to ensure convergence of the calculations. An additional relative binding free energy calculation was performed to transform each core into the target ligand to estimate the contribution from non-core regions (Fig. S28). Universally, the alchemical transfer method tended to overestimate the binding energy, possibly due to having two sets of restraint potentials. However, this procedure correctly predicted the relative affinities of the four ligands (Table. S4).

We next used potential of mean force calculations, an orthogonal physical-pathway methodology (*32*), to compute absolute binding energies, and found that the results were in remarkably good agreement with experiment (Fig. S29). The RMS error between the predicted and experimental values is 1.3 kcal/mol, and the correct rank order of affinities was observed. This error is close to the experimental error in the measurement of KD for rucaparib (Fig. 5B, 5C, Fig. S30-S33, Table. S5). We also obtained very good agreement between computation and experiment for a set of four mutants of PiB (Fig. 5B, 5C, Table. S5). This agreement bodes well for the use of alchemical and physical-pathway-based binding free energy calculations to evaluate potential binding energies of de novo small-molecule binding proteins.

## Conclusion

The *de novo* design of proteins with very high affinity for small-molecule ligands has been a challenging endeavor, because the design process generally relies on sampling and scoring functions that are optimized for predicting protein structure rather than binding to the diversity of functionality encountered in small molecules. In this work we enhanced the COMBS method, which uses van der Mers to sample realistic binding poses and key interactions within a protein framework. In this procedure, vdMs are used to identify binding-site residues capable of forming key interactions such as hydrogen bonds and aromatic interactions to enable binding of a small molecules, in much the same way that natural proteins bind a wide diversity of small molecules using a set of 20 amino acids. While the energetics of these interactions can vary depending on the specific small molecule bound, the fundamental geometries required to achieve optimal binding remain relatively constant. Thus, a common set of vdMs should serve to bind a wide range of compounds.

A number of other methods for designing ligand-binding proteins are appearing in the recent literature (*12*). However, at the end of any design procedure, the designer is presented with a list of candidates for further consideration. With sufficient resources it is feasible to synthesize genes encoding thousands of candidates and evaluate them individually using robotics or yeast display to identify a few binders. Additionally, because so many variants are screened, the characterization of the binders has relied on organic synthesis of fluorescently labeled or biotinylated versions of the ligand, again increasing the barriers to widespread economical use of computational design.

Here we identify a series of physically grounded steps to help identify the tightest binders in a computational library of potential proteins (Fig. S34). 1) We prioritized the principle of preorganization: AlphaFold2 predicted a preorganized site, and we find greater discrimination using all-atom MD simulations. Even in the absence of the drug, the conformation of the backbone and binding-site residues of PiB were stable on the nanosecond/microsecond timescale. While we conducted relatively long 2 μsec simulations, we found that 100 nsec simulations were sufficient to identify the successful binder. 2) We focus on designs that maximize burial of apolar atoms based on calculation of solvent-accessible surface area in the bound versus free state. 3) Importantly, the H-bond potential of buried polar atoms should be maximized for both the bound drug as well as the interacting sidechains, as the loss of water interactions have to be recouped by drug–protein H-bonding, a highly unfavorable process (*33*). This is apparent if we compare PiB to ABLE, the first COMBS-designed drug binder (*10*). The PiB:rucaparib (app. 1 nM) binding site maximizes the H-bonding potential of both the drug and the first-shell liganding sidechains, whereas the ABLE:apixaban (5 μM) binding site has four unsatisfied hydrogen-bonding sites (Fig. S35). Water-bridging H-bonds can also be identified by vdMs (that include explicit bridging waters) and MD simulations. 4) We placed the charged ammonium group of rucaparib and interacting sidechains at a surface location that minimizes the Born solvation energy (*34*), simplifying the task of recognizing charged groups.

Finally, while MD simulations of *de novo* proteins (*14, 34*–*36*) can help screen designs (*14, 34*–*36*), free energy calculations have not previously been applied to designed proteins. Although they are more computationally intensive and require more user-specified parameters, we obtained excellent quantitative agreement between computed and experimentally measured binding free energies using the designed models as the starting structures. These data demonstrate for the first time the possibility of designing proteins with high affinity (< 5 nM) to small molecules using fully rational criteria for design and “physics-based” force fields to evaluate the complex.

It is interesting to compare the mode of interaction and interaction energy of rucaparib for PiB versus the natural protein PARP1. Although we designed PiB without considering the natural binding-site interactions in RARP1, COMBS identified a very similar set of interactions to bind rucaparib (Fig. S36): a ligand-Asp salt bridge, solvent-mediated hydrogen bond to rucparib’s indole NH, and a two-coordinate H-bond to the drug’s amide. This commonality likely reflects the fact that proteins have only a limited repertoire of functional groups, and COMBS is capable of identifying highly favorable interactions, similar to those used in nature.

Rucaparib binds to the human PARP1 enzyme with a K_D_ ranging from 0.1 to 1.5 nM, depending on the experimental conditions, within the range observed for PiB and PiB’(*26, 27*). Rucaparib is a third-generation drug that was discovered only after synthesizing hundreds of analogues in multiple groups. Thus, it is noteworthy that we were able to design a protein with similar affinity to rucaparib as PARP1 after screening only a few proteins. Ligand efficiency is often used as a guiding rule in drug discovery to determine whether the affinity of a molecule of a given size is within a range typically seen in highly optimized developed small-molecule drugs and natural organic ligands for proteins (*37, 38*). As ligands become larger, they have more opportunities to form favorable interactions with their target proteins. Thus, the maximal affinity possible roughly scales with the size of a small molecule, and the ligand efficiency is defined as the free energy of binding (1 M standard state) divided by the number of heavy atoms in the ligand. Most drugs have ligand efficiency around 0.3 kcal/(mol * heavy atom count) (*37, 38*), although higher values are observed for highly optimized drugs such as rucaparib, which has a ligand efficiency of 0.5 kcal/(mol*heavy atom count). The ligand efficiency of a drug is similarly a good measure of how well optimized a *de novo* protein is for binding to a small molecule. The 0.5 kcal/(mol * heavy atom count) ligand efficiency of PiB is an considerable improvement over the 0.21 to 0.26 ligand efficiency of the first COMBS-designed apixaban binders, demonstrating the significance of incorporating the design principles discussed.

In nature, small molecule-binding proteins serve a range of functions, and we might expect a similarly broad spectrum of practical applications of designed binders for sensing and pharmaceutical applications. For example, de novo proteins might be used as antidotes to neutralize drugs that have accumulated to toxic levels, or to reverse the action of anti-coagulants prior to surgery to reverse the risk of bleeding. In other cases, proteins could be used as drug carriers to tune their pharmacokinetic properties. Drugs might also be released in the vicinity of tumors or sites of infection by introducing proteolytic sites into flexible loops that allow cleavage by proteases enriched in these environments (*39*). Furthermore, de novo proteins might be targeted to tumor environments by fusing sequences (e.g., disulfide-linked cyclic RGD peptides (*40*) or peptide hormones such as somatostatin (*41*) to bind to proteins enriched on the surface of cancer cells. Finally, designed proteins can be used to sequester environmental toxins or as components in detection devices.

## Acknowledgments

We thank members of the DeGrado lab for discussion. Beamline 8.3.1 of the Advanced Light Source, a DOE Office of Science User Facility under Contract No. DE-AC02-05CH11231, is supported in part by the ALS-ENABLE program funded by the National Institutes of Health, National Institute of General Medical Sciences, grant P30 GM124169-01. We are grateful to George Meigs from LBNL helping with the setup.

## Author Contributions

L.L. and W.F.D conceived the idea for the project, L.L. and N.P. developed computer code, and L.L. performed the computational design. L.L., X.G., S.I.M., X.Z., H.J. performed the experiments. L.L., H.Y., X.G., Y.W. performed the data analysis. N.P., S.T., D.G., J.V. ran and analyzed MD stimulations. X.G., M.D., A.A., and W.F.D. designed the cell assay experiments. L.L., N.P., and W.F.D. wrote the paper with input from all authors.

## Data and materials availability

Coordinates and structure files have been deposited to the Protein Data Bank (PDB) with accession codes: 8TN1 (apo-PiB), 8TN6 (rucaparib-bound PiB), 8TNB (mefuparib-bound PiB), 8TNC (niraparib-bound PiB), 8TND (veliparib-bound PiB). Computational code and design scripts are available in the supplementary materials and at GitHub. All other relevant data are available in the main text or the supplementary materials.

## Supplementary Materials

Materials and Methods

Figs. S1 to S36

Tables S1 to S5

Supplementary Text

References

