## Supplementary file for "*De novo* design of drug-binding proteins with predictable binding energy and specificity"

Science
AAAS

### Supplementary Materials for

#### Computational *de novo* design and energetic analysis of high-affinity drug-binding proteins

Lei Lu[1], Xuxu Gou[2], Sophia K Tan[1], Samuel I. Mann[1], Hyunjun Yang[1], Xiaofang Zhong[3], Dimitrios Gazgalis[4], Jesús Valdiviezo[4], Hyunil Jo[1], Yibing Wu[1], Morgan E. Diolaiti[2], Alan Ashworth[2], Nicholas F. Polizzi[4], William F. DeGrado[1]*

**The PDF file includes:**

Materials and Methods
Supplementary Text
Figs. S1 to S36
Tables S1 to S5
References

#### Materials and Methods

Overview of Design process

We used a library of the previously generated parameterized four-helix bundles from CCCP program as a starting point to design the PAPRi binders (available in the supplementary zip file) (*10, 20*). These four-helix bundles are highly designable scaffolds that can accommodate many sequences and contain a range of binding cavities for ligand binding as the bundles used the Crick parameters to mimic the natural or designed porphyrin-binding proteins. We then extract the shared chemical groups from the PARP inhibitors (**Fig. 1**). The drugs contain three polar chemical groups, consisting of indole NH and carboxamide that are common to all four PARP inhibitors, and a structurally more variable basic amine. The conformations of the drugs were computed using molecular mechanics (Maestro in Schrodinger software), and the lowest energy structure used for design. Given the variability of the "amine" portion of the drugs, we first concentrated on finding vdMs for the indole (Hid and Hie vdMs were used to approximate the pyrrole group of the indole) and carboxamide (conh2 vdMs from Asn and Gln) of rucaparib. Only inward directed residues were considered, and these were chosen based on the alpha-hull algorithm in COMBS, which defines the surface of a protein from only its backbone coordinates. Two backbones gave satisfactory solutions, with good overlap between the vdM chemical groups and those of the ligand. Only poses that showed no steric overlaps between any portion of the drug molecules and the protein backbone were considered. Poses were ranked based on their vdM score (*10*) loops were introduced between the helices using python scripts based on MASTER (*42*). The output backbone, including the selected vdMs and the pose of rucaparib, was submitted to variable backbone sequence design using a previously published script (provided in the supplementary text, page 48).

We selected the 50 best-scoring outputs (lowest Rosetta energy units, using the Rosetta Ref2015 energy function) from 1000 Rosetta flexible-backbone sequence designs. We next ran structure prediction using AlphaFold2 (*43, 44*) and OmegaFold (*45*) to eliminate sequences that were unlikely to adopt the desired fold. Sequences that gave an RMSD difference between the design model and prediction of less than 1.0 Å and a plDDT score >0.9 were further examined to assure that the design models satisfied the following criteria. 1) All H-bonds to all polar groups of the ligand were satisfied; 2) The heterocyclic core of viraparib, niraparib and mefuparib were superimposed onto pose of rucaparib in the designed protein models, and any design in which any of the three drugs showed a steric clash was eliminated. This step helped assure that the PiB protein would bind all four drugs considered. 3) The amine group of the drugs should be placed near the protein surface to minimize the Born energy of burial. This procedure resulted in three initial designs.

For design of PiB, we next examined the vdM scores of each residue in direct contact with rucaparib, including residues that were introduced by the Rosetta sequence design algorithm. vdMs were exhaustively evaluated for each chemical group present in rucaparib (e.g., Asp($CONH_2$) would be probed in place of Asn($CONH_2$)) and the vdMs with the better (higher) cluster score (*C* score) were chosen so long as they still satisfied the full hydrogen-bonding requirements. When more than one vdM had similar scores at a given position, the one with the greatest number of hydrogen bonds was chosen (e.g., bindentate would be favored over monovalent H-bonds).

At this step, the benzylic methyl amino group of rucaparib presented a special challenge, because it is a secondary amine (with a single NH group) that is not present in any protein sidechain. However, the intent of using vdMs is simply to sample positions where a given amino acid can form a hydrogen bond to the targeted chemical group, so we reasoned that any secondary amine or amide of similar steric encumbrance would be able to recruit an appropriate hydrogen-bond donating sidechain. A backbone Gly residue presents a disubstituted NH group, which was sterically similar to the secondary amine of rucaparib. Thus, we chose this to search for Gly NH as the chemical group (a special class of the more generic backbone chemical group). Even though the precise electronic structure differs from that of a secondary amine, it still recruited Asp29, because of Asp's ability to form a strong hydrogen bond with an NH group. This finding shows the versatility of van der Mers to identify favorable interactions, even when the only commonalities between the target chemical group and those in our library relate only to sterics and hydrogen-bonding potential. This process resulted in a total of three substitutions (N29D, W90L, N131D). Note that both hydrogen-bonding as well as apolar vdMs were considered, resulting in the Trp to Leu substitution. Finally, the sequence was re-designed using the same Rosetta flexible backbone script giving the structures from the last round of Rosetta design as shown in Fig.1. (The starting structures, the files to run initial vdM sampling, and model coordinates are provided in supplemental data as zip files).

This process resulted in three final models, PiB, PiB1 and PiB2. Visual inspection suggested that PiB2 bound in a somewhat strained conformation with non-linear hydrogen bonds and poor local packing, while PiB and PiB1 appeared to have more canonical H-bond geometries with the ligand (only MD was able to differentiate these two, see below). As a test of the design process, all three sequences were selected for experimental characterization.

vdM sampling

We used COMBS (https://github.com/npolizzi/Combs2) and extra python scripts in Rvrsprot (https://github.com/lonelu/Rvrsprot) to sample ligand-protein interactions (*10*).

Loop construction

Loops were selected using python scripts in Rvrsprot (https://github.com/npolizzi/Combs2). Rvrsprot called MASTER (*36*) (Method of Accelerated Search for Tertiary Ensemble Representatives) to query for loops using two segments (7 residues from each helix) of adjacent helices with specified length connecting them within a given RMSD. The program then clustered the loops and generated sequence logo information for the purpose of residue selection.

Flexible backbone sequence design

We used the program Rosetta (version linux_2020.08.61146) to perform flexible-backbone sequence design with a custom protocol (provided in the supplementary text, page 48). Simply, the protocol proceeds through a cycle of backbone relaxation, side-chain relaxation, and fixed backbone design with filtering based on ligand constraints and core packing.

Structure prediction

OmegaFold was installed locally for rapid prediction of low-scoring Rosetta designs (*45*). The Colab version of AlphaFold2 (https://github.com/sokrypton/ColabFold) was used to further predict the selected designed sequences (*44*).

Protein expression

The genes for the designed proteins (including an N-terminal TEV protease cleavage side followed by 6xHis-tag) were codon optimized. The genes were inserted in pET28a plasmids between Nde1 and Xho1 and were ordered from IDT or GenScript. Note that the residue numbering does not contain the N-terminal 6*His tag and TEV protease cleavage site.
The plasmids were transformed into *E.coli* BL21(DE3) cells and grown in LB media with kanamycin at 37 ˚C. For NMR samples, the cells were grown in M9 minimal media with C13-labeled glucose and N-15 labeled ammonia from Cambridge Isotopes. The cells were induced with IPTG when OD at 600 nm reached 0.8 and were pelleted by centrifugation after 4 hours and frozen at -80 ˚C. Cells were resuspended in TBS buffer (50 mM Tris, 100 mM NaCl, pH=7.4) and lysed by sonication. The proteins were purified by Ni NTA affinity column (Invitrogen) and their molecular weight was confirmed by gel electrophoresis. The proteins were desalted by buffer exchange and their His-tag was cleaved with His-tag TEV protease by overnight incubation at room temperature. The cleaved proteins were collected from the flow-through of Ni-NTA affinity column and concentrated for FPLC purification with Superdex 75 Increase 10/300 column. Both TEV-cleaved and His-tagged proteins were used in binding assays as they showed no significant difference in binding. In the other experiments, we used TEV-cleaved proteins.

Construction of mutants

Single amino-acid mutations of PiB plasmid were made via the Q5 Site-Directed Mutagenesis Kit (NEB) following the provided protocol of the kit and primers were designed using the QuickChange primer design program and ordered from IDT. The mutants were confirmed by Sanger sequencing (GeneWiz) of single colonies with the transformed genes.

Size exclusion chromatography

We obtained gel filtration profiles using a Superdex 75 Increase 10/300 column on a GE Healthcare AKTA FPLC system. The proteins apo- or holo-PiB samples (100 µl sample size, 400 µM) were injected onto the column and eluted with TBS buffer mobile phase at a flow rate of 0.5 mL/min with UV detection on 280 nm and 360 nm corresponding to protein and rucaparib UV absorbance (**Fig. S4**).

Native mass spectrometry

The proteins apo- and rucaparib-bound PiB (50 µL, 20 µM in TBS) were buffer exchanged into 200 mM ammonium acetate using Bio-Spin P-6 gel columns (Bio-Rad) according to the manufacturer's instructions. Briefly, the column was washed with 0.5 mL water and equilibrated with 200 mM ammonium acetate four times, prior to loading 50 µL sample and centrifuging at 1000 rcf for 4 min. 10 µL sample was loaded into the borosilicate emitter (ThermoFisher ES380) and analyzed on an Exactive Plus EMR Orbitrap mass spectrometer (Thermo Fisher). The instrument was operated in positive ion mode using the recommended setting. The typical parameters were set with a mass range of 1,000-10,000 *m*/*z*, a resolution of 8,750 (at *m*/*z* 200), spray voltage of 1.1-1.5 kV, trapping gas pressure of 5. The in-source fragmentation and HCD collision energy were turned off. The raw native mass spectra were deconvoluted in UniDec (*46*) to produce zero-charge spectra.

NMR

The NMR samples were prepared from isotope labeled M9 medium as described and concentrated to 0.5 mM in TBS buffer (50 mM Tris, 100 mM NaCl, pH 7.4) with 5% *d6*-dimethylsulfoxide.

The 2-dimensional $^1H$-$^{15}N$ HSQC spectra were collected on Bruker 800 MHz spectrometer with parameters: carrier 120 ppm, spectra width 35-40 ppm, ~2 hours. The 2-dimensional $^1H$-$^{13}C$ HSQC spectra were collected on the same instrument with parameters: carrier 45 ppm, spectra width 70 ppm, ~2 hours.

Determination of binding dissociation constant

We used spectral titration experiments to determine the binding dissociation constants for rucaparib, mefuparib, niraparib and veliparib with PiB, PiB' and the PiB mutants. The protein concentrations were determined by fitting linear equations with a serial measurements of absorbance 280 nm at different concentrations.

For rucaparib, we performed fluorescence emission experiment by a BioTek Synergy Neo2 Reader with excitation wavelength of 355 nm. The aliquots of protein from 1 μM or 2 μM stocks in TBS were added up to 160 μL solutions with rucaparib concentrations at 50 nM and 100 nM, respectively. Three to four replicates were performed for SD calculation. The changes in emission at 420 nm of the bound complex was used to fit to a single-site, protein-ligand model (*10*). For PiB mutants, we performed fluorescence emission experiments with rucaparib at 100 nM concentration.

For mefuparib, aliquots of the drug from 300 μM or 600 μM stocks in DMSO (final %DMSO < 2%) were serially added to 2 mL solutions of PiB or PiB' at 1 μM and 2 μM concentration. And the absorbance signal at 336 nm was detected twice per point using a Cary 300 Bio spectrophotometer in a 10-mm path-length quartz cuvette. The change in absorption at 336 nm of the bound complex relative to free mefuparib and PiB or PiB' was used to fit to a single-site, protein-ligand binding model (*10*). Similarly, the 600 μM and 1500 μM stocks of niraparib were used to titrate PiB or PiB' with concentration at 2 μM and 5 μM where the Kd is determined by fitting changes of absorption at 317 nm, as well as 4 mM and 6 mM stocks of veliparib were used to titrate PiB with concentration at 15 μM and 20 μM where the Kd is determined by fitting changes of absorption at 312 nm.

Circular dichroism

Proteins were prepared in 10 μM concentrations in buffer 1.25 mM Tris, 2.5 mM NaCl, and pH 7.4. 300 μL proteins are added in a 0.1 cm path length quartz cuvette for CD spectra collection using a J-810 CD spectrometer. Full spectra were collected from 200 nm to 250 nm in continuous scanning mode for PiB-1, PiB, PiB-2 and PiB' (**Fig. S3**). The parameters for full spectra collections were set up as follows: bandwidth is set to 2 nm, the scanning speed is set to 50 nm/min, and the average of accumulations is set to 3. Temperature-dependent spectra were collected from 20 to 95 °C using temperature/wavelength mode for bound and unbound PiB (**Fig. S4**). The parameters for temperature-dependent spectra collections were set up as follows: interval is set to 5 °C, temperature increase rate is set to 2 °C/minute, and wavelength range from 200 to 250 nm. Rucaparib-bound protein solution contained 10 μM rucaparib.

Cell viability assay

We tested five PARPi, including four that bind the PiB and PiB' (two approved by FDA, rucaparib and niraparib, and two under clinical trials, veliparib and mefuparib) and olaparib that does not bind as a negative control (*18, 47*). DLD-1 BRCA2 mutated cells or SUM149 cells (Horizon, Cat# HD-1005-007) (*25*) were cultured in RPMI 1640 medium (Gibco, Cat# A10491) supplemented with 10% FBS (Corning, Cat# 35-010-CV) and 1% Pen Strep (Gibco, Cat# 15140-122) in 37°C,

5% $CO_2$ incubator. Cells were plated at the density of 1,000 cells/well in 96-well plates. One day after plating, five PARP inhibitors, rucaparib, niraparib, veliparib, mefuparib and olaparib in the same concentration were pre-incubated with increasing ratios of PiB or PiB' (ratios of protein/Ligand include 1:1, 2.5:1, 5:1 and 10:1) at room temperature for 5 minutes. Cells were then treated with serial 4-fold dilutions of drugs alone or the mixture of drugs and PiB or PiB'. After 7 days of treatment, cell confluence was measured using an IncuCyte Live Cell Imager (Essen Bioscience). Confluence values were normalized to that of cells treated by DMSO vehicle. Half maximal inhibitory concentration $(IC)_{50}$ concentrations were calculated using GraphPad Prism 9 (RRID:SCR_002798).

<u>Unbiased classical molecular dynamics</u>

The MD system was prepared using Gaussian09 and AmberTools's Antechamber program in Amber18 (*48, 49*). The four compounds were parameterized by first optimizing their geometries at the B3LYP/6-31G* level, and then calculating their electrostatic potentials using the Merz-Singh-Kollman method in Gaussian09 (*50*). Charge fitting was then performed using Antechamber's RESP program within AmberTools (*51, 52*). All other small molecule parameters were assigned by Antechamber based on the GAFF2 database (*53, 54*). The starting structures for the MD simulations were derived from the rucaparib-bound PiB design model. Structures of PiB in complexes with the other three ligands were modeled by superimposing the ligands onto rucaparib, and close contacts were alleviated during minimization, as described below.

The simulation box was built by solvating each drug complex with TIP3P-modeled waters (*55*) in a box with 8 Å padding from the protein, and the system was charge-neutralized. All simulations were performed in Amber18 with the ff14SB and GAFF2 forcefields (*54, 56*). Simulations began with 1,000 restrained steepest-descent minimization steps before switching to a maximum of 6,000 steps in conjugate gradient steps. The system was then heated up to 293 K over 50 ps in the NVT ensemble with Langevin thermostat control of temperature, using a 1 fs timestep. The simulation was then switched to the NPT ensemble, and pressure was maintained at 1 atm using the Monte Carlo barostat (*57*). Throughout equilibration steps, protein and drug heavy atoms were initially restrained with harmonic potentials at 15 kcal/(mol·Å2), and ramped down to 0 kcal/(mol·Å2) over 6 equilibration steps, totaling 200 ps. Each simulation was then carried out for an unrestrained production run under periodic boundary conditions with 2 fs timesteps. Three independent 2 microsecond simulations were performed for each drug complex. The SHAKE algorithm (*58, 59*) was used to restrain hydrogens, and the Particle Mesh Ewald method (*60*) was used for long-range electrostatics, cutting off short-range non-bonded electrostatic and Lennard-Jones interactions at 10 Å. Trajectory analysis was performed using the python packages MDAnalysis (*61, 62*) and ProDy (*63*).

<u>X-ray crystallography</u>

The original design protein PiB had no crystals obtained in 96 well hanging drop trays from NeXtal JCSG Core Suite I, II, and III and Hempton Reseach PEG/ION 1 and PEG/ION 2. PiB showed no precipitate in the hanging drop with a concentration range from 10 mg/ml to 80 mg/ml.

We redesigned the surface of PiB and purified the redesigned protein PiB' using the same method described above. His-tag-cleaved PiB' was concentrated in buffer (10 mM Tris, 20 mM NaCl, pH 7.4) at 60 mg/ml. DMSO were removed by FPLC or filters for holo-protein sample. Crystals of PiB' showed in multiple conditions from NeXtal JCSG Core Suite II, PEG/ION 1 and PEG/ION 2. The crystals were further scaled and optimized in the following conditions: a) 2.0 M ammonium sulfate, b) 2.0 M ammonium sulfate with 0.1 M Sodium acetate trihydrate (pH 4.6). Crystals were harvested and cryo-protected in 15% glycerol in the crystallization buffer for 30 seconds before being flash-frozen in liquid nitrogen.

Data collection was performed remotely with the BOS/B3 software at Advanced Light Source (ALS) using beamline 8.3.1 at a wavelength of 1.11583 Å (11111 eV). The 720° rotation with 0.1° rotation interval was employed. Reflections were processed and merged via the automated XDS program (*64*). The structure was then solved by molecular replacement with Phaser using the PiB' design model (*65*). The structure was refined several rounds in Phenix and Coot with manual adjustment (*66, 67*). The protonated ligands were generated with eLBOW under Phenix or Coot (*67, 68*). Diffraction data and refinement statistics are shown in **Table S2**.

Calculation of solvent accessible surface areas

The ligand solvent accessible surface area (SASA) is calculated with SASA (*29*) module of Biopython-1.79 using default parameters (probe_radius=1.40, n_points=100).

Binding free energy calculations using the alchemical transfer method

The host–guest systems were prepared for each ligand from their respective crystal structure. Each crystal structure was superimposed based on the backbone $C_{alpha}$ atoms. No steric clashes due to rearrangement of side chains upon ligand binding was identified. For free energy calculations, protein coordinates were taken from the mefuparib protein complex. Super imposed ligands were stripped from the protein and served as the initial conformations for quantum mechanical optimization. Ligand geometry was optimized at the WB97X-D3level (*69*) using Psi4 1.8.0 (*70*) and simple-DFTD3 1.0 (*71, 72*). A restricted electrostatic potential (RESP) plugin (*73, 74*) for Psi4 was used to calculate RESP charges via a two-stage charge fitting protocol for the optimized ligand conformer. Due to each ligand being charged in the complex, the net charge of each ligand was set to +1 with a multiplicity of 1. Additionally, the crystal structure ligands were truncated to retain the fused ring core that forms the majority of protein ligand contacts. These truncated cores were parameterized under the same scheme as the full ligands. A modified AToM-OpenMM package (*75–77*) was used to prepare and run both sets of free energy calculations. AmberTools23 (*78*) was used to generate simulation input files. Force field parameters for small molecules were generated using antechamber with the derived RESP charges and GAFF2 force field (*79*). Protein residues were treated using the Amber14SB force field (*56*). The TIP3P water model (*55*) was used to generate the solvation box. Each complex system is built in LEaP and consists of the target receptor and a pair of pre aligned ligands. Based on the transformation, one ligand is selected to be translocated based on a displacement vector. Due to the long range electrostatics cut off being 10 Å, it was chosen to translocate one ligand such that the minimum distance between the protein and ligand would be over 20 Å. For these systems, this corresponded to a translocation vector of [-25.0 Å, 20.0 Å, -25.0 Å] or a magnitude of ~40 Å from the bound ligand. This ensured that at least three layers of water molecules were present between the protein and the translocated ligand, and that the protein or translocated ligand were exposed to the only long range electrostatics of the solvent and not of other solutes. An additional 10 Å solvent buffer was added to form the periodic cell. Sodium and chloride ions were added to neutralize the system. Solvated complexes were saved as Amber parameter/topology (prmtop) and Amber coordinate (inpcrd) files.

All alchemical molecular dynamics calculations employed the OpenMM 8.0 molecular dynamics engine (*80*) and the alchemical transfer method meta-force OpenMM plugin (*75*) using the CUDA platform. The ASyncRE software (*77*) was used for the Hamiltonian Replica Exchange in λ space.

Initial complexes were minimized and slowly heated to a target temperature of 300 K. Each system is then annealed from the bound state (λ = 0) to the symmetric alchemical intermediate (λ = 0.5) for 250 ps. This symmetric alchemical intermediate is the starting point for the Hamiltonian Replica Exchange in λ space. Two sets of constraints are to be applied to accelerate convergence of the binding free energy estimates. Flat-bottom distance restraints with a harmonic potential of 25 kcal/mol and a distance tolerance of 1.5 Å are applied to the $C_{alpha}$ atoms of residues within 3.0 Å of the ligand. Additionally, these restraints are used to fix the orientation of the ligand by three chosen reference atoms that define the center of mass of the ligand, the plane of the constrained core, and the central axis of the ligand, respectively. A soft plus alchemical potential (*81*) was used for all lambda windows for bound state (λ = 0) to the symmetric alchemical intermediate (λ = 0.5) and from the symmetric alchemical intermediate (λ = 0.5) to the unbound state (λ = 1). Asynchronous Hamiltonian replica exchanges are performed every 10 ps. A Langevin thermostat with a time constant of 2 ps was used for temperature regulation. A total of ~40 ns per relative binding free energy transformation was sampled. This is in line with the typical FEP+ (36 or 60 ns per ΔΔG) (*30*) or Amber thermodynamic integration ( ~50 ns per ΔΔG) (*31*) A total of ~150 ns per absolute binding free energy transformation was sampled. This corresponds to a minimum of 7.5 ns per lambda window. Binding free energies and their corresponding uncertainties were calculated from the perturbation energy samples take at the point of replica exchange using the UWHAM (*82*) method. Samples were take from the second half of each trajectory. All simulations were ran on two Nvidia RTX 3080 graphics cards using driver version 525.116.04 and CUDA 12.0.

Potential of mean force (PMF)

Simulations were conducted using the GROMACS package, version 2022.4 (*83*) with the CHARMM36m force field (*84*). The ligand parameters were generated utilizing CgenFF (*85*). CM5 charges (*86*) calculated at the M062X/cc-pVDZ level of theory (*87, 88*) using Gaussian 16 (*89*) were applied to the ligands. Short-range nonbonded interactions were truncated at 1.2 nm, while long-range electrostatics were computed using the particle mesh Ewald (PME) algorithm (*90*). Periodic boundary conditions were implemented along all dimensions. Each protein-ligand system was enclosed within a rectangular box of TIP3P water (*55*) with dimensions sufficient to adhere to the minimum image convention and allow for pulling simulations along the z-axis. Neutralization of charge was achieved by adding Na+ and Cl- ions. The same z-axis orientation was used as the pulling direction across all systems. This orientation was established based on the rucaparib-bound PiB configuration, aligning it in such a way as to promote the direct dissociation of the drug from the binding pocket while minimizing potential clashes. This criterion ensured a standardized approach to evaluating the unbinding process. Although the ligand was primarily dissociated by increasing center-of-mass (COM) distance between the protein and ligand along the z-axis (*vide infra*), it retained the freedom to move in other axes (Fig. S29). The z-axis orientation in our simulations—uniform across all systems and aligned with the rucaparib-bound PiB configuration—was chosen to streamline the dissociation analysis. This approach facilitated a consistent and efficient evaluation of the unbinding process. Our primary goal was to ensure reproducible and accurate results, focusing on a methodologically sound framework for binding energy calculations, rather than an exhaustive exploration of all possible orientations (*91*).

Our calculation of the binding energy ($\Delta G_{bind}$) was based on a methodology used in earlier studies, especially for the equilibration and pulling simulation steps (*92*). Our approach differed in a few ways. We used a different water model and capped the N- and C-termini with amine (-

NH3+) and carboxylate (-COO–) groups, respectively. Also, a lower spring constant was chosen for the pulling force to enhance the precision in creating the dissociation pathway. After an initial steepest descents minimization step, each system was equilibrated under NPT ensemble conditions for 100 ps (T=300 K and P=1 bar) using the Bussi-Donadio-Parrinello thermostat (*84*) and the Parrinello-Rahman barostat (*93*). Following equilibration, restraints on the ligand were removed, enabling it to be pulled away from the protein along the z-axis over a period of 500 ps. A spring constant of 200 kJ $mol^{-1}$ $nm^{-2}$ and a pull rate of 0.01 nm $ps^{-1}$ (0.1 Å ps−1) were used, achieving a final center-of-mass (COM) distance between the protein and ligand of approximately 5.5 nm. Snapshots from these trajectories were used to generate initial configurations for the subsequent umbrella sampling.

For umbrella sampling (*94*), a symmetric distribution of sampling windows was established, with a window spacing of 0.1 nm based on the COM separation. In each window, 10 ns of molecular dynamics (MD) simulations were performed. The binding energy, derived from the potential of mean force, was calculated by analyzing the umbrella sampling data using the Weighted Histogram Analysis Method (WHAM). (*95*). The binding energies showed a similar ordering to that observed with the alchemical transfer method, closely matching experimental results. The binding energies agree with experimental data (Table S5), with minor deviations observed for ligands interacting with PiB', leading to a modestly increased computed affinity for these complexes. These deviations may be linked to reduced fluctuations at the binding site, as reflected by a slightly lower root mean square fluctuation (RMSF) and C-alpha carbon root mean square deviation for drug-bound PiB' (Figs S31 and S32). An analysis of the structural differences between PiB and PiB' in the simulations used to compute the binding energies showed that the binding-site residues remain essentially identical between the two proteins (Fig S33); and there are modest differences in C-alpha RMSD between regions containing the Ala mutations, which might account for the slight shift in the computed binding free energies for PiB' (Table S5). Despite any small differences in the simulated structural ensembles, the relative ordering of the binding free energies was found to be the same between PiB and PiB'.

#### Supplementary Figures

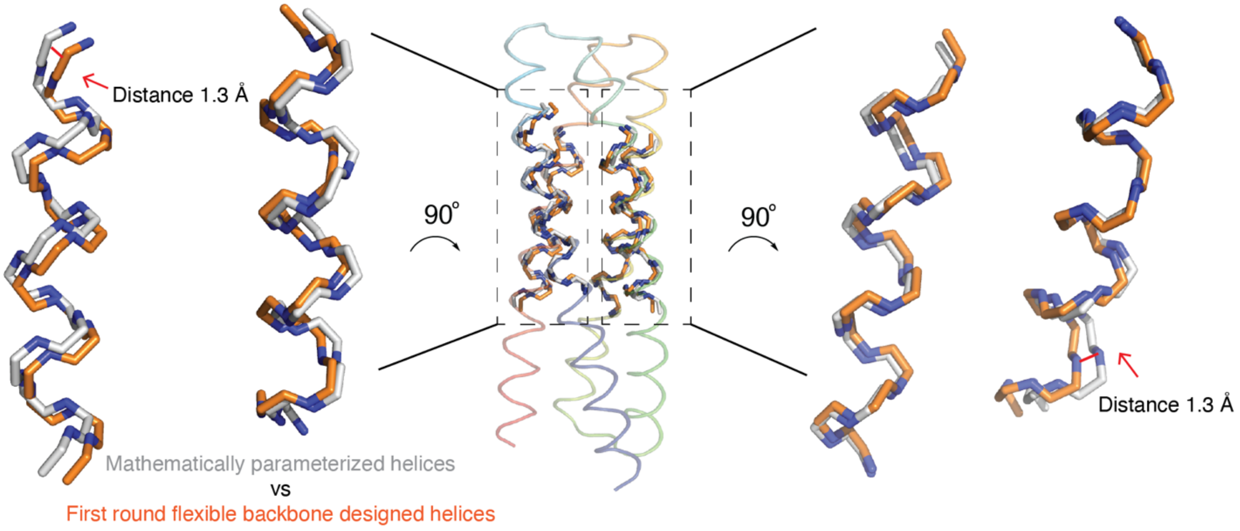


**Fig. S1. The flexible backbone design shifts the backbone and vdM optimization found better interactions.**

(A) Mathematically parameterized helices are first generated from the CCCP program. After the first round of flexible backbone design, the backbone shifted significantly (binding site Cα RMSD ~1 Å). The distance labeled in the figure is the atom-atom distance. Model for 60 residues (15 residues per helix) surrounding the drug binding position (residue numbers defined in Fig. 2) before and after flexible backbone sequence design.

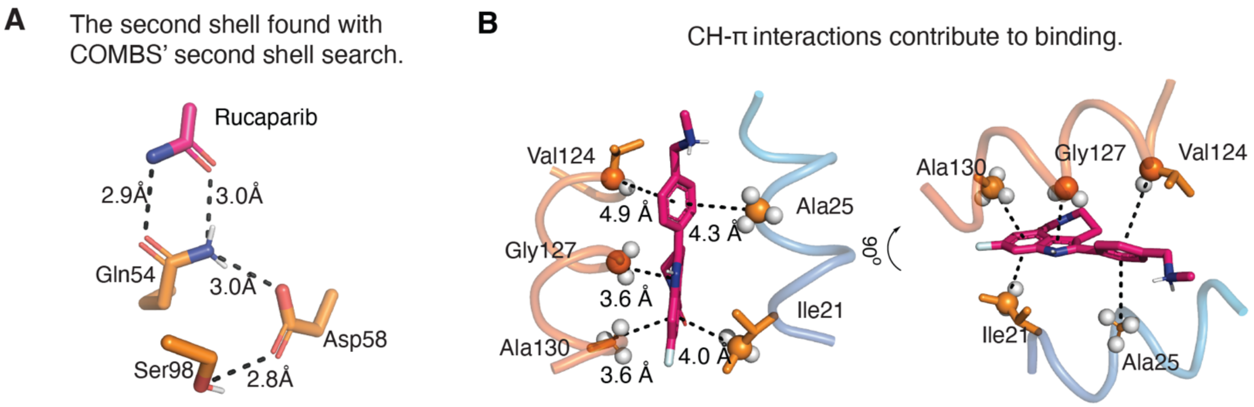


**Fig. S2. Design with second-shell hydrogen bonds, van der Waals and CH-π interactions.**

(A) COMBS identifies Asp58 as a second shell which forms hydrogen bond with Gln54 and Rosetta built the third shell Ser98. (C) In the design, rucaparib forms multiple CH- π interactions (Ile21, Ala25, Val124, Gly127, Ala130) with the helix backbone or sidechains.

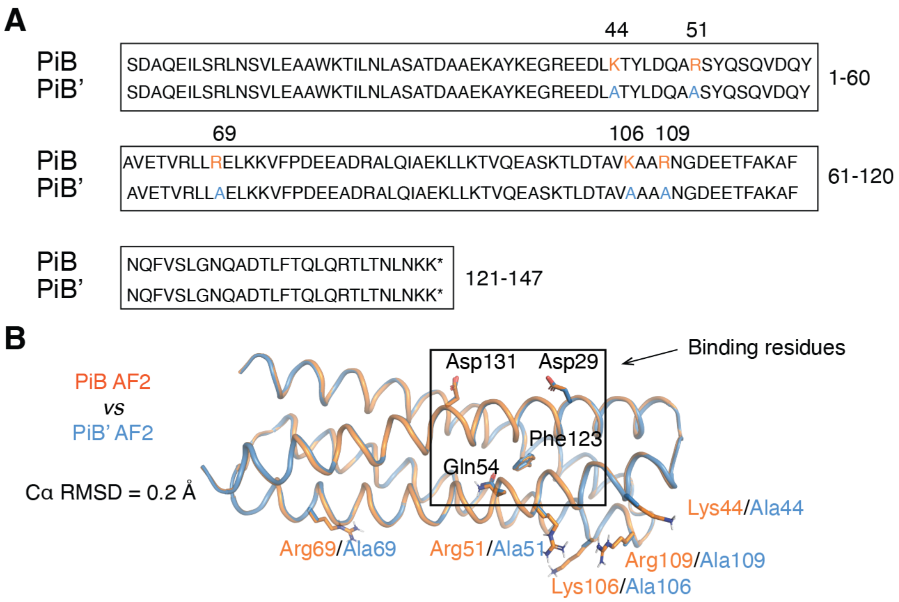


**Fig. S3. The sequence and predicted structure of PiB’ and PiB.**

(A) Sequence comparison between PiB and PiB’. Two Lysine (Lys44, Lys106) and three arginine (Arg51, Arg69, Arg109) were substituted to alanine to decrease overall surface charge, minimize the presence of flexible side chains on the protein surface, and improve crystallization. (B) The AlphaFold2 predicted structure alignment of PiB and PiB’. The surface mutations did not affect the binding site sequence nor did they alter the overall structure. The surface charge of the protein changed from -3 in PiB to -8 in PiB’.

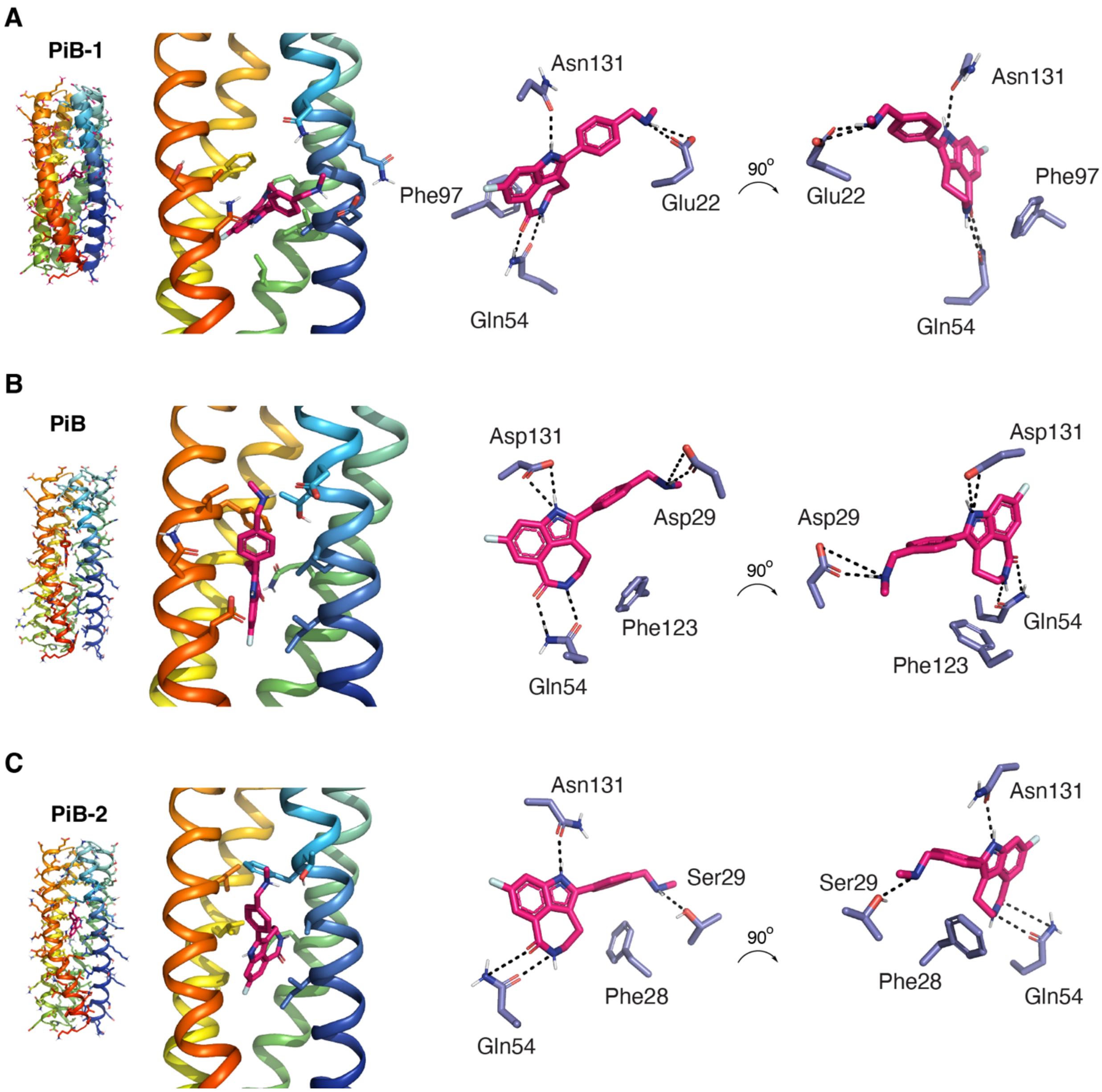


**Fig. S4. Computational models of the three designed proteins.**

The computational models of (A) PiB-1 (B) PiB (C) PiB-2. All the designs are the same length. The protein sequences and DNA sequences are listed in Table S1. We purposely selected designs that have all the polar groups satisfied with hydrogen bonds.

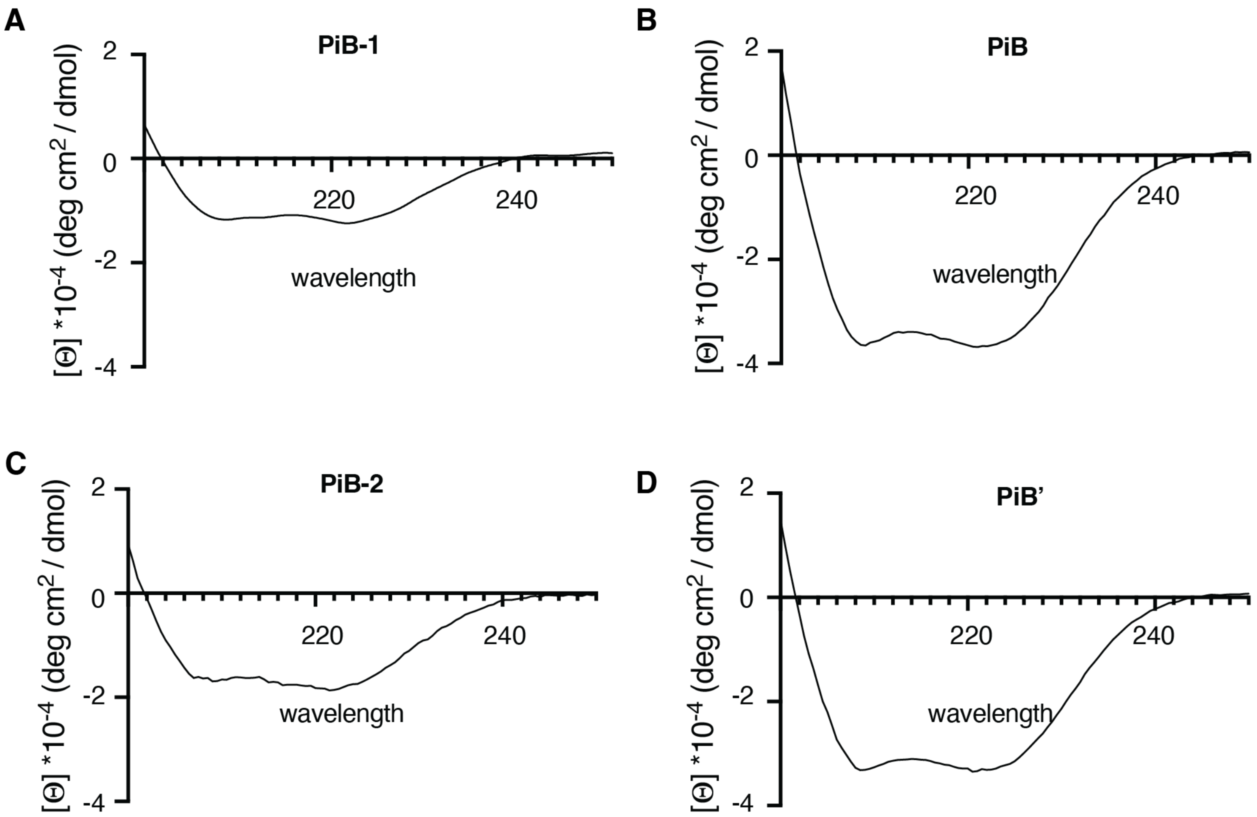


**Fig. S5. Circular dichroism spectra shows that all the designed proteins are helical.**
Spectra were collected at room temperature with 10 μM protein (in buffer 1.25 mM Tris, 2.5 mM NaCl, pH 7.4) in 0.1 cm path length quartz cuvette.

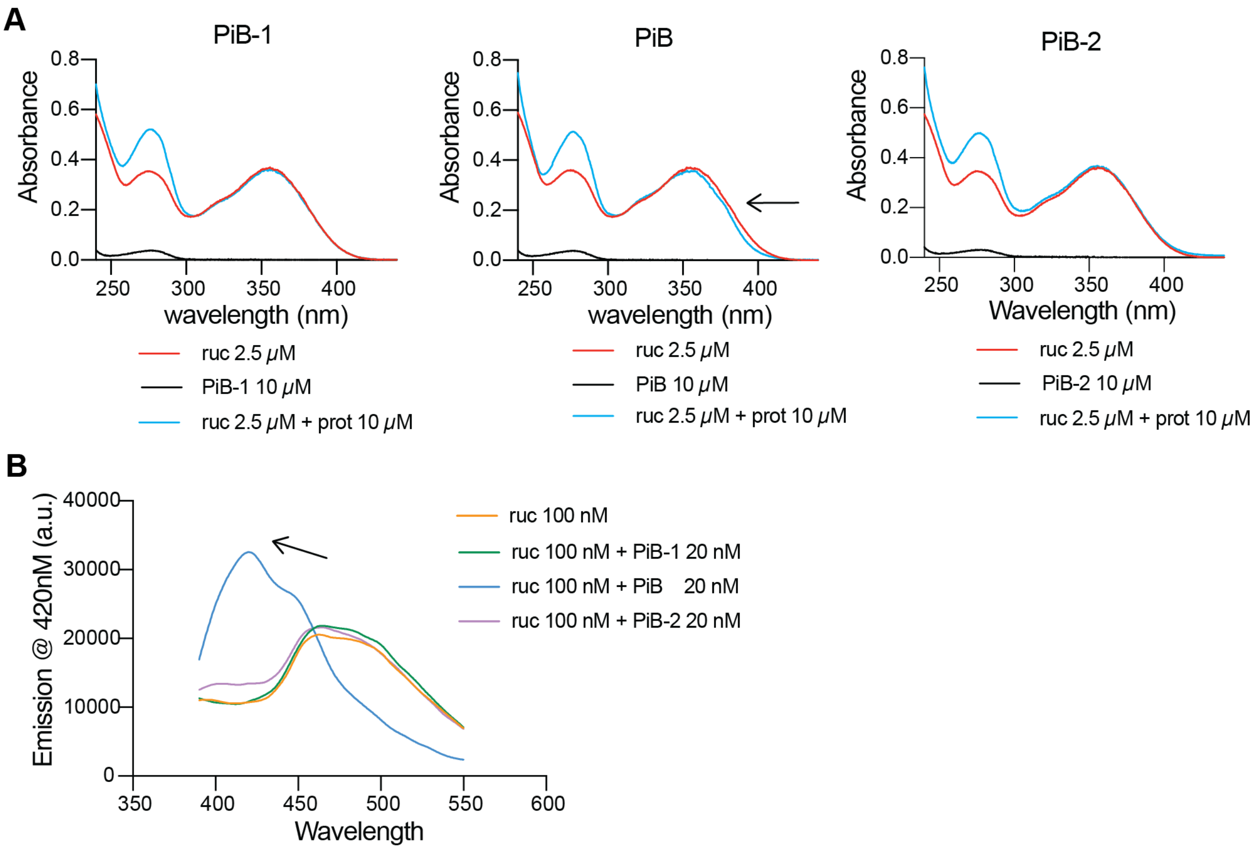


**Fig. S6. PiB bound with rucaparib and induced UV-VIS and fluorescence peak shifts.**

(A) Compared with PiB-1 and PiB-2, only PiB produced a small shift in the peak between 350 and 400 nm when added into 2.5 μM rucaparib (arrow in 1A). The electronic absorption spectrum was measured in 1.0 cm path length quartz cuvette. (B) PiB induced significant fluorescence emission shift from 480 to 420 nm. For both experiments, the drugs in DMSO are mixed with the buffer or proteins in TBS buffer (50 mM Tris, 100 mM NaCl, pH 7.4). with final DMSO concentration < 2% for 5 minutes before measurement. The fluorescence emission was measured in 96 well plates with excitation at 355 nm. All the spectra were measured at room temperature.

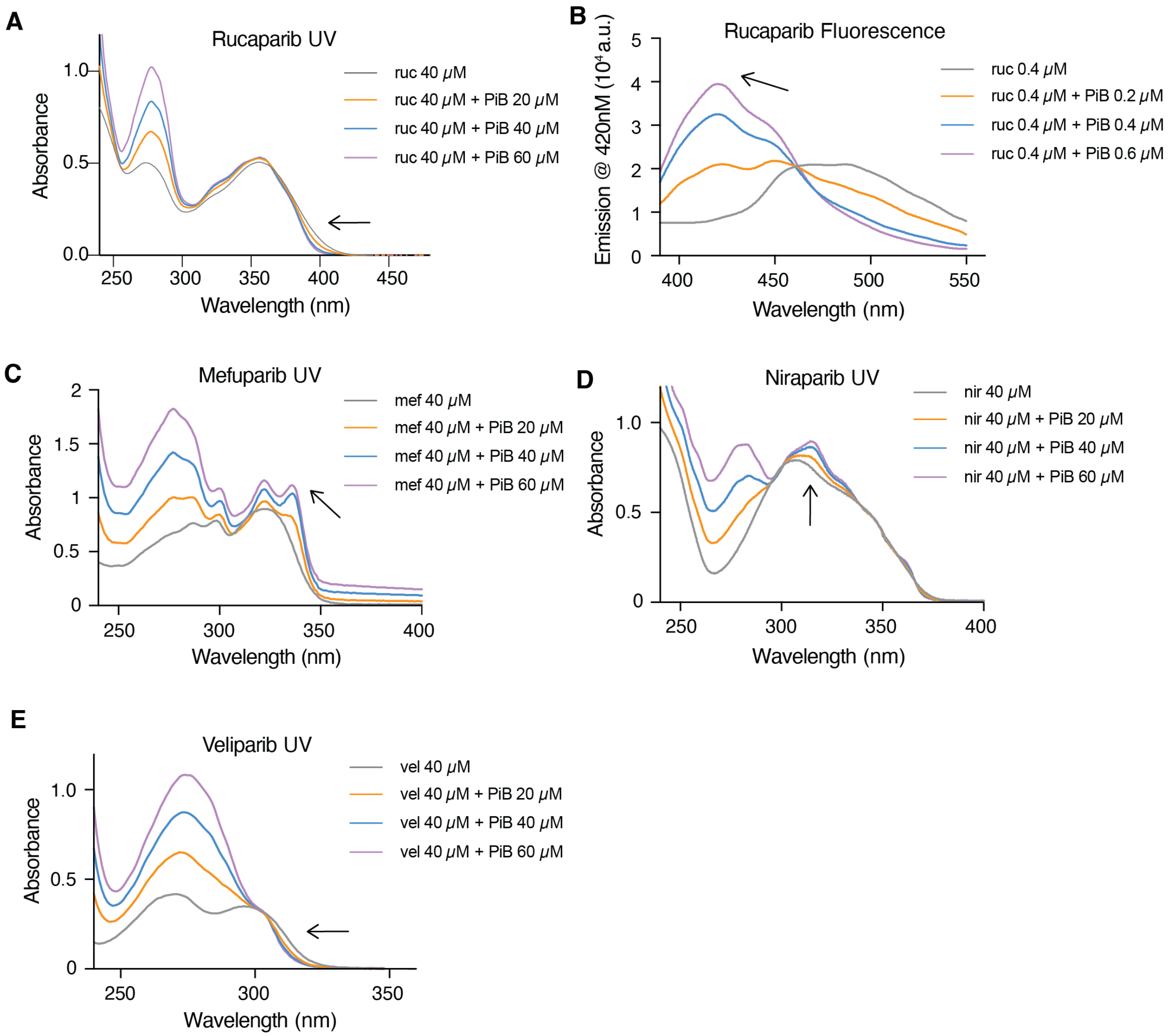


**Fig. S7. PiB bound the four PARPi ligands and induced changes in fluorescence or UV-VIS spectrum in a dose response manner.**

Electronic absorption spectra of PiB upon binding with (A) 40 μM rucaparib, (B) 0.4 μM rucaparib (fluorescence emission spectrum, excitation wavelength = 355 nm), (C) 40 μM niraparib, (D) 40 μM veliparib, (E) 40 μM mefuparib at a ligand:protein ratio of 1:0.5, 1:1, 1:1.5 and ligand alone. The electronic absorbance spectra of rucaparib and veliparib are blue-shifted upon binding PiB, while mefuparib and niraparib are red-shifted. The fluorescence emission spectrum of rucaparib is blue-shifted upon binding to PiB. The drugs in DMSO are mixed with the proteins in TBS buffer (50 mM Tris, 100 mM NaCl, pH 7.4) with final DMSO concentration < 2% for 5 minutes before measurement. The electronic absorption was measured in 1.0 cm path length quartz cuvette. The fluorescence emission was measured in 96 well plates. All the spectra were measured at room temperature.

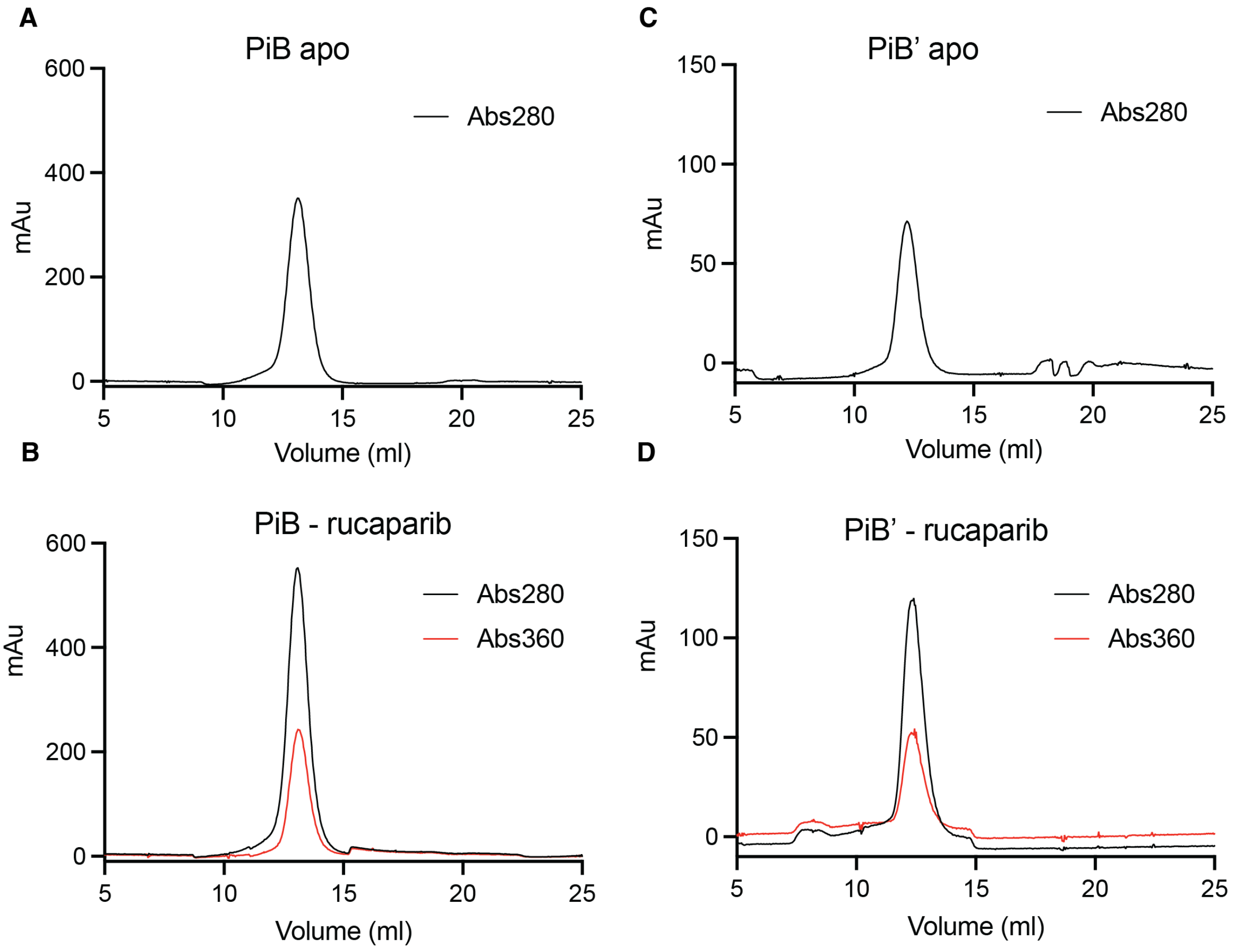


**Fig. S8. PiB and PiB' are monomers either in bound form or unbound form.**

Uncomplexed (A, C) and complexed PiB or PiB' (B, D; with 1.0 equivalent rucaparib for the complexed samples) are monomers based their elution as a single peak at 13.0 mL elution volume on a Superdex 75 Increase 10/300 column. The holo-PiB and PiB' elute together with rucaparib based on absorbance at 360 nm.

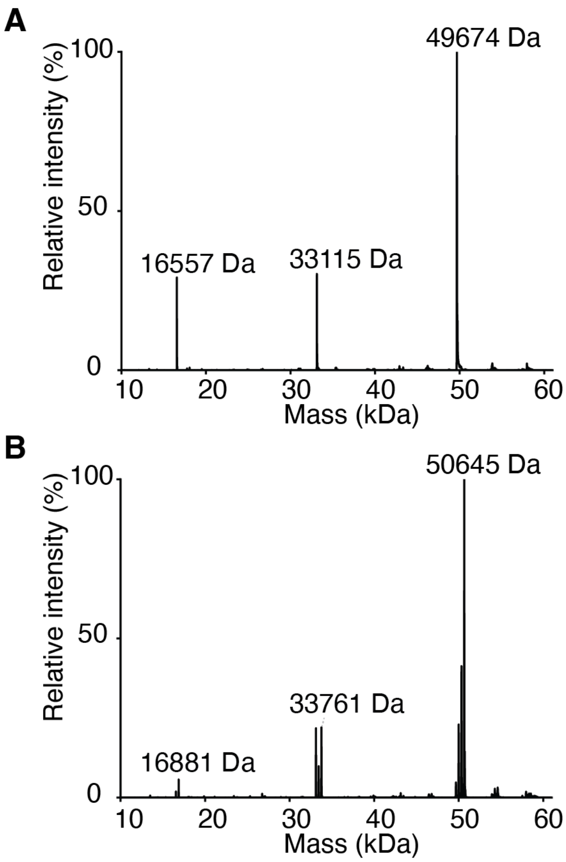


**Fig. S9. Native mass spectrometry of apo-PiB and rucaparib-bound PiB.**

Zero-charged native mass spectra of (A) apo-PiB and (B) rucaparib-bound PiB. For both the apo- and holo-PiB, we observed masses of monomer, dimer and trimer. The mass differences between apo- and holo-PiB are mass of one rucaparib for monomers, two rucaparib for dimers and three rucaparib for trimers. The proteins were buffered exchanged to 100 mM ammonium acetate and analyzed on an Exactive Pluse EMR Orbitrap mass spectrometry in positive ion mode with resolution of 8,750 (at m/z 200).

**A**

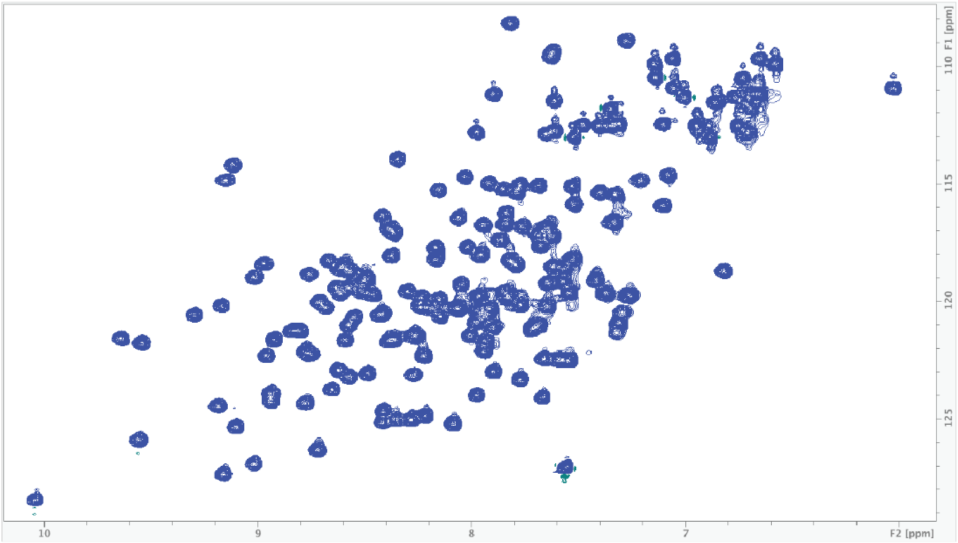


**B**

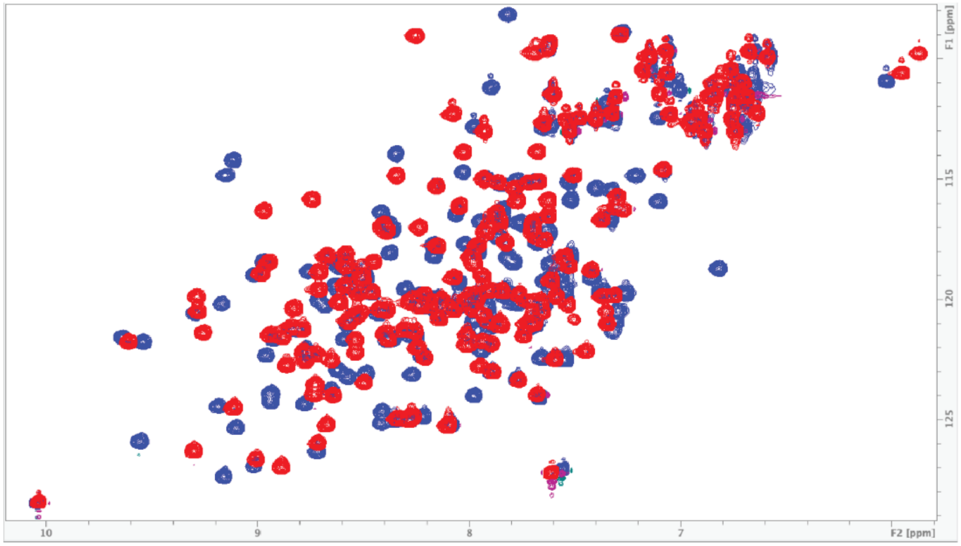


**Fig. S10. $^{15}$N-HSQC spectra of PiB and rucaparib-bound-PiB**.
(A) Spectrum of the drug-bound form and (B) Comparison between free (red) and complexed (blue) protein, showing ligand binding. The well-dispersed $^{15}$N-HN peaks spectra are typical of well-folded structures in solution. In B, peak positions show significant changes while the peak count is close to 140 upon addition of the ligand, demonstrating tight binding between the ligand and the protein. Both spectra were recorded on a Bruker 800 MHz instrument at 298K.

A

B

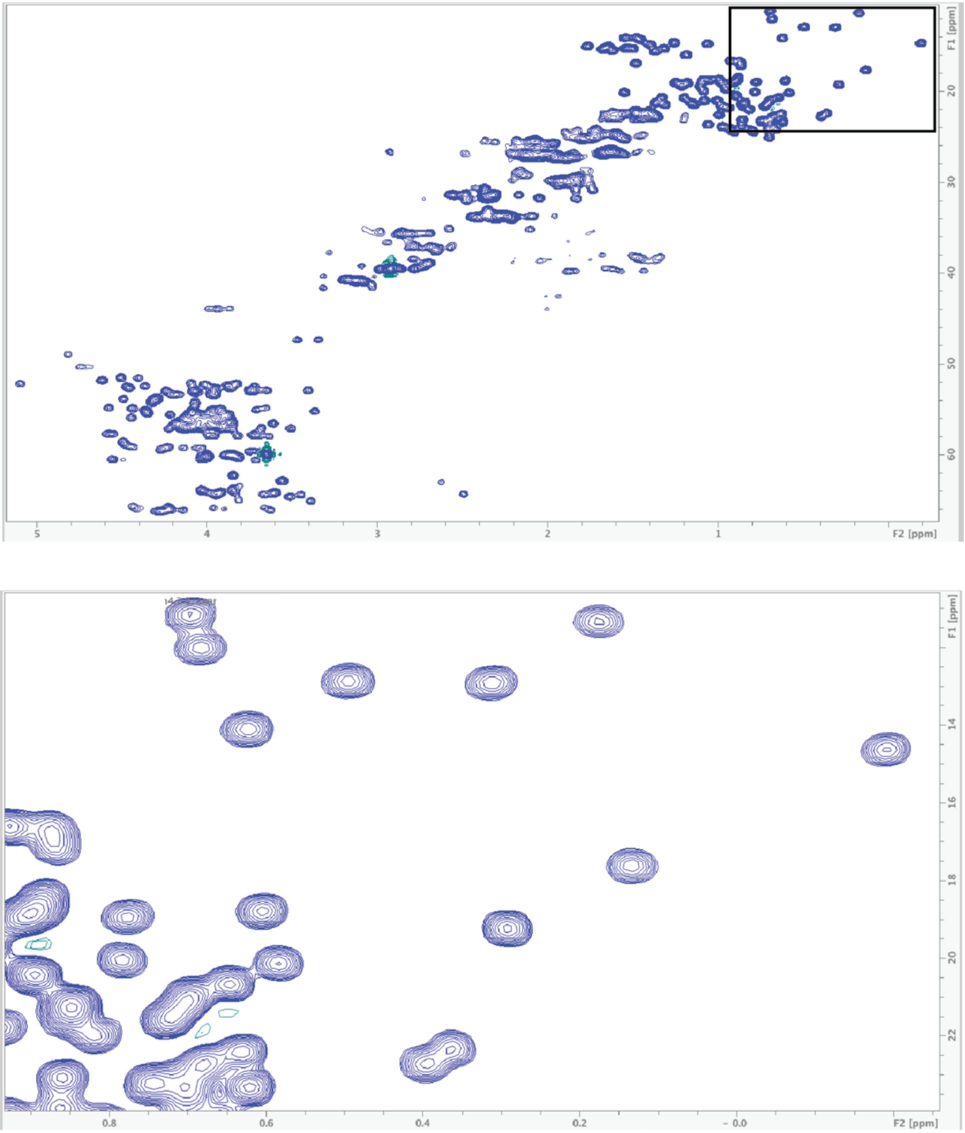

**Fig. S11. $C^{13}$-HSQC spectrum of rucaparib-bound-PiB (holo).**

(A) Spectrum of the complexed form in the aliphatic region and (B) Expanded view highlighting methyl groups. In A, the peak linewidth aligns with the expected value for the designed protein at this size. The CA-HA peaks indicate an alpha-helical protein, as most of HA chemical shifts are smaller than 4.70 ppm. Panel B reveals deviations in chemical shifts for certain methyl groups from the intrinsic position (0.7 ppm), with some even falling below 0.2 ppm. The significant chemical shift difference in methyl group chemical shifts reflect variations in the chemical environment and structural properties of those methyl groups within the designed protein, suggesting a well-packed protein core in solution.

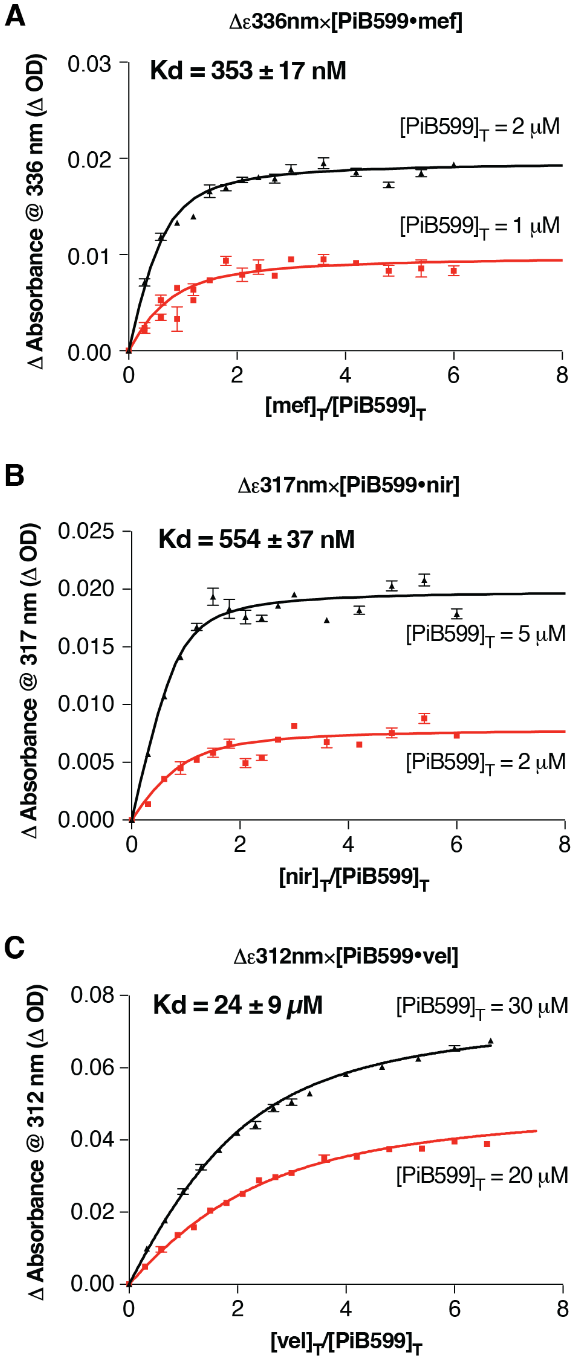


**Fig. S12. Spectral titrations of PiB' with PARPi.**

Determination of binding constants for various drugs for PiB' from global fits of a single-site binding model to absorbance changes as a function of the concentration of PiB'. Indicated wavelengths for the titration were chosen to maximize the difference in absorption for the free versus bound drug. The binding constant of PiB' to rucaparib is showed in Fig.2.

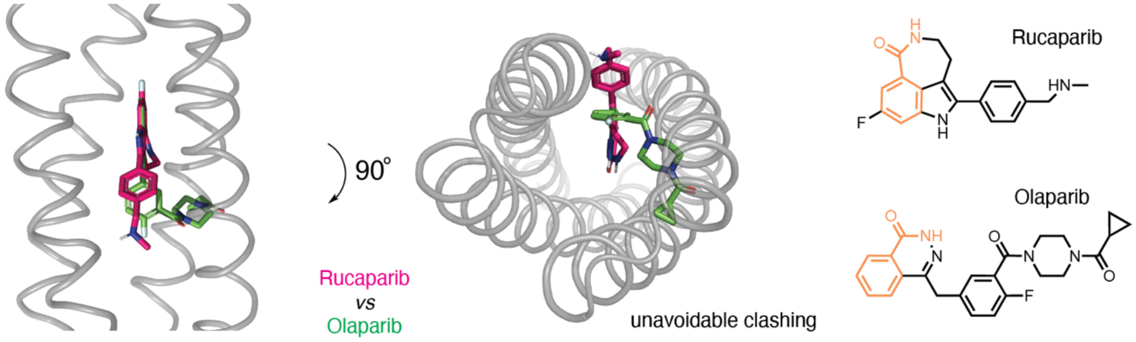


**Fig. S13. The model of olaparib showing close approach with the designed backbone.**

The structure of Olaparib (shown overlayed with the carboxamide chemical group on Rucaparib) makes it impossible to fit in the four helix bundles and thus is used as a negative control.

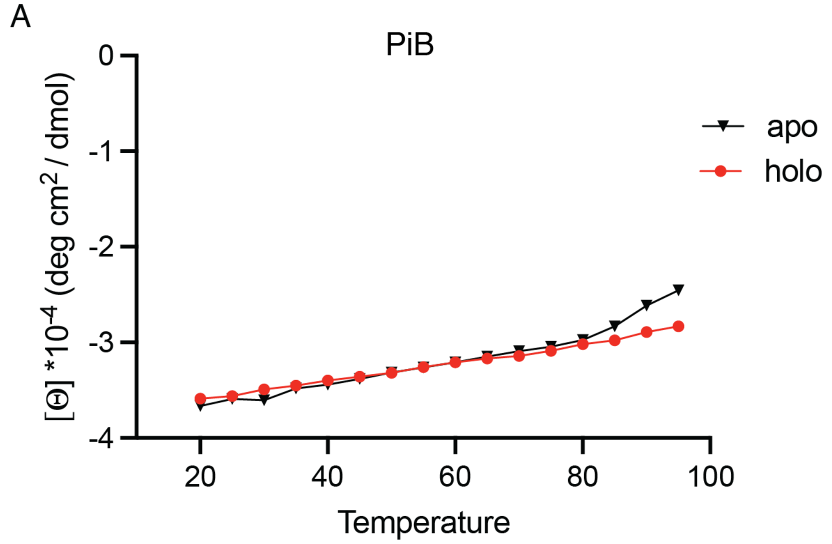


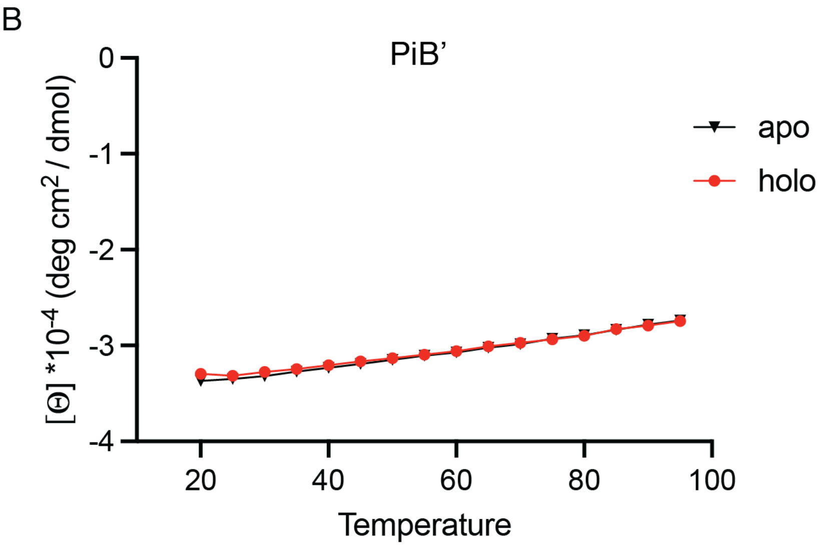


**Fig. S14. Free and rucaparib-bound PiB and PiB’ are thermostable.**

Temperature dependent circular dichroism signals measured at 222 nm shows that the apo- and rucaparib-bound PiB and PiB’ have melting temperatures > 80 °C. In the absence of rucaparib PiB’ is more stable than PiB, which shows the beginning of an unfolding transition near 80 °C. The enhanced stability is likely because PiB’s has five Ala mutations, which are known to stabilize proteins when placed on the surface of helices (*96*, *97*). In the presence of a single equivalent of rucaparib neither protein showed any unfolding. Proteins (10 µM concentrations in buffer 1.25 mM Tris, 2.5 mM NaCl, pH 7.4 with 0 or 1 eq. of rucaparib added) were measured in a 0.1 cm path length quartz cuvette. Temperature increase rate is set to 2 °C/minute during collection.

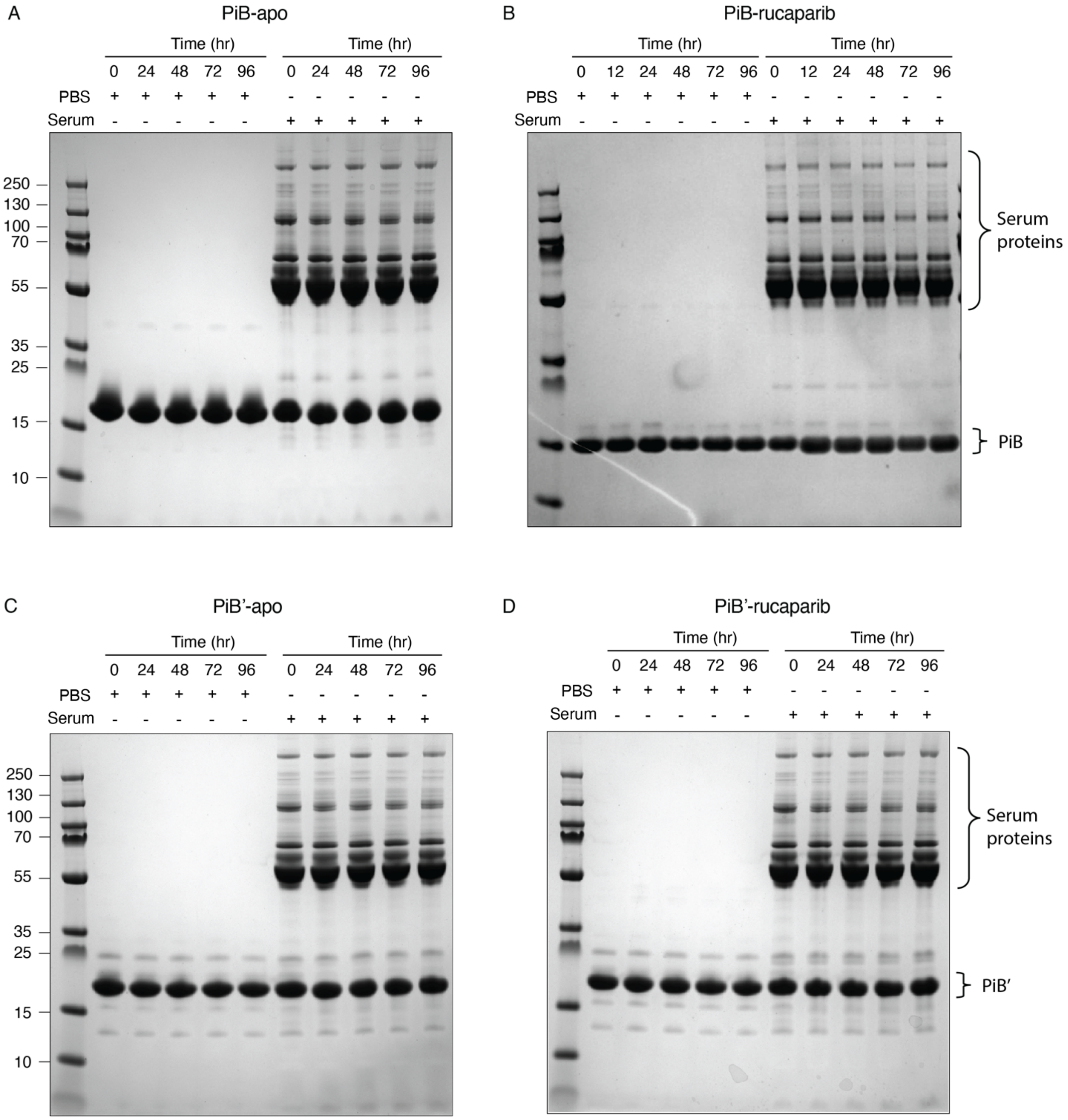


**Fig. S15. The rucaparib-bound PiB is stable in PBS and Serum for 4 days.**

We incubated 100 μM apo or holo PiB and PiB' (holo with added 100 μM Rucaparib on top of 100 μM protein) at 37 °C for 4 days and no degradation was observed. The left lane contains MW standards, followed by time points collected in PBS, and time points collected in serum. The extra bands at higher molecular weight in the serum samples are from native serum proteins.

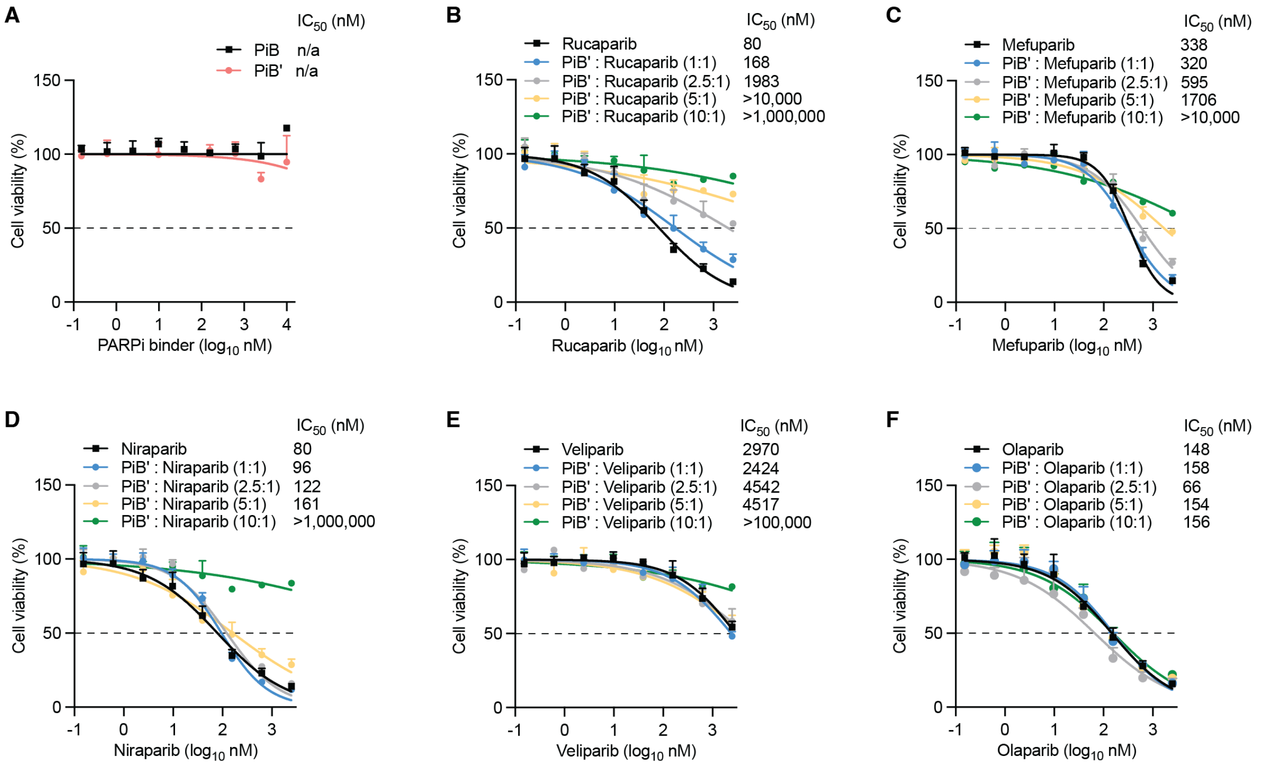


**Fig. S16. The cell viability assays of PiB and PiB' with PARPi in DLD-1 BRCA2 mutated cells.**

(A) Seven-day cell viability assays in DLD-1 BRCA2 mutated cells show that adding PiB and PiB' along doesn't affect cell viability. (B-F) PiB' alleviates the effects of rucaparib, mefuparib, niraparib and veliparib toxicity in a dose-dependent manner. The PARP inhibitors were pre-incubated with PiB' in media at room temperature for 5 minutes at multiple concentration ratios (protein: ligand) of 0:1, 1:1, 2.5:1, 5:1 and 10:1.

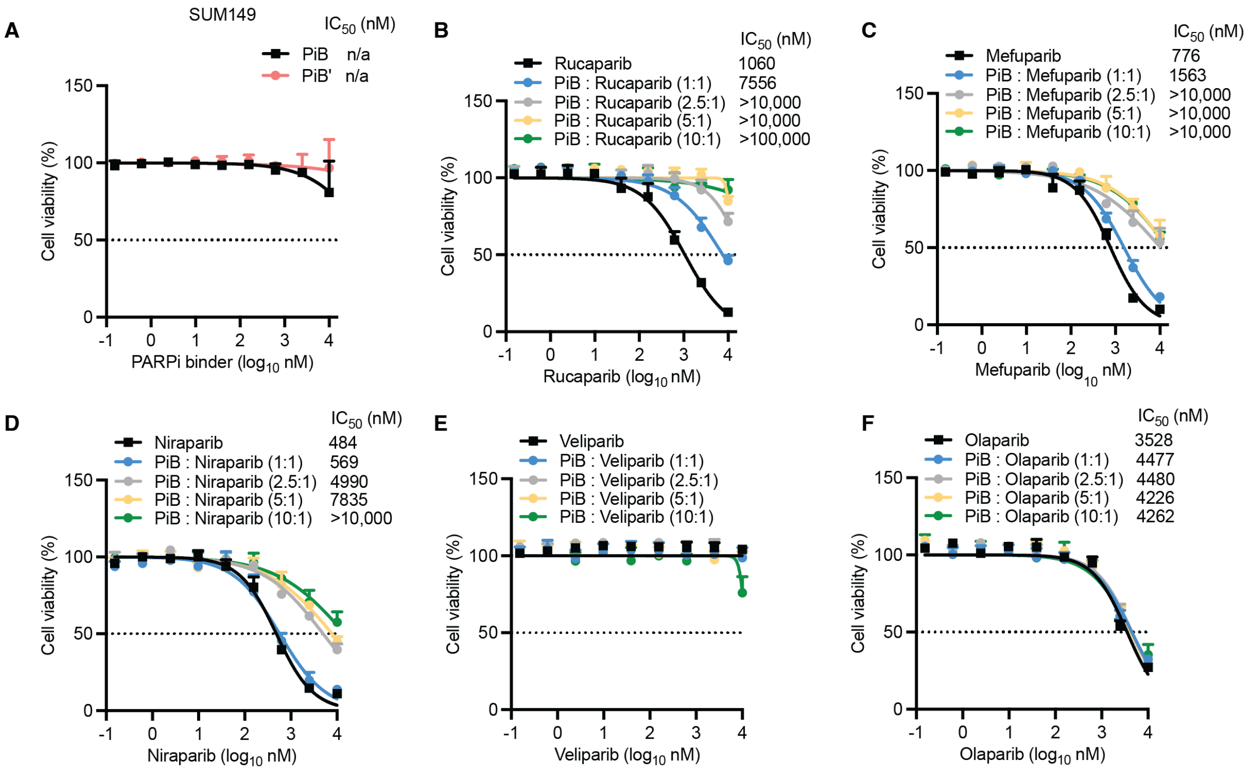


**Fig. S17. The cell viability assays of PiB with PARPi in SUM149 cells.**

(A) Seven-day cell viability assays in SUM149 cells show that adding PiB and PiB' along doesn't affect cell viability. (B-F) PiB alleviates the effects of rucaparib, mefuparib and niraparib toxicity in a dose-dependent manner. Veliparib does not show any effect on SUM149 cells. The PARP inhibitors were pre-incubated with PiB in media at room temperature for 5 minutes at multiple concentration ratios (protein: ligand) of 0:1, 1:1, 2.5:1, 5:1 and 10:1.

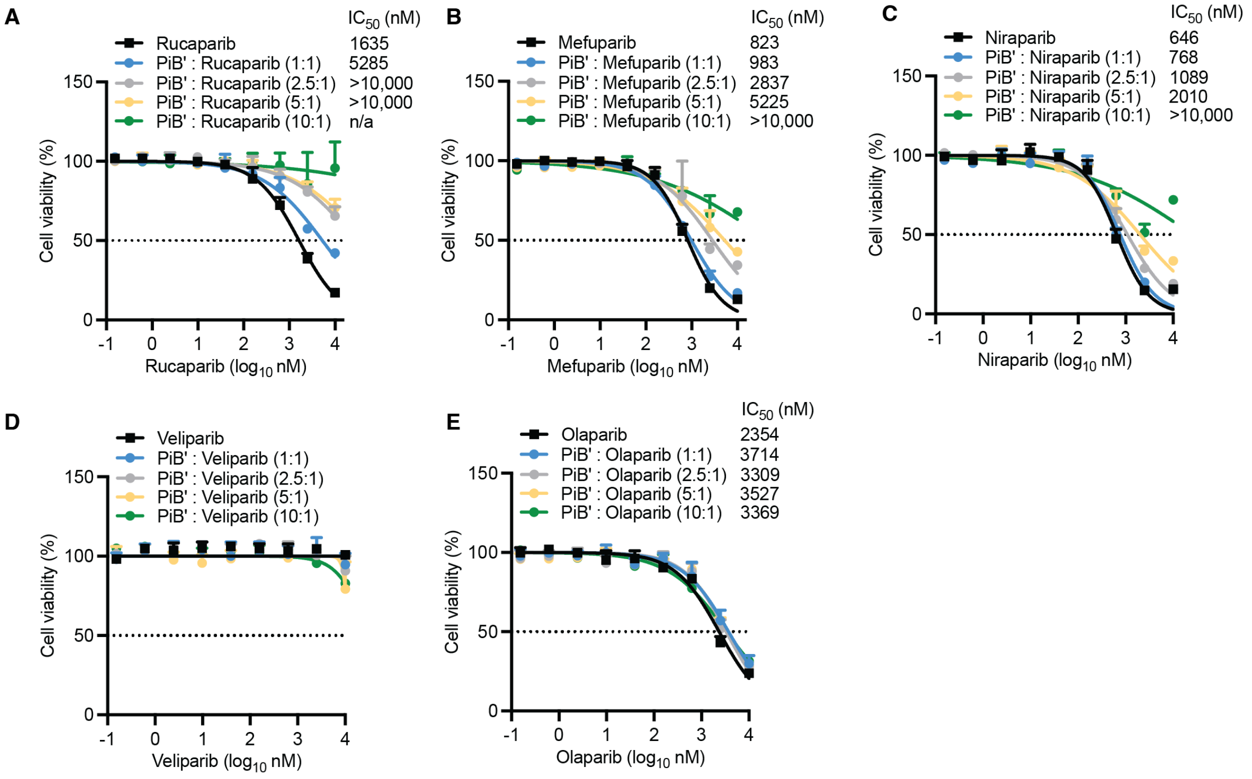


**Fig. S18. The cell viability assays of PiB' with PARPi in SUM149 cells.**

Seven-day cell viability assays in SUM149 cells show that PiB' alleviates the effects of rucaparib, mefuparib and niraparib toxicity in a dose-dependent manner. Veliparib doesn't show any effect on SUM149 cells. The PARP inhibitors were pre-incubated with PiB' in media at room temperature for 5 minutes at multiple concentration ratios (protein: ligand) of 0:1, 1:1, 2.5:1, 5:1 and 10:1.

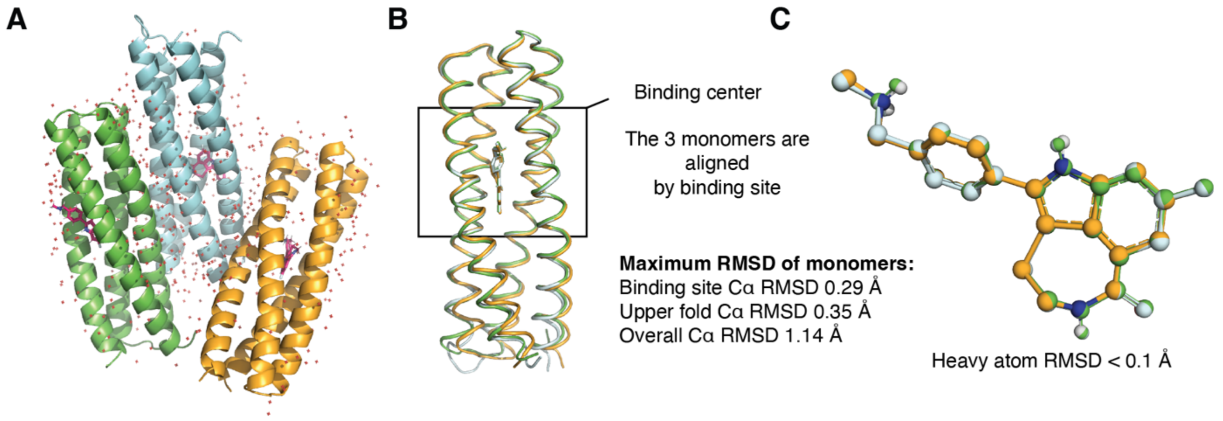


**Fig. S19. Crystallographic asymmetric unit of rucaparib-bound PiB'.**

(A) The asymmetric unit of rucaparib-bound PiB' contains three similar monomers. The monomers are colored in green, light blue and orange. Rucaparib is colored in pink. (B) The 3 monomers are superimposed on the chain A monomer binding site. The binding site has Cα RMSDs range from 0.15. to 0.29 Å. Greater deviation in Cα RMSD is seen near the N- and C-terminus and the middle loops (Panel B) (C) Rucaparib (colored in green, light blue and orange) from the three subunits by the protein superimposition shared the same conformation (heavy atom RMSD < 0.1 Å with heavy atoms).

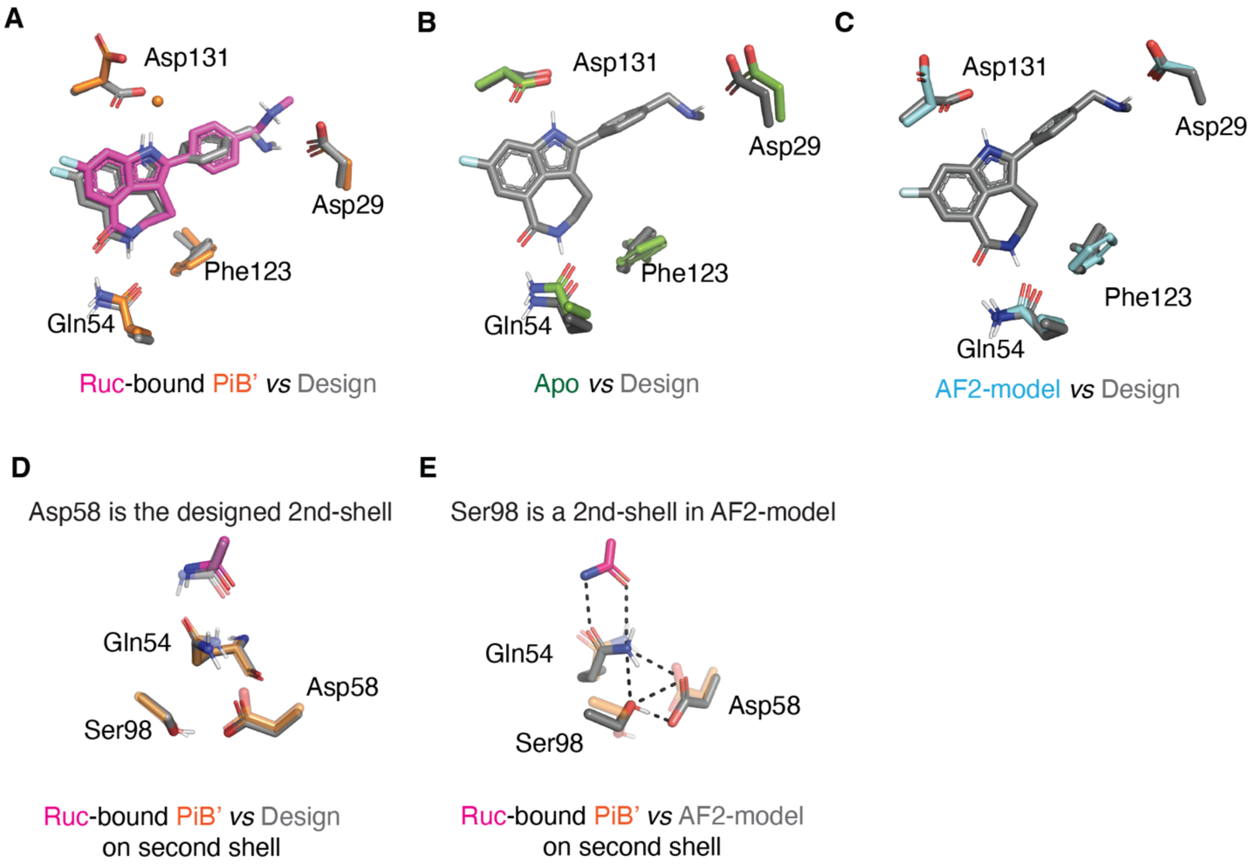


**Fig. S20. Sidechain comparison between rucaparib-bound PiB', apo-PiB' and AlphaFold2 model.**

(A) Overlay of designed interactions (gray) with the rucaparib-bound PiB' X-ray diffraction structure (protein in orange, rucaparib in hot pink). The Asp131 of the rucaparib-bound PiB' is in a different rotamer which forms a water bridged interaction with the rucaparib indole. (B) Overlay of designed interactions (gray) with the apo PiB' structure (green). The position of Asp31 in the apo PiB' agrees with the designed rotamer. (C) Overlay of designed interactions (gray) with the AlphaFold2 model (protein in cyan). The position of Asp131 of AlphaFold2 model agrees with the designed rotamer, despite AlphaFold2 having no information about the ligand existed. (D) The design model (gray) agrees with the rucaparib-bound PiB' (protein in orange, rucaparib in pink) regarding the second shell and third shell. (E) The AlphaFold2 model differs in the second shell and third shell compared with the rucaparib-bound PiB' (protein in orange, rucaparib in pink). Note that all the structures above were superimposed onto the binding site Cα atoms of the designed model before overlay.

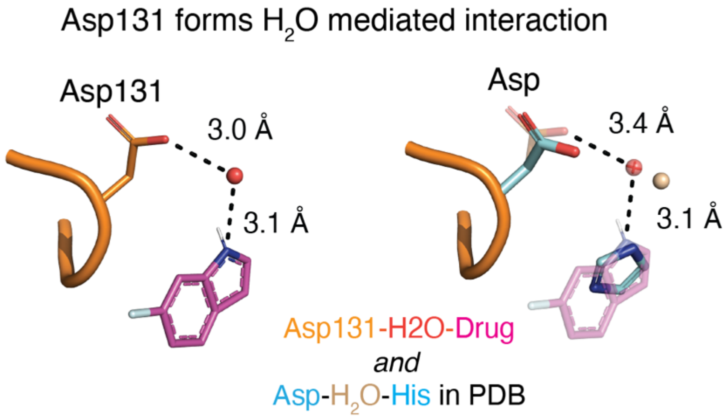


**Fig. S21. Water bridged interaction between Asp131 and indole NH of rucaparib.**
Asp131 in the rucaparib-bound PiB’ formed a water mediated interaction with the drug as in the MD simulations. Such water mediated interaction (Asp-$H_2O$-His) could be found in other structure from PDB database.

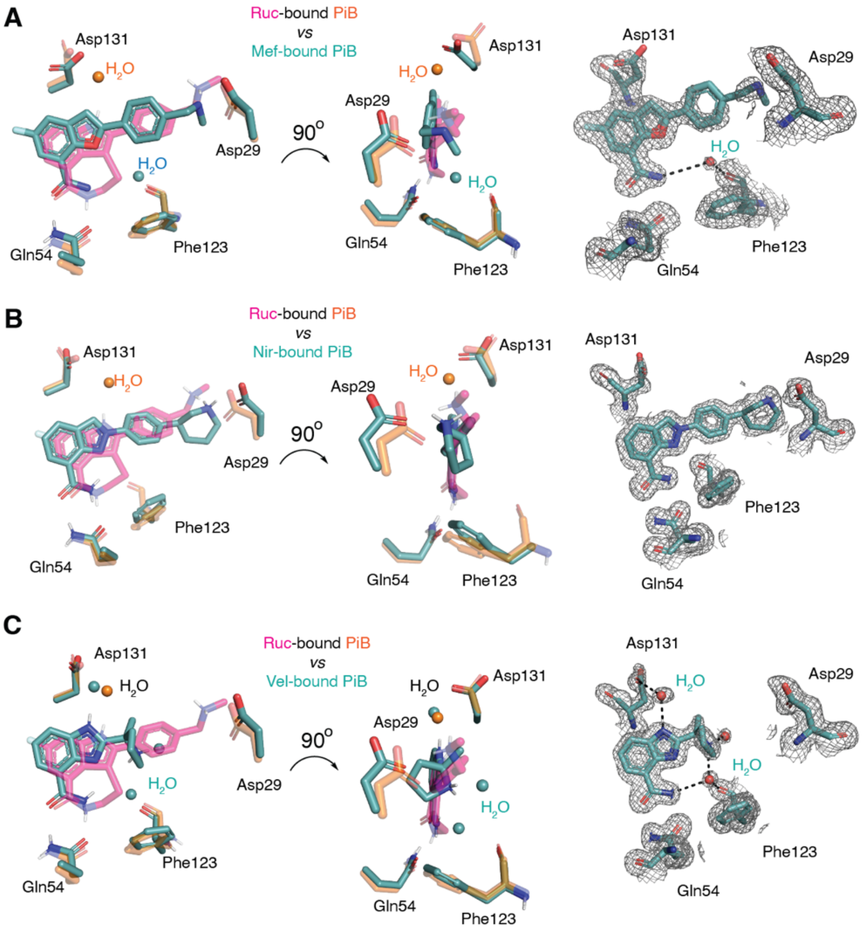


**Fig. S22. Each drug positions the bicyclic core in the same orientation, and forms key hydrogen bonds to Gln54.**

(A) Mefuparib-bound PiB' (green) vs rucaparib-bound PiB' (pink). Veliparib-bound PiB' (green) vs rucaparib-bound PiB' (pink). (C) Niraparib-bound PiB' (green) vs rucaparib-bound PiB' (pink). The 2mFo-DFc composite omit maps are contoured at 1.6 σ. The maps were generated from a model that omitted coordinates of the drugs.

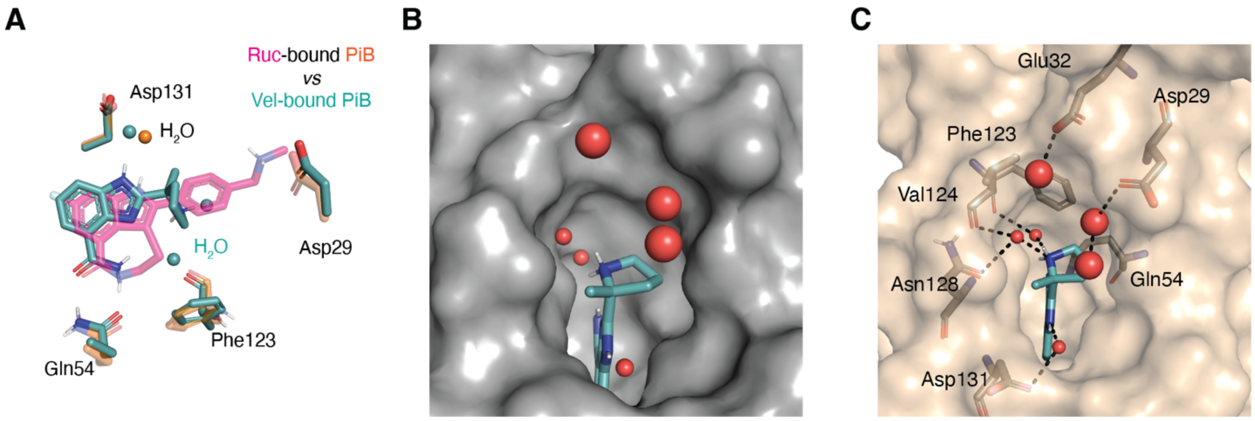


**Fig. S23. The veliparib-bound PiB’ has a water-filled cavity that is filled by a phenyl and a methylamine in rucaparib.**

(A) Overlay of veliparib-bound PiB’ interactions (protein and veliparib in green) and rucaparib-bound PiB’ interactions (protein in orange, rucaparib in purple) after superimposing onto the binding site Cα atoms. (B) The veliparib-bound PiB’ showed a large unfilled cavity that is occupied by water molecules. (C) The unfilled cavity of veliparib-bound PiB’ buried multiple waters which formed H-bonds with surrounding residues. Note that the basic pyrrolidine does not hydrogen bond directly to any amino acids. Rather, it interacts indirectly to main chain of Val124 and Asn128 via two water molecules.

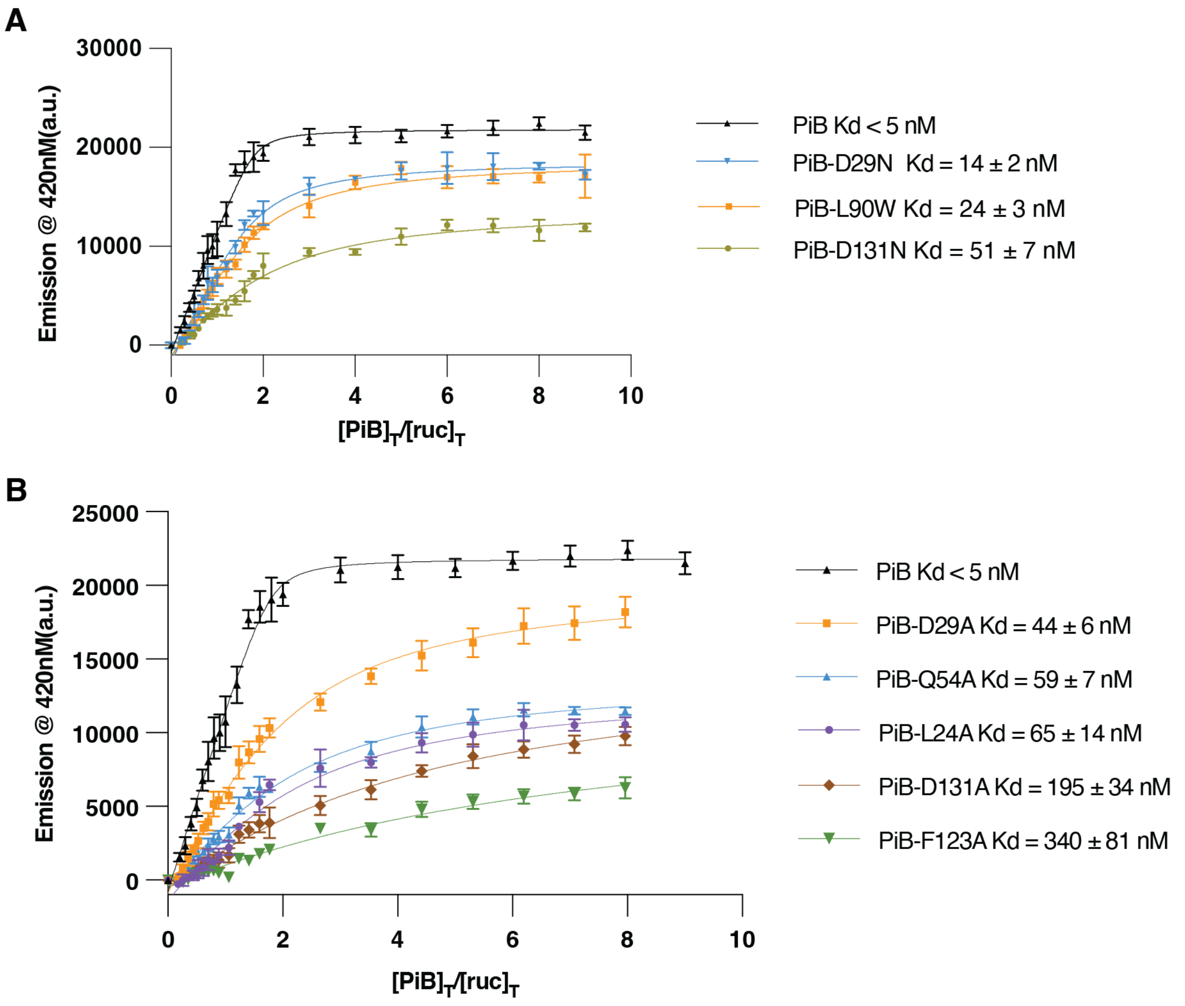


**Fig. S24. Fluorescence emission data of single-site mutants of PiB.**

(A) The measure dissociation constants ($K_D$) of rucaparib to PiB-D29N, PiB-L90W and PiB-D131N. (B) The measure dissociation constants ($K_D$) of rucaparib to PiB-D29A, PiB-Q54A, PiB-L24A, PiB-D131A and PiB-F123A. Variable protein concentrations were added into constant rucaparib (100 nM) for the binding assay in 96 well plates. Three or four replicate experiments are performed for each mutant. The fluorescence emission data was fit to a single-site binding model (solid lines). Errors are the standard deviation of the fitted parameters.

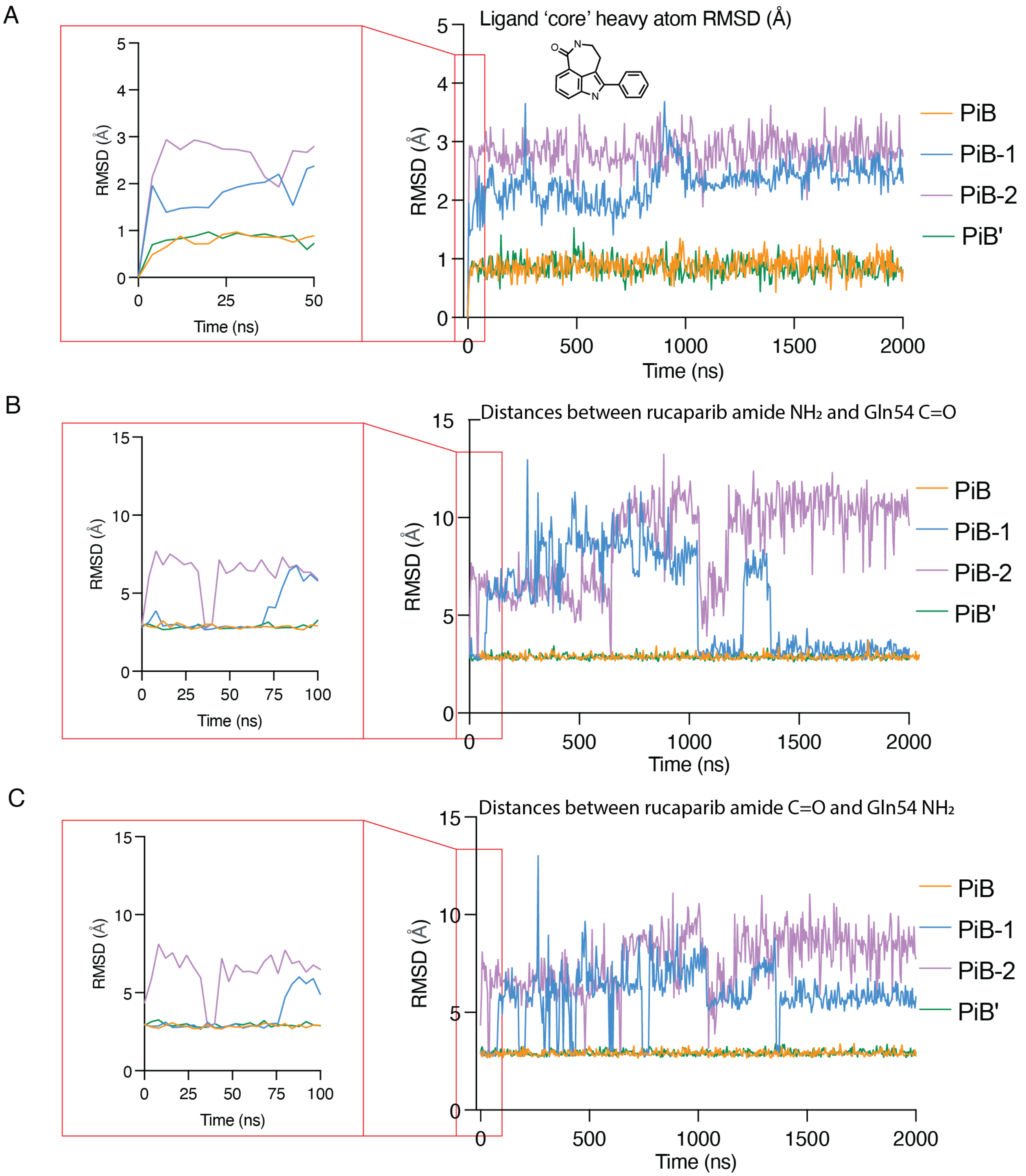


**Fig. S25. Comparison of ligand flexibility between 4 PiB designs in MD stimulations.** (A) RMSDs of the core rucaparib atoms reveal that rucaparib is conformationally most stable in PiB and PiB'. (B) (C) The stimulations showed that only PiB and PiB' consistently maintains the designed hydrogen bond interactions between the rucaparib amide group and the Gln54.

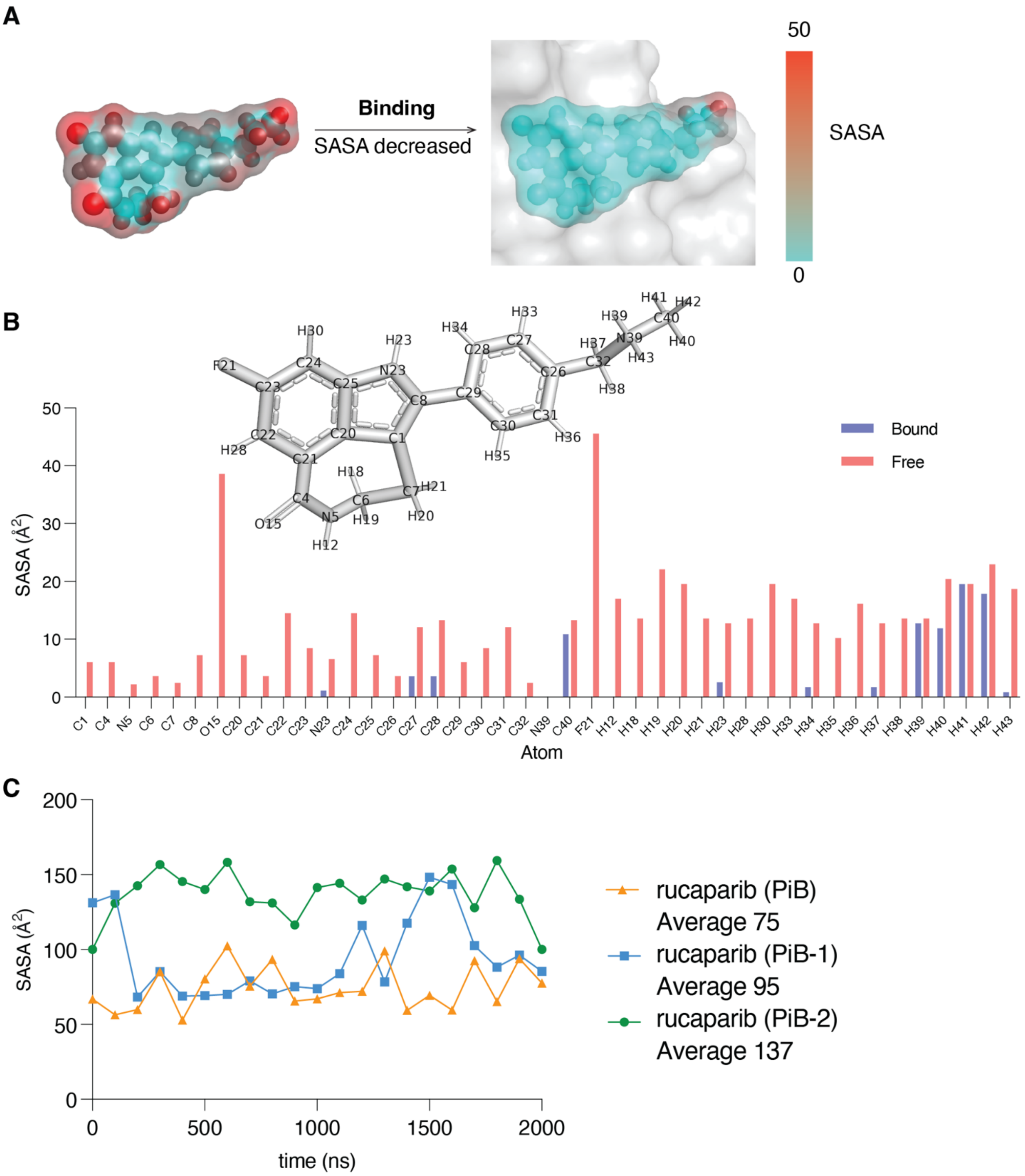


**Fig. S26. The solvent accessible surface area (SASA) of rucaparib in PiB’ and in individual frames from MD simulations.**

(A) The SASA of rucaparib decreases upon binding. (B) The SASA value of each atom of rucaparib in the free and bound forms. (C) The SASA of individual frames from MD simulations.

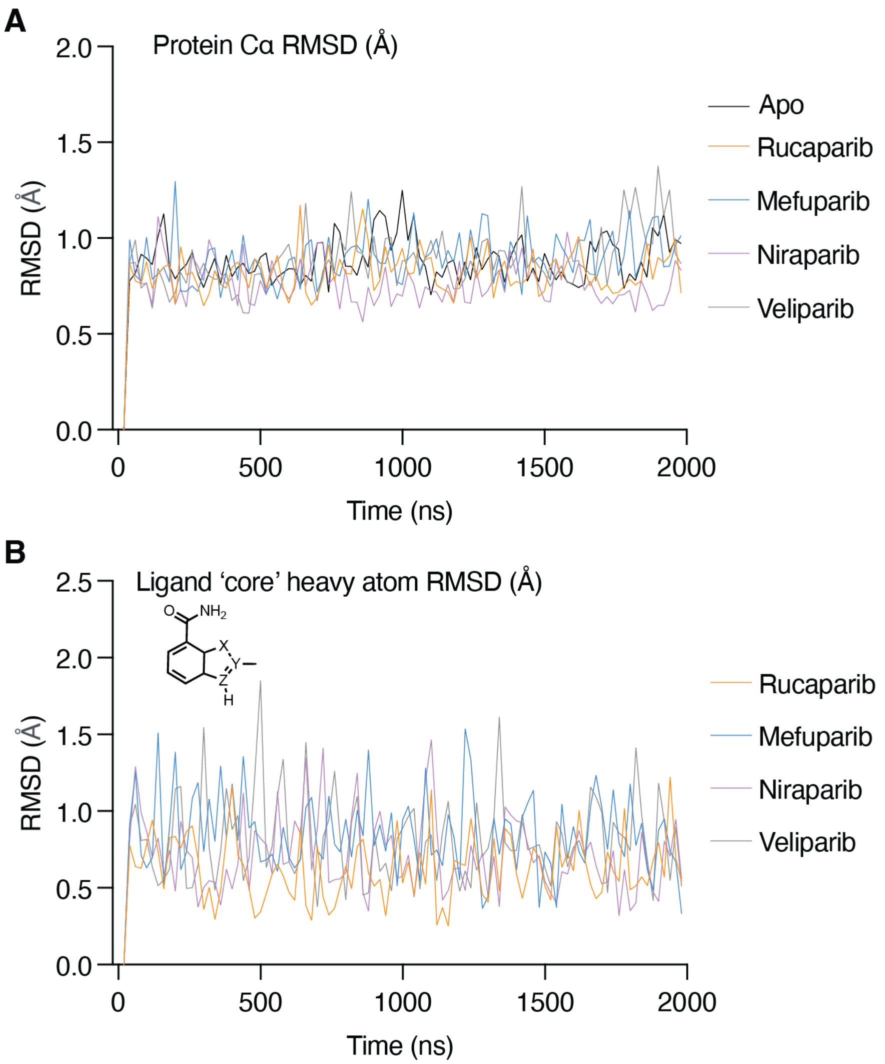


**Fig. S27. PiB and drug RMSD throughout the simulations.**
(A) RMSD of PiB C-alpha is evenly maintained and not increased over the course of the simulation, indicating overall structural stability. (B) Likewise, RMSD of ligand cores are stable throughout the simulations. In both instances, one representative trajectory of each drug complex is shown for clarity.

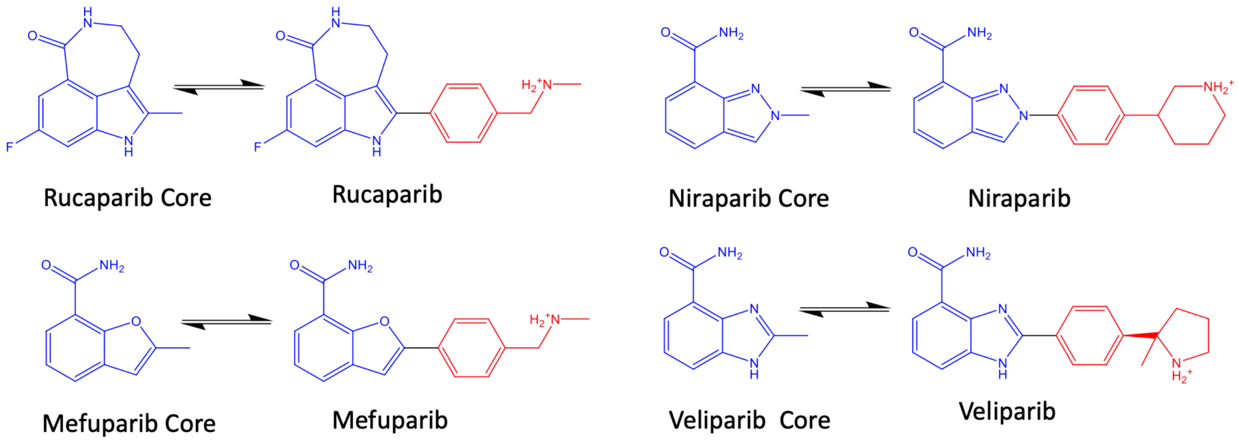


**Fig. S28. Thermodynamics pathways used for free energy calculations.**
Absolute binding free energy calculations were performed on the constrained fused ring cores (blue). Relative binding free energy calculations were used to estimate the free energy contribution of non-core regions (red) of each molecule.

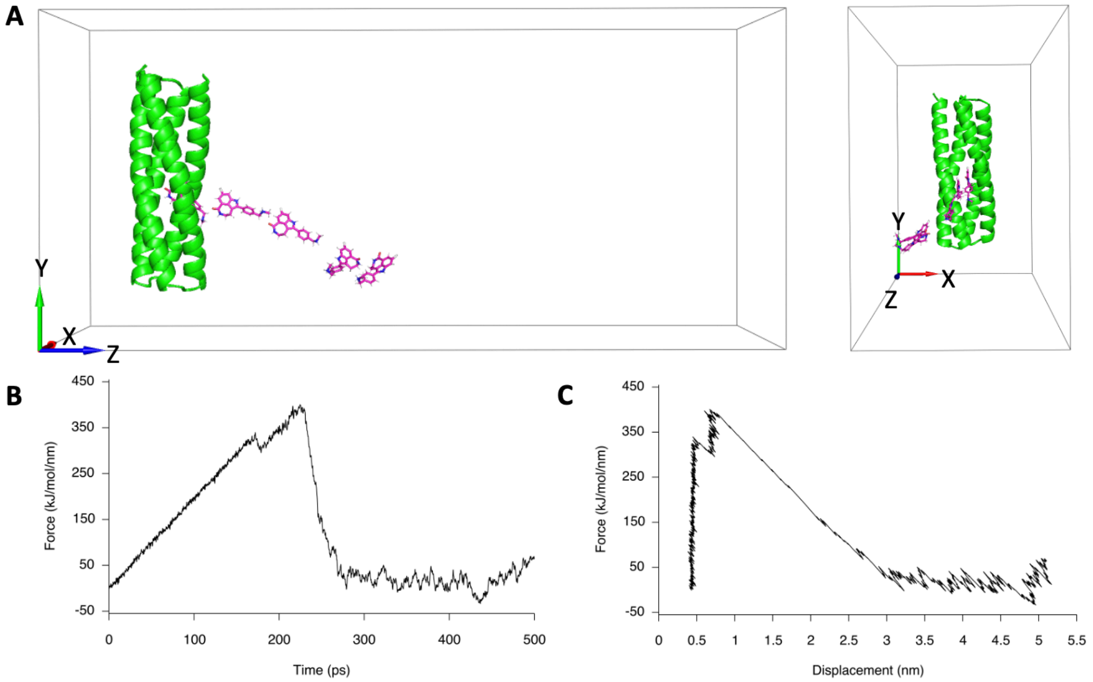


**Fig. S29. Example steered molecular dynamics simulation pulling rucaparib away from PIB.**

(A) Illustration of the pulling simulation. The pulled group (ligand) is shown in magenta sticks, and the restrained reference group (PIB) is shown in green cartoon. Besides the primary movement along the z-axis (pulling axis), the ligand's trajectory experiences orthogonal deviations due to random solvent fluctuations. (B) Plot of pull force against simulation time. (C) Plot of pull force against the displacement of the pulled group.

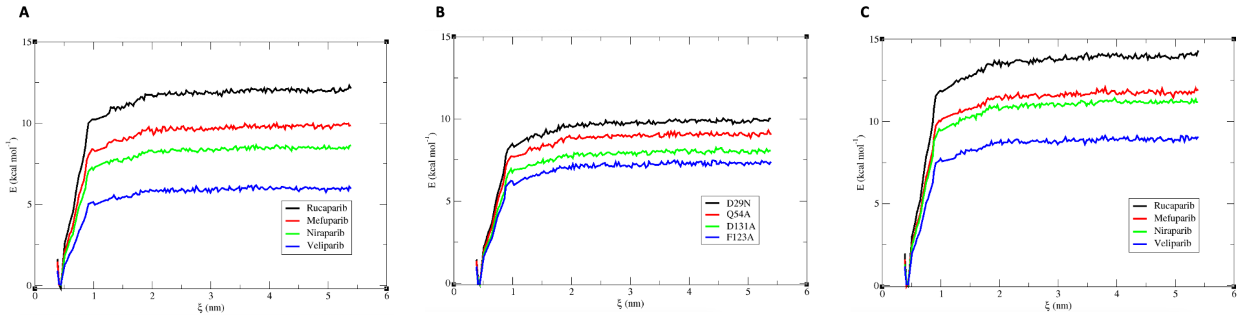


**Fig. S30. Potential of mean force (PMF) curves for systems simulated with molecular dynamics**. (A) PMF curves of PiB with PARPi analogues. (B) PMF curves of PiB mutants with Rucaparib. (C) PMF curves of PiB’ with PARPi analogues.

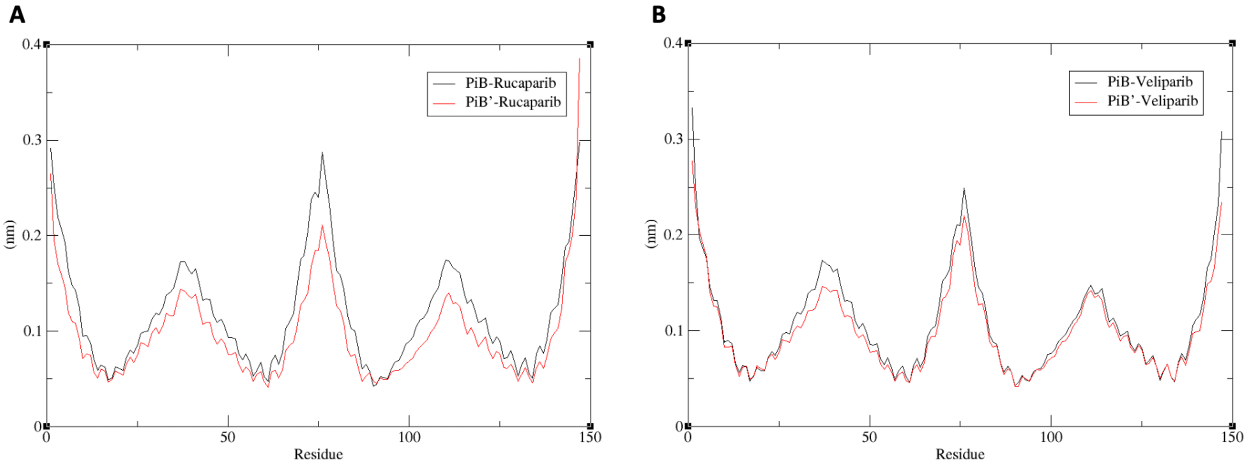


**Fig. S31. Root Mean Square Fluctuation (RMSF) of selected protein-ligand complexes.** Drug-bound PiB' systems show lower RMSF values, suggesting a stronger interaction between PiB' and the ligand.

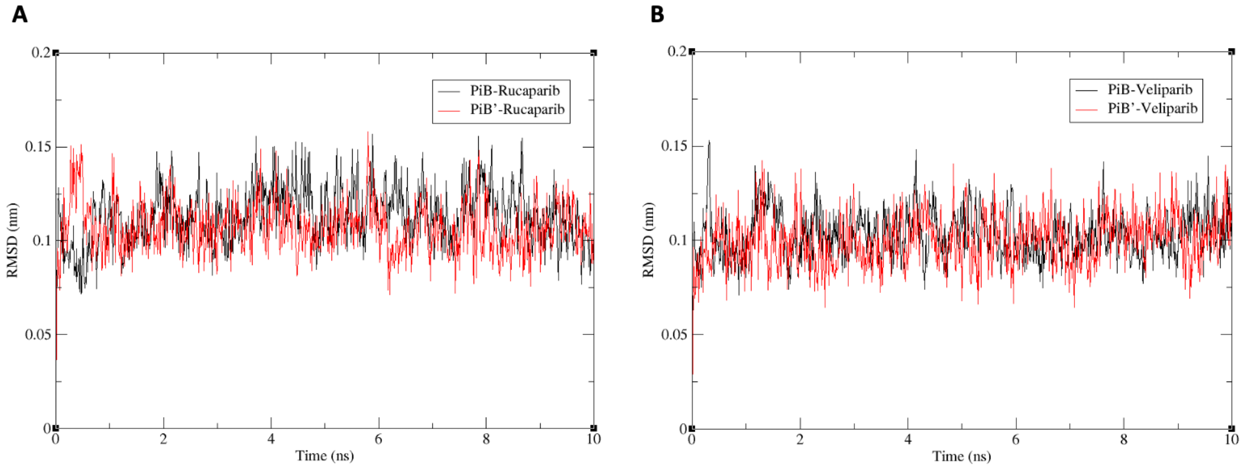


**Fig. S32. Root Mean Square Deviation (RMSD) of alpha-carbon atoms (Cα) for selected protein-ligand complexes.**

This figure illustrates a slightly lower average RMSD for PiB' in comparison to PiB. In the case of Rucaparib, the average RMSD values are 0.11 for PiB and 0.10 for PiB'. For Veliparib, the values are 0.10 for PiB and 0.09 for PiB'.

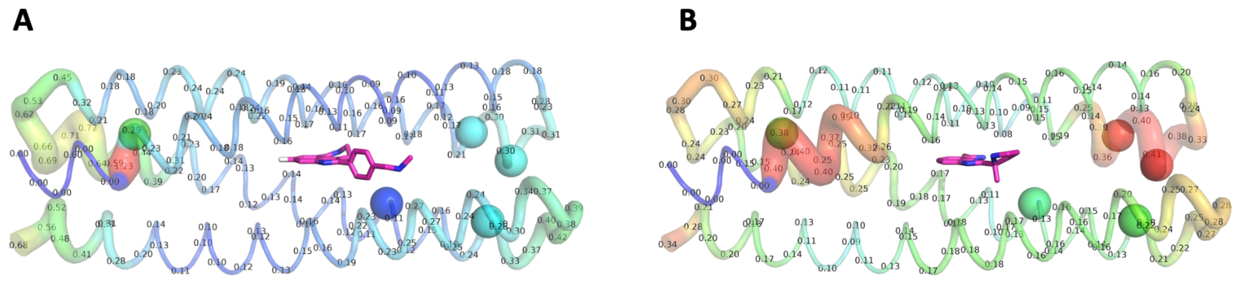


**Fig. S33. Root Mean Square Deviation (RMSD) of alpha-carbon atoms between average simulated structures of PiB and PiB' in the presence of (A) rucaparib and (B) veliparib.** After superposition of average C-alpha coordinates (averaged over the MD simulations used to compute the binding free energies) of PiB onto PiB', we computed the C-alpha RMSD between sliding windows of 7 residues, shown numerically as a label on the first residue of the sliding window (in units of Ångstroms). The sliding-window RMSD is also represented with a blue-green-red color gradient, where red corresponds to higher values of RMSD (internally normalized, so A and B are not on the same scale). The last six residues of the C-terminus are highlighted in blue and omitted in the sliding residue analysis. Spheres indicate alanine mutations unique to PiB', positioned at sites of structural difference (elevated RMSD) between the PiB and PiB' proteins in the simulations. Note that the RMSD values are very low between these sliding windows (< 0.1 – 0.7 Å), denoting overall agreement between the two proteins with little structural divergence during the simulations.

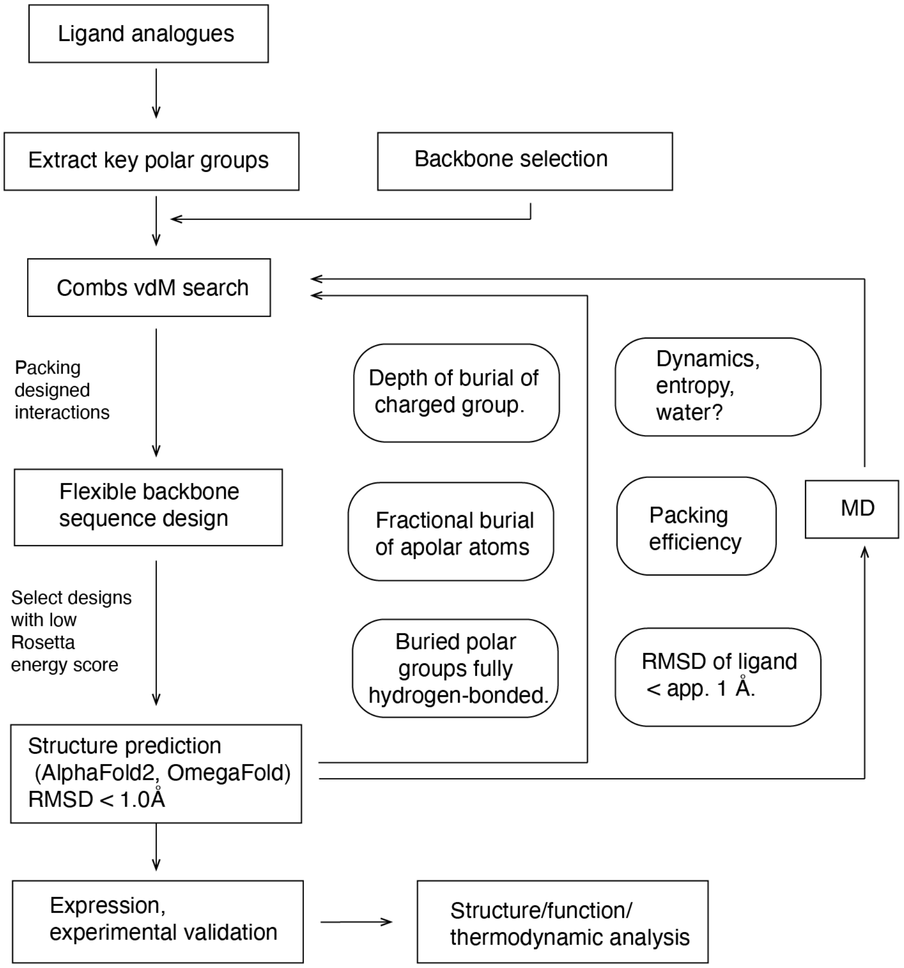


**Fig. S34. Computational design process.**

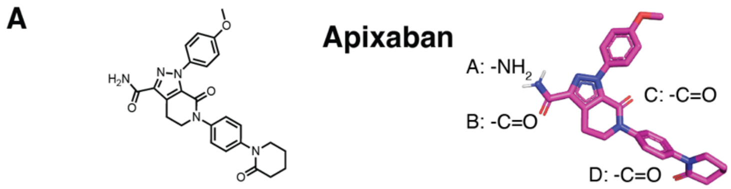


**B Hydrogen bonds in Apixaban-bound ABLE (or exposed to solvent).**

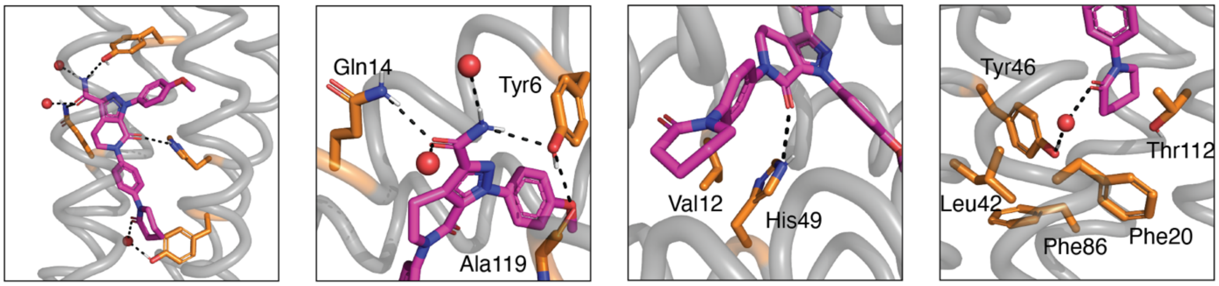


| | | | |
|---|---|---|---|
| **Hydrogen Bonds (HB) to Apixaban (Apx)** | Apx's A: -$NH_2$ to:<br>NH1: Tyr6's -OH (1 HB),<br>NH2: exposed to solvent.<br>Apx's B:-C=O to:<br>Gln14's -$NH_2$ (1 HB),<br>Exposed to solvent. | Apx's C: -C=O to:<br>His49's Imidazole NH (1 HB)<br>Lack 1 HB | Apx's D: -C=O to:<br>Tyr46's -OH (1 HB)<br>Lack 1 HB |
| **Hydrogen Bonds to first-shell amino acids** | Tyr6's -OH to:<br>Apx's A:- $NH_2$ (1HB),<br>Ala119's backbone's -C=O (1 HB).<br>Gln14's -C=O to:<br>Apx's B:-C=O (1 HB):<br>The rest exposed to solvent. | (Chain C) His49's imidazole NH to:<br>Apx's C:-C=O (1 HB)<br>Lack 1 HB | (Chain C) Tyr46's -OH to:<br>Apx's D:-C=O (1 HB)<br>Lack 1 HB |

**C Hydrogen bonds in Apixaban-bound Factor Xa (or exposed to solvent).**

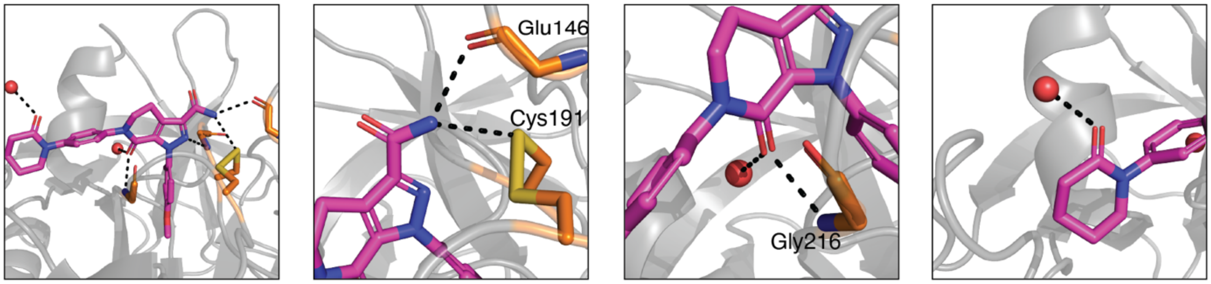


| | | | |
|---|---|---|---|
| **Hydrogen Bonds to Apixaban (Apx)** | Apx's A: -$NH_2$ to:<br>NH1: Glu146's backbone -C=O (1 HB),<br>NH2: Cys191's -S- (1 HB)<br>Apx's B:-C=O to:<br>Exposed to solvent. | Apx's C: -C=O to:<br>Gly216's backbone NH (1 HB)<br>Water (1 HB) | Apx's D: -C=O to:<br>Exposed to solvent. |
| **Hydrogen Bonds to first-shell amino acids** | Glu146's backbone -C=O to:<br>Apx's A:- $NH_2$ (1 HB),<br>Exposed to solvent.<br>Cys191's -S- to:<br>Apx's B:-C=O (1 HB) | Gly216's backbone NH to:<br>Apx's C:-C=O (1 HB) | |

**Fig. S35. The structure of apixaban in ABLE ($K_D$ 5 µM) (PDB 6w70) shows that ABLE is not as efficiently engaged in hydrogen-bond as Factor Xa ($K_D$ 0.1 nM) (PDB 2P16).**
(A) The structure and polar groups of apixaban. (B) The H-bonds in apixaban-bound ABLE (in chain C of PDB 6w70). Note that two buried chemical groups each has only one of the two possible

H-bonds satisfied. Additionally, two sidechain HB sites are not satisfied. (C) The polar interactions of apixaban to Factor Xa. All potential H-bonded groups are exposed to solvent or H-bonded.

**A** **Hydrogen bonds in rucaparib-bound PiB’ (or exposed to solvent)**

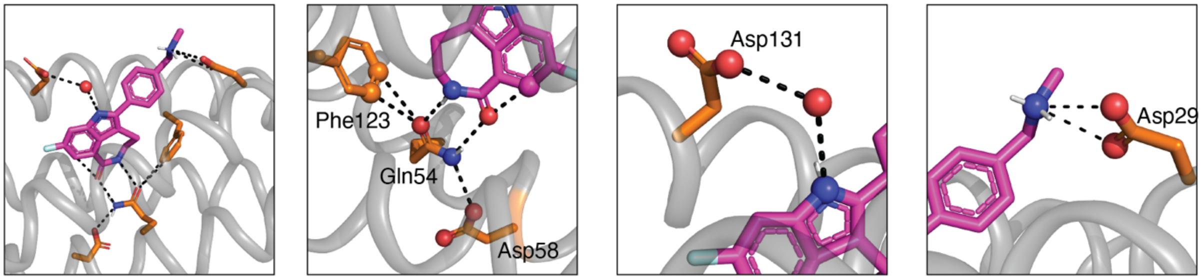


| | | | |
|---|---|---|---|
| **Hydrogen Bonds (HB) to Rucaparib (Ruc)** | Ruc’s -C=O to:<br>Gln54’s -NH (1 HB),<br>Intramolecular (arometic CH HB).<br>Ruc’s -NH- to:<br>Gln54’s -C=O (1 HB). | Ruc’s indole NH to:<br>Asp131’s $-COO^-$<br>(water-mediated HB). | Ruc’s ammonium $-(NH_2)^+$:<br>NH1: Asp29 (Direct bidentate HB)<br>NH2: exposed to solvent. |
| **Hydrogen Bonds to first-shell amino acids** | Gln54’s $-NH_2$ to:<br>Ruc’s -C=O (1 HB),<br>Asp58 $-COO^-$ (1 HB).<br>Gln54’s -C=O to:<br>ruc’s -NH- (1 HB):<br>Phe123 (2 arometic CH HB). | Asp131’s $COO^-$ to:<br>Ruc’s indole NH<br>(water-mediated HB),<br>The rest exposed to solvent. | Asp29 ’s $COO^-$ to:<br>Ruc’s ammonium $-(NH_2)^+$ (2 HB)<br>The rest exposed to solvent. |

**B** **Hydrogen bonds in rucaparib-bound PARP1 (or exposed to solvent)**

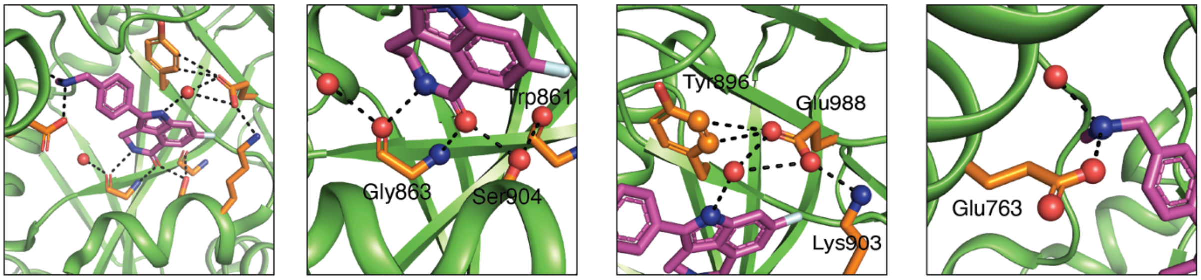


| | | | |
|---|---|---|---|
| **Hydrogen Bonds to Rucaparib (Ruc)** | Ruc’s -C=O to:<br>Gly868 backbone -NH- (1 HB),<br>Ser904 -OH (1 HB)<br>Intramolecular (arometic CH HB).<br>Ruc’s -NH- to:<br>Gly868 backbone -C=O (1 HB). | Ruc’s indole NH to:<br>Glu988 (water-mediated HB) | Ruc’s ammonium $-(NH_2)^+$:<br>NH1: Glu763 (1 HB)<br>NH2: exposed to solvent. |
| **Hydrogen Bonds to first-shell amino acids** | Gly868’s backbone -NH- to:<br>Ruc’s -C=O (1HB).<br>Gly868’s backbone -C=O to:<br>Ruc’s -NH- (1 HB):<br>Water (1 HB)<br>Ser904’s -OH to:<br>Ruc’s -C=O (1 HB),<br>Trp861 backbone -C=O (1 HB). | Glu988’s $COO^-$ to:<br>Ruc’s indole NH<br>(water-mediated HB),<br>Tyr896 (2 arometic CH HB)<br>Lys903’s $-NH_3^+$ (1 HB) | Glu763 ’s $COO^-$ to:<br>Ruc’s ammonium $-(NH_2)^+$ (1 HB)<br>The rest exposed to solvent. |

**Fig. S36. The potential H-bonding sites in rucaparib-bound PiB’ ($K_D$ < 5 nM) and PARP1 ($K_D$ 0.1 - 1 nM) (PDB 6wkk) are fully saturated, as are the first shell interacting amino acids polar groups.**

(A) The H-bonds to rucaparib in PiB’. Note that all the HB potentials of rucaparib are satisfied as the first shell interaction amino acids (Asp29, Gln54, Asp131) are either H-bonds to second shell or exposed to solvent. (B) The H-bonds to rucaparib in PARP1.

**Table S1. DNA and protein sequences used in this study.**

| | Protein sequence | DNA sequence |
|---|---|---|
| PiB_01 | MGSGHHHHHHGGSE<br>NLYFQ/SEAQELLSRL<br>ASLLETANKTAETAA<br>QVWNTAQKAYANGD<br>EEAVKSYLEELRQLQ<br>AQFDTYATQAVKLTQ<br>QVKNVNPDEEGDKT<br>YSTLVKLYKIAVEFSR<br>LLEEARQAAANGDKE<br>SYNKYLNQLRSAASA<br>GNQALTEFTKLFNTW<br>VKK* | ATGGGGTCAGGACATCACCATCACCACCAC<br>GGCGGTAGCGAGAACCTGTACTTCCAGAGC<br>GAAGCGCAAGAGCTGTTGTCTCGTCTGGCTT<br>CGCTGCTGGAAACGGCGAACAAAACCGCGG<br>AAACCGCGGCTCAAGTGTGGAATACTGCGC<br>AAAAAGCGTACGCGAATGGCGACGAGGAG<br>GCCGTCAAGTCCTATTTGGAAGAGCTGCGC<br>CAGCTTCAGGCACAATTTGATACCTACGCG<br>ACCCAGGCCGTGAAGCTGACGCAGCAGGTG<br>AAGAACGTTAATCCGGATGAGGAGGGTGAC<br>AAAACGTACTCCACCCTGGTAAAGCTCTAC<br>AAAATCGCCGTTGAGTTCAGCCGCTTGTTGG<br>AAGAGGCGCGTCAAGCTGCGGCTAACGGCG<br>ACAAAGAAAGCTACAACAAGTATCTGAATC<br>AGCTGCGTAGCGCAGCATCTGCGGGTAACC<br>AAGCGCTGACCGAGTTCACCAAATTATTTA<br>ACACCTGGGTTAAAAAGTAA |
| PiB | MGSGHHHHHHGGSE<br>NLYFQ/SDAQEILSRL<br>NSVLEAAWKTILNLA<br>SATDAAEKAYKEGRE<br>EDLKTYLDQARSYQS<br>QVDQYAVETVRLLRE<br>LKKVFPDEEADRALQ<br>IAEKLLKTVQEASKTL<br>DTAVKAARNGDEETF<br>AKAFNQFVSLGNQAD<br>TLFTQLQRTLTNLNK<br>K | ATGGGGTCAGGACATCACCACCATCACCAC<br>GGCGGCAGCGAAAACCTGTACTTCCAGTCT<br>GATGCTCAGGAGATCCTGAGCCGTTTGAAT<br>AGCGTTCTGGAAGCGGCTTGGAAAACCATC<br>TTAAACCTGGCAAGCGCAACCGATGCTGCC<br>GAGAAGGCGTATAAAGAAGGTCGTGAAGA<br>GGACTTGAAGACGTACCTGGACCAAGCACG<br>CTCCTACCAGTCGCAGGTTGATCAATATGCG<br>GTAGAGACCGTTCGTCTGCTGCGTGAGTTG<br>AAGAAGGTGTTCCCGGATGAAGAGGCCGAC<br>CGCGCACTGCAAATTGCGGAAAAATTGCTT<br>AAGACCGTGCAGGAGGCCTCCAAAACCCTC<br>GACACTGCGGTGAAAGCGGCTCGTAATGGT<br>GATGAAGAGACGTTTGCGAAAGCGTTTAAC<br>CAGTTTGTCAGCCTGGGTAATCAAGCGGAC<br>ACTCTGTTCACCCAGCTGCAACGCACCCTGA<br>CCAACCTAAACAAAAAGTAA |
| PiB_02 | MGSGHHHHHHGGSE<br>NLYFQ/SRAQELLSRA<br>AQVLTSLAKTIEQAA<br>QTFTALVRALRNGD<br>WDSAKSYTEQLVQLQ<br>KQADSLARELVNLFR<br>EVAKVNPDEEGEKLL | ATGGGGTCAGGACATCACCACCATCACCAC<br>GGCGGTAGCGAGAACCTGTATTTCCAGAGC<br>CGCGCTCAAGAACTGTTATCTCGCGCGGCG<br>CAGGTTCTGACCAGCTTGGCGAAGACCATT<br>GAACAAGCTGCGCAAACGTTTACCGCGTTG<br>GTCAGAGCCCTGCGTAATGGCGACTGGGAT<br>AGCGCAAAATCCTATACCGAGCAGTTGGTT |

| | | |
|---|---|---|
| | QTAEALLKATETLSKI<br>LSEAKKAADNGDED<br>KLNTALEQLKQAGEQ<br>ANTYFTELVKLFNTY<br>VSK* | CAGCTGCAGAAACAGGCTGACTCGCTGGCG<br>CGTGAGCTGGTGAACTTGTTTCGTGAAGTTG<br>CAAAGGTGAACCCGGATGAGGAAGGTGAA<br>AAATTGCTGCAAACTGCGGAAGCGCTGCTT<br>AAGGCTACCGAAACCCTGTCTAAGATCCTG<br>TCCGAGGCCAAGAAAGCAGCCGACAACGGC<br>GACGAGGATAAACTGAATACCGCATTGGAA<br>CAACTGAAGCAGGCGGGTGAGCAAGCGAAT<br>ACGTACTTCACCGAGCTCGTAAAACTGTTCA<br>ACACATACGTGAGCAAATAA |
| PiB’ | MGSGHHHHHHGGSE<br>NLYFQ/SDAQEILSRL<br>NSVLEAAWKTILNLA<br>SATDAAEKAYKEGRE<br>EDLATYLDQAASYQS<br>QVDQYAVETVRLLAE<br>LKKVFPDEEADRALQ<br>IAEKLLKTVQEASKTL<br>DTAVAAAANGDEETF<br>AKAFNQFVSLGNQAD<br>TLFTQLQRTLTNLNK<br>K* | ATGGGGTCAGGACATCACCACCATCACCAC<br>GGCGGCAGCGAAAACCTGTACTTCCAGTCT<br>GATGCTCAGGAGATCCTGAGCCGTTTGAAT<br>AGCGTTCTGGAAGCGGCTTGGAAAACCATC<br>TTAAACCTGGCAAGCGCAACCGATGCTGCC<br>GAGAAGGCGTATAAAGAAGGTCGTGAAGA<br>GGACTTGGCGACGTACCTGGACCAAGCAGC<br>CTCCTACCAGTCGCAGGTTGATCAATATGCG<br>GTAGAGACCGTTCGTCTGCTGGCTGAGTTG<br>AAGAAGGTGTTCCCGGATGAAGAGGCCGAC<br>CGCGCACTGCAAATTGCGGAAAAATTGCTT<br>AAGACCGTGCAGGAGGCCTCCAAAACCCTC<br>GACACTGCGGTGGCAGCGGCTGCTAATGGT<br>GATGAAGAGACGTTTGCGAAAGCGTTTAAC<br>CAGTTTGTCAGCCTGGGTAATCAAGCGGAC<br>ACTCTGTTCACCCAGCTGCAACGCACCCTGA<br>CCAACCTAAACAAAAAGTAA |

**Table S2. Crystallographic properties, crystallization conditions, data collection, and model refinement statistics for drug-free and drug-bound PiB'.**

| | 8TN1 | 8TN6 | 8TNB | 8TNC | 8TND |
|---|---|---|---|---|---|
| Ligand | - | rucaparib | mefuparib | niraparib | veliparib |
| PDB ID | 8TN1 | 8TN6 | 8TNB | 8TNC | 8TND |
| space group | *P6122* | *P6122* | *P6122* | *P6122* | *P6122* |
| *a, b, c* (Å) | 90.52 90.52 202.33 | 91.25 91.25 201.95 | 90.21 90.21 202.11 | 91.01 91.01 203.65 | 90.75 90.75 202.35 |
| α, β, γ (˚) | 90.0, 90.0, 120.0 | 90.0, 90.0, 120.0 | 90.0, 90.0, 120.0 | 90.0, 90.0, 120.0 | 90.0, 90.0, 120.0 |
| protein per asymmetric unit | 3 | 3 | 3 | 3 | 3 |
| crystallization conditions | 2M Ammonium Sulfate | 2M Ammonium Sulfate | 2M Ammonium Sulfate | 2M Ammonium Sulfate | 2M Ammonium Sulfate |
| **Data collection** | | | | | |
| wavelength (Å) | 1.11583 Å (11111 eV) | 1.11583 Å (11111 eV) | 1.11583 Å (11111 eV) | 1.11583 Å (11111 eV) | 1.11583 Å (11111 eV) |
| resolution (Å) | 202.33-1.61 (1.64-1.61) | 201.95-1.36 (1.38-1.36) | 202.11-1.39 (1.41-1.39) | 203.65-1.39 (1.42-1.39) | 202.35–1.29 (1.31-1.29) |
| total reflections | 61435 (728) | 107193 (4918) | 97882 (4412) | 99665 (4668) | 123927 (5369) |
| Multiplicity | 55.7 (11.8) | 74.5 (66.4) | 74.2 (60.1) | 37.0 (36.1) | 67.6 (24.5) |
| completeness (%) | 95.2 (22.7) | 99.7 (93.5) | 99.3 (92.2) | 99.8 (96.6) | 99.4 (88.7) |
| mean I/σ | 26.8 (0.3) | 38.1 (1.4) | 24.9 (0.2) | 17.0 (0.6) | 31.2 (0.8) |
| $R_{merge}$ | 0.116 (5.957) | 0.082 (4.049) | 0.138 (30.827) | 0.169 (6.178) | 0.090 (3.378) |
| $R_{measure}$ | 0.117 (6.220) | 0.083 (4.110) | 0.139 (31.085) | 0.172 (6.266) | 0.091 (3.448) |
| $CC_{1/2}$ | 1.0 (0.092) | 1.0 (0.468) | 1.0 (0.005) | 1.0 (0.212) | 1.0 (0.363) |
| **Refinement** | | | | | |
| $R_{work}$ | 0.2128 | 0.1952 | 0.2181 | 0.2020 | 0.2022 |
| $R_{free}$ | 0.2497 | 0.2169 | 0.2367 | 0.2262 | 0.2236 |

| | | | | | |
|---|---|---|---|---|---|
| $RMS_{bonds}$ | 0.0072 | 0.0073 | 0.0070 | 0.0058 | 0.0056 |
| $RMS_{angles}$ | 0.87 | 0.99 | 0.99 | 0.79 | 0.79 |
| Ramachandran allowed (%) | 98.16 | 99.31 | 98.62 | 98.62 | 98.85 |
| Ramachandran outliers (%) | 0.46 | 0.28 | 0.84 | 0.84 | 0.28 |
| clashscore | 4.53 | 4.46 | 6.82 | 2.59 | 3.31 |
| average B-factor | 33.22 | 29.70 | 35.42 | 29.11 | 25.75 |

#### Table S3. Binding site Cα RMSD.

| | apo_A | apo_B | apo_C | ruc_A | ruc_B | ruc_C | mef_A | mef_B | mef_C | nir_A | nir_B | nir_C | vel_A | vel_B | vel_C | Design | AF2 |
|---|---|---|---|---|---|---|---|---|---|---|---|---|---|---|---|---|---|
| apo_A | 0 | 0.25 | 0.42 | 0.23 | 0.28 | 0.41 | 0.25 | 0.21 | 0.36 | 0.32 | 0.33 | 0.4 | 0.18 | 0.27 | 0.42 | 0.47 | 0.25 |
| apo_B | 0.25 | 0 | 0.27 | 0.33 | 0.27 | 0.35 | 0.22 | 0.29 | 0.24 | 0.3 | 0.35 | 0.33 | 0.26 | 0.2 | 0.29 | 0.48 | 0.28 |
| apo_C | 0.42 | 0.27 | 0 | 0.48 | 0.4 | 0.37 | 0.42 | 0.46 | 0.26 | 0.46 | 0.49 | 0.39 | 0.45 | 0.41 | 0.31 | 0.63 | 0.36 |
| ruc_A | 0.23 | 0.33 | 0.48 | 0 | 0.15 | 0.29 | 0.26 | 0.22 | 0.35 | 0.29 | 0.29 | 0.32 | 0.22 | 0.29 | 0.39 | 0.38 | 0.28 |
| ruc_B | 0.28 | 0.27 | 0.4 | 0.15 | 0 | 0.21 | 0.26 | 0.26 | 0.28 | 0.28 | 0.3 | 0.28 | 0.26 | 0.27 | 0.32 | 0.39 | 0.27 |
| ruc_C | 0.41 | 0.35 | 0.37 | 0.29 | 0.21 | 0 | 0.35 | 0.37 | 0.23 | 0.34 | 0.36 | 0.2 | 0.36 | 0.35 | 0.22 | 0.46 | 0.35 |
| mef_A | 0.25 | 0.22 | 0.42 | 0.26 | 0.26 | 0.35 | 0 | 0.14 | 0.26 | 0.15 | 0.21 | 0.29 | 0.12 | 0.08 | 0.3 | 0.42 | 0.31 |
| mef_B | 0.21 | 0.29 | 0.46 | 0.22 | 0.26 | 0.37 | 0.14 | 0 | 0.31 | 0.18 | 0.17 | 0.32 | 0.09 | 0.19 | 0.36 | 0.41 | 0.29 |
| mef_C | 0.36 | 0.24 | 0.26 | 0.35 | 0.28 | 0.23 | 0.26 | 0.31 | 0 | 0.29 | 0.33 | 0.18 | 0.31 | 0.26 | 0.12 | 0.52 | 0.34 |
| nir_A | 0.32 | 0.3 | 0.46 | 0.29 | 0.28 | 0.34 | 0.15 | 0.18 | 0.29 | 0 | 0.1 | 0.25 | 0.18 | 0.17 | 0.3 | 0.39 | 0.34 |
| nir_B | 0.33 | 0.35 | 0.49 | 0.29 | 0.3 | 0.36 | 0.21 | 0.17 | 0.33 | 0.1 | 0 | 0.29 | 0.2 | 0.23 | 0.35 | 0.37 | 0.32 |
| nir_C | 0.4 | 0.33 | 0.39 | 0.32 | 0.28 | 0.2 | 0.29 | 0.32 | 0.18 | 0.25 | 0.29 | 0 | 0.31 | 0.28 | 0.16 | 0.46 | 0.38 |
| vel_A | 0.18 | 0.26 | 0.45 | 0.22 | 0.26 | 0.36 | 0.12 | 0.09 | 0.31 | 0.18 | 0.2 | 0.31 | 0 | 0.15 | 0.34 | 0.41 | 0.29 |
| vel_B | 0.27 | 0.2 | 0.41 | 0.29 | 0.27 | 0.35 | 0.08 | 0.19 | 0.26 | 0.17 | 0.23 | 0.28 | 0.15 | 0 | 0.28 | 0.41 | 0.33 |
| vel_C | 0.42 | 0.29 | 0.31 | 0.39 | 0.32 | 0.22 | 0.3 | 0.36 | 0.12 | 0.3 | 0.35 | 0.16 | 0.34 | 0.28 | 0 | 0.52 | 0.4 |
| Design | 0.47 | 0.48 | 0.63 | 0.38 | 0.39 | 0.46 | 0.42 | 0.41 | 0.52 | 0.39 | 0.37 | 0.46 | 0.41 | 0.41 | 0.52 | 0 | 0.41 |
| AF2 | 0.25 | 0.28 | 0.36 | 0.28 | 0.27 | 0.35 | 0.31 | 0.29 | 0.34 | 0.34 | 0.32 | 0.38 | 0.29 | 0.33 | 0.4 | 0.41 | 0 |

*apo-A: apo structure of PiB’ chain A. Similarly for all others.
*AF2: AlphaFold2 model.

**Table S4. Binding free energies of PiB with PARPi analogues using biased simulations in AToM-OpenMM and GROMACS.**

| Ligand | Ligand G (kcal/mol) | | Experimental G (kcal/mol) at 298 K |
|---|---|---|---|
| | ATоM-OpenMM | GROMACS | |
| Mefuparib | -15.16 | -9.96 | -9.18 |
| Niraparib | -14.20 | -8.62 | -8.45 |
| Rucaparib | -20.07 | -12.22 | -11.82 |
| Veliparib | -11.77 | -6.06 | -7.98 |

**Table S5. Calculated binding free energies of PiB, PiB' and mutated PiB with PARPi analogues derived from original and duplicate steered molecular dynamics trials.**

| Protein | Ligand | ΔGBind-Run1 (kcal/mol) | ΔGBind-Run2 (kcal/mol) | Avg (kcal/mol) | Std Dev (kcal/mol) |
|---|---|---|---|---|---|
| PiB | Rucaparib | -12.22 | -12.46 | -12.34 | 0.17 |
| PiB | Mefuparib | -9.96 | -9.55 | -9.76 | 0.29 |
| PiB | Niraparib | -8.62 | -8.92 | -8.77 | 0.21 |
| PiB | Veliparib | -6.06 | -6.36 | -6.21 | 0.21 |
| PiB' | Rucaparib | -14.18 | -13.86 | -14.02 | 0.23 |
| PiB' | Mefuparib | -11.95 | -11.68 | -11.82 | 0.19 |
| PiB' | Niraparib | -11.34 | -11.01 | -11.18 | 0.23 |
| PiB' | Veliparib | -9.07 | -9.43 | -9.25 | 0.25 |
| PiB-D29N | Rucaparib | -10.04 | -10.34 | -10.19 | 0.21 |
| PiB-Q54A | Rucaparib | -9.21 | -9.01 | -9.11 | 0.14 |
| PiB-D131A | Rucaparib | -8.15 | -7.96 | -8.056 | 0.13 |
| PiB-F123A | Rucaparib | -7.40 | -7.68 | -7.54 | 0.21 |

#### Supplementary Text

##### Rosetta design scripts and settings

**RosettaScript (rscript_dock_fixbb_design.xml)**

```
<ROSETTASCRIPTS>
	<SCOREFXNS>
		<ScoreFunction name="ligand_soft_rep" weights="ligand_soft_rep" />
		<ScoreFunction name="hard_rep" weights="ligandprime" />
		<ScoreFunction name="ref15" weights="ref2015">
			<Reweight scoretype="atom_pair_constraint" weight="5"/>
		</ScoreFunction>
		<ScoreFunction name="ref15_1" weights="ref2015">
			<Reweight scoretype="aa_composition" weight="1" />
			<Reweight scoretype="netcharge" weight="1.0" />
			<Reweight scoretype="atom_pair_constraint" weight="5"/>
			<Set aa_composition_setup_file="helix_composition.comp" />
			<Set netcharge_setup_file="helix_netcharge.charge" />
		</ScoreFunction>
	</SCOREFXNS>
	<RESIDUE_SELECTORS>
	</RESIDUE_SELECTORS>
	<TASKOPERATIONS>
		<InitializeFromCommandline name="ifcl"/>
		<ReadResfile name="resfile" filename="resfile_topology.txt"/>
		<ExtraRotamersGeneric name="extrachi" ex1="1" ex2="1"
ex1_sample_level="1" ex2_sample_level="1" extrachi_cutoff="14"/>
		<DetectProteinLigandInterface name="design_interface" cut1="6.0" cut2="8.0"
cut3="10.0" cut4="12.0" design="1" resfile="resfile_topology.txt" />
	</TASKOPERATIONS>
	<FILTERS>
		<PackStat name="pstat" confidence="0" threshold="0" repeats="10"/>
		<PackStat name="pstat_mc" threshold="0" repeats="10"/>
		<NetCharge name="net_charge" confidence="0"/>
		<LigInterfaceEnergy name="ligen"  scorefxn="ref15" include_cstE="0"
energy_cutoff="0" confidence="0"/>
		<ScoreType name="total_score_1" scorefxn="ref15_1" score_type="total_score"
threshold="0"/>
		<ScoreType name="total_score" scorefxn="ref15" score_type="total_score"
threshold="0"/>
		<BuriedUnsatHbonds name="bu" report_sc_heavy_atom_unsats="true"
scorefxn="ref15" cutoff="0" residue_surface_cutoff="20.0" ignore_surface_res="true"
print_out_info_to_pdb="true" confidence="0" />
	</FILTERS>
	<LIGAND_AREAS>
		<LigandArea name="docking_sidechain" chain="X" cutoff="6.0"
add_nbr_radius="true" all_atom_mode="true" minimize_ligand="10" />
```

```
		<LigandArea name="final_sidechain" chain="X" cutoff="6.0"
add_nbr_radius="true" all_atom_mode="true" />
		<LigandArea name="final_backbone" chain="X" cutoff="7.0"
add_nbr_radius="false" all_atom_mode="true" Calpha_restraints="0.3" />
	</LIGAND_AREAS>
	<INTERFACE_BUILDERS>
		<InterfaceBuilder name="side_chain_for_docking"
ligand_areas="docking_sidechain" />
		<InterfaceBuilder name="side_chain_for_final" ligand_areas="final_sidechain"
/>
		<InterfaceBuilder name="backbone" ligand_areas="final_backbone"
extension_window="3" />
	</INTERFACE_BUILDERS>
	<MOVEMAP_BUILDERS>
		<MoveMapBuilder name="docking" sc_interface="side_chain_for_docking"
minimize_water="true" />
		<MoveMapBuilder name="final" sc_interface="side_chain_for_final"
bb_interface="backbone" minimize_water="true" />
	</MOVEMAP_BUILDERS>
	<SCORINGGRIDS ligand_chain="X" width="46">
		<ClassicGrid grid_name="vdw" weight="1.0" />
	</SCORINGGRIDS>
	<MOVERS>
		<ConstraintSetMover name="atomic" cst_file="constraints_helix.cst" />
		<Transform name="transform" chain="X" box_size="10.0" move_distance="0.1"
angle="5" cycles="500" repeats="1" temperature="5" rmsd="4.0" use_constraints="true"
cst_fa_file="constraints_helix.cst" cst_fa_weight="5" />
		<PackRotamersMover name="design_interface" scorefxn="hard_rep"
task_operations="design_interface" />
		<InterfaceScoreCalculator name="add_scores" chains="X" scorefxn="hard_rep"
/>
		<ParsedProtocol name="low_res_dock">
			<Add mover_name="transform" />
		</ParsedProtocol>
		<PackRotamersMover name="fixbb_aro" scorefxn="ref15_1"
task_operations="resfile,extrachi,ifcl"/>
		<PackRotamersMover name="pack" scorefxn="ref15_1"
task_operations="ifcl,resfile,extrachi"/>
		<PackRotamersMover name="pack_fast" scorefxn="ref15_1"
task_operations="ifcl,resfile"/>
		<MinMover name="min_bb" scorefxn="ref15" tolerance="0.0000001"
max_iter="1000" chi="false" bb="true">
			<MoveMap name="map_bb">
				<Span begin="1" end="200" bb="true" chi="false" />
				<Span begin="201" end="999" bb="false" chi="false"/>
			</MoveMap>
```

```
			</MinMover>
			<Idealize name="idealize"/>
			<MinMover name="min_sc" scorefxn="ref15" tolerance="0.0000001"
max_iter="1000" chi="true" bb="false">
				<MoveMap name="map_sc">
					<Span begin="1" end="200" bb="false" chi="true" />
					<Span begin="201" end="999" bb="false" chi="false"/>
				</MoveMap>
			</MinMover>
			<MinMover name="min_sc_bb" scorefxn="ref15" tolerance="0.0000001"
max_iter="1000" chi="true" bb="true">
				<MoveMap name="map_sc_bb">
					<Span begin="1" end="200" bb="true" chi="true" />
					<Span begin="201" end="999" bb="false" chi="false"/>
				</MoveMap>
			</MinMover>
			<ParsedProtocol name="parsed_pack_fast" >
				<Add mover_name="pack_fast"/>
				<Add mover_name="min_bb"/>
			</ParsedProtocol>
			<ParsedProtocol name="parsed_pack" >
				<Add mover_name="pack"/>
				<Add mover_name="min_bb"/>
				<Add mover_name="min_sc"/>
			</ParsedProtocol>
			<GenericMonteCarlo name="pack_mc" preapply="0" trials="3"
temperature="0.03" filter_name="pstat_mc" sample_type="high" mover_name="parsed_pack">
					<Filters>
						<AND filter_name="total_score_1" temperature="15"
sample_type="low"/>
					</Filters>
			</GenericMonteCarlo>
			<GenericMonteCarlo name="pack_fast_mc" preapply="0" trials="2"
temperature="0.03" filter_name="pstat_mc" sample_type="high"
mover_name="parsed_pack_fast">
					<Filters>
						<AND filter_name="total_score_1" temperature="15"
sample_type="low"/>
					</Filters>
			</GenericMonteCarlo>
		</MOVERS>
		<PROTOCOLS>
			<Add mover="atomic"/>
			<Add mover_name="fixbb_aro"/>
			<Add mover_name="low_res_dock" />
			<Add mover_name="design_interface" />
```

```
			<Add mover_name="parsed_pack_fast"/>
			<Add mover_name="pack_fast_mc"/>
			<Add mover_name="pack_mc"/>
			<Add mover_name="min_sc_bb"/>
			<Add mover_name="add_scores" />
			<Add filter_name="pstat"/>
			<Add filter_name="net_charge"/>
			<Add filter_name="ligen"/>
			<Add filter_name="total_score"/>
			<Add filter_name="bu"/>
	</PROTOCOLS>
	<OUTPUT scorefxn="ref15_1"/>
</ROSETTASCRIPTS>
```

**Content of flag file (option_helix.txt)**

```
-out:path:pdb output
-out:path:score output
-extra_res_fa RUC.params
-packing:multi_cool_annealer 10
-packing:linmem_ig 10
-ignore_zero_occupancy false
-s bb_prep.pdb
```

**Command line Inputs**

```
~/rosetta_bin_linux_2020.08.61146_bundle/main/source/bin/rosetta_scripts.static.linuxgccreleas
e -database ~/rosetta_bin_linux_2020.08.61146_bundle/main/database/ -parser:protocol
rscript_dock_fixbb_design.xml @options_helix.txt
```

**Content of residue file (resfile_topology.txt)**

```
NATRO
ALLAAxc
NOTAA MTC
USE_INPUT_SC
start

1 A PIKAA TNVSDEQRKAG
2 A PIKAA TNVSDEQRKA
3 A PIKAA FTNIVWSLYDEQRKA
4 A PIKAA TNVSQA
5 A PIKAA TNVSDEQA
6 A PIKAA FTNIVWSLYDEQRKA
7 A PIKAA FTIVWSLYAG
8 A PIKAA TNVSQA
9 A PIKAA TNVSQRKA
10 A PIKAA FTIVWSLYAG
```

11 A PIKAA TNVSQA
12 A PIKAA TNVSQA
13 A PIKAA FTNIVWSLYDEQRKA
14 A PIKAA FTIVWSLYAG
15 A PIKAA TNVSDEQA
16 A PIKAA TNVSQA
17 A PIKAA FTIVWSLYAG
18 A PIKAA TNVSQA
19 A PIKAA TNVSQRKA
20 A PIKAA TNVSQRKA
21 A PIKAA FTIVWSLYAG
22 A PIKAA TNVSDEQA
23 A PIKAA TNVSQA
24 A PIKAA FTIVWSLYAG
25 A PIKAA TNVSDEQA
26 A PIKAA TNVSQA
27 A PIKAA TNVSQRKA
28 A PIKAA FTIVWSLYAG
29 A PIKAA TNVSQA
30 A PIKAA TNVSQA
31 A PIKAA FTIVWSLYAG
32 A PIKAA TNVSDEQA
33 A PIKAA TNVSDEQRKA
34 A PIKAA TNVSDEQRKA
35 A PIKAA ALICQY
36 A PIKAA DRNKAL
37 A PIKAA QARENL
38 A PIKAA G
39 A PIKAA HDNKR
40 A PIKAA LEYFVW
41 A PIKAA EDA
42 A PIKAA TNVSDEQRKA
43 A PIKAA FTIVWSLYAG
44 A PIKAA TNVSQRKA
45 A PIKAA TNVSQA
46 A PIKAA FTIVWSLYAG
47 A PIKAA FTIVWSLYAG
48 A PIKAA TNVSDEQA
49 A PIKAA TNVSDEQA
50 A PIKAA FTIVWSLYAG
51 A PIKAA TNVSQRKA
52 A PIKAA TNVSQA
53 A PIKAA FTIVWSLYAG
54 A NATRO
55 A PIKAA TNVSQRKA
56 A PIKAA TNVSDEQA

57 A PIKAA FTIVWSLYAG
58 A NATRO
59 A PIKAA TNVSQA
60 A PIKAA FTIVWSLYAG
61 A PIKAA FTIVWSLYAG
62 A PIKAA TNVSQRKA
63 A PIKAA TNVSDEQA
64 A PIKAA FTIVWSLYAG
65 A PIKAA TNVSQA
66 A PIKAA TNVSQRKA
67 A PIKAA FTIVWSLYAG
68 A PIKAA FTIVWSLYAG
69 A PIKAA TNVSQRKA
70 A PIKAA TNVSDEQA
71 A PIKAA FTIVWSLYAG
72 A PIKAA TNVSDEQRKA
73 A PIKAA LRPEKN
74 A PIKAA VEAN
75 A PIKAA FNHW
76 A PIKAA PD
77 A PIKAA SDT
78 A PIKAA PEYTS
79 A PIKAA EY
80 A PIKAA FTIVWSLYAG
81 A PIKAA TNVSDEQA
82 A PIKAA TNVSQRKA
83 A PIKAA FTIVWSLYAG
84 A PIKAA FTNIVWSLYDEQRKA
85 A PIKAA TNVSQA
86 A PIKAA FTIVWSLYAG
87 A PIKAA FTIVWSLYAG
88 A PIKAA TNVSDEQA
89 A PIKAA TNVSQRKA
90 A PIKAA FTIVWSLYAG
91 A PIKAA FTNIVWSLYDEQRKA
92 A PIKAA TNVSQRKA
93 A PIKAA FTNIVWSLYDEQRKA
94 A PIKAA FTIVWSLYAG
95 A PIKAA TNVSDEQA
96 A PIKAA TNVSDEQA
97 A PIKAA FTIVWSLYAG
98 A PIKAA FTNIVWSLYDEQRKA
99 A PIKAA TNVSQRKA
100 A PIKAA FTNIVWSLYDEQRKA
101 A PIKAA FTIVWSLYAG
102 A PIKAA TNVSDEQA

103 A PIKAA TNVSDEQA
104 A PIKAA FTIVWSLYAG
105 A PIKAA FTNIVWSLYDEQRKA
106 A PIKAA TNVSQRKA
107 A PIKAA FTIVWSLYAG
108 A PIKAA A
109 A PIKAA EADRQ
110 A PIKAA NAKLQ
111 A PIKAA G
112 A PIKAA HNDRQ
113 A PIKAA LVEDK
114 A PIKAA ED
115 A PIKAA TNVSDEQRKA
116 A PIKAA FTIVWSLYAG
117 A PIKAA TNVSQA
118 A PIKAA TNVSQRKA
119 A PIKAA FTIVWSLYAG
120 A PIKAA FTIVWSLYAG
121 A PIKAA TNVSDEQA
122 A PIKAA TNVSQA
123 A PIKAA FTIVWSLYAG
124 A PIKAA TNVSQRKA
125 A PIKAA TNVSQA
126 A PIKAA FTIVWSLYAG
127 A PIKAA FTIVWSLYAG
128 A PIKAA TNVSDEQA
129 A PIKAA TNVSDEQA
130 A PIKAA FTIVWSLYAG
131 A NATRO
132 A PIKAA TNVSQA
133 A PIKAA FTIVWSLYAG
134 A PIKAA FTIVWSLYAG
135 A PIKAA TNVSQA
136 A PIKAA TNVSDEQA
137 A PIKAA FTIVWSLYAG
138 A PIKAA TNVSQA
139 A PIKAA TNVSQRKA
140 A PIKAA FTIVWSLYAG
141 A PIKAA FTIVWSLYAG
142 A PIKAA TNVSQA
143 A PIKAA TNVSDEQA
144 A PIKAA FTIVWSLYAG
145 A PIKAA TNVSQA
146 A PIKAA TNVSDEQRKA
147 A PIKAA TNVSDEQRKAG

**Content of RUC parameter file (RUC.params)**

```
NAME RUC
IO_STRING RUC Z
TYPE LIGAND
AA UNK
ATOM  C8  aroC  X   -0.08
ATOM  C4  aroC  X   -0.08
ATOM  C1  aroC  X   -0.08
ATOM  C2  aroC  X   -0.08
ATOM  C5  aroC  X   -0.08
ATOM  N1  Ntrp  X   -0.57
ATOM  H3  Hpol  X   0.47
ATOM  C10 aroC  X   -0.08
ATOM  C12 aroC  X   -0.08
ATOM  C11 aroC  X   -0.08
ATOM  C6  aroC  X   -0.08
ATOM  C9  COO   X   0.66
ATOM  O1  ONH2  X   -0.51
ATOM  N2  Ntrp  X   -0.57
ATOM  C7  CH2   X   -0.14
ATOM  C3  CH2   X   -0.14
ATOM  H1  Hapo  X   0.13
ATOM  H2  Hapo  X   0.13
ATOM  H4  Hapo  X   0.13
ATOM  H5  Hapo  X   0.13
ATOM  H6  Hpol  X   0.47
ATOM  H8  Haro  X   0.15
ATOM  F1  F     X   -0.21
ATOM  H7  Haro  X   0.15
ATOM  C13 aroC  X   -0.08
ATOM  C16 aroC  X   -0.08
ATOM  C15 aroC  X   -0.08
ATOM  C17 aroC  X   -0.08
ATOM  C14 aroC  X   -0.08
ATOM  H10 Haro  X   0.15
ATOM  H12 Haro  X   0.15
ATOM  C18 CH2   X   -0.14
ATOM  N3  Ntrp  X   -0.57
ATOM  C19 CH3   X   -0.23
ATOM  H16 Hapo  X   0.13
ATOM  H17 Hapo  X   0.13
ATOM  H18 Hapo  X   0.13
ATOM  H15 Hpol  X   0.47
ATOM  H13 Hapo  X   0.13
ATOM  H14 Hapo  X   0.13
ATOM  H11 Haro  X   0.15
```

```
ATOM  H9  Haro  X   0.15
BOND_TYPE  C1   C2  4
BOND_TYPE  C1   C3  1
BOND_TYPE  C1   C4  4
BOND_TYPE  N1   C4  4
BOND_TYPE  N1   C5  4
BOND_TYPE  N1   H3  1
BOND_TYPE  O1   C9  2
BOND_TYPE  C2   C5  4
BOND_TYPE  C2   C6  4
BOND_TYPE  N2   C7  1
BOND_TYPE  N2   C9  1
BOND_TYPE  N2   H6  1
BOND_TYPE  C3   C7  1
BOND_TYPE  C3   H1  1
BOND_TYPE  C3   H2  1
BOND_TYPE  N3   C18 1
BOND_TYPE  N3   C19 1
BOND_TYPE  N3   H15 1
BOND_TYPE  C4   C8  1
BOND_TYPE  C5   C10 4
BOND_TYPE  C6   C9  1
BOND_TYPE  C6   C11 4
BOND_TYPE  C7   H4  1
BOND_TYPE  C7   H5  1
BOND_TYPE  C8   C13 4
BOND_TYPE  C8   C14 4
BOND_TYPE  C10  C12 4
BOND_TYPE  C10  H7  1
BOND_TYPE  C11  C12 4
BOND_TYPE  C11  H8  1
BOND_TYPE  C13  C16 4
BOND_TYPE  C13  H9  1
BOND_TYPE  C14  C17 4
BOND_TYPE  C14  H10 1
BOND_TYPE  C15  C16 4
BOND_TYPE  C15  C17 4
BOND_TYPE  C15  C18 1
BOND_TYPE  C16  H11 1
BOND_TYPE  C17  H12 1
BOND_TYPE  C18  H13 1
BOND_TYPE  C18  H14 1
BOND_TYPE  C19  H16 1
BOND_TYPE  C19  H17 1
BOND_TYPE  C19  H18 1
BOND_TYPE  C12  F1  1
```

```
CHI 1  C15  C18  N3   C19
CHI 2  C13  C8   C4   C1
CHI 3  C16  C15  C18  N3
NBR_ATOM  C8
NBR_RADIUS 7.842337
ICOOR_INTERNAL   C8    0.000000    0.000000    0.000000   C8    C4    C1
ICOOR_INTERNAL   C4    0.000000  180.000000    1.445007   C8    C4    C1
ICOOR_INTERNAL   C1   -0.000000   49.296676    1.378582   C4    C8    C1
ICOOR_INTERNAL   C2  179.847859   72.459130    1.429139   C1    C4    C8
ICOOR_INTERNAL   C5    1.421276   73.244984    1.398909   C2    C1    C4
ICOOR_INTERNAL   N1   -1.390885   72.601175    1.375455   C5    C2    C1
ICOOR_INTERNAL   H3 -179.649669   55.435909    1.010614   N1    C5    C2
ICOOR_INTERNAL   C10 -179.812702   56.962350    1.398479   C5    C2    N1
ICOOR_INTERNAL   C12    0.266108   62.461209    1.394085   C10   C5    C2
ICOOR_INTERNAL   C11    0.499098   59.063143    1.393157   C12   C10   C5
ICOOR_INTERNAL   C6   -0.622982   58.698116    1.409650   C11   C12   C10
ICOOR_INTERNAL   C9  176.353215   61.871368    1.484207   C6    C11   C12
ICOOR_INTERNAL   O1  -20.624404   61.971761    1.232903   C9    C6    C11
ICOOR_INTERNAL   N2 -179.263291   58.677876    1.382317   C9    C6    O1
ICOOR_INTERNAL   C7   -7.998563   50.588441    1.455300   N2    C9    C6
ICOOR_INTERNAL   C3   67.006167   67.530499    1.523069   C7    N2    C9
ICOOR_INTERNAL   H1  164.499103   71.790637    1.095996   C3    C7    N2
ICOOR_INTERNAL   H2 -117.481758   69.238629    1.096390   C3    C7    H1
ICOOR_INTERNAL   H4 -124.333099   71.788720    1.096017   C7    N2    C3
ICOOR_INTERNAL   H5 -115.761455   73.681546    1.095471   C7    N2    H4
ICOOR_INTERNAL   H6 -173.748863   64.931578    1.014929   N2    C9    C7
ICOOR_INTERNAL   H8  179.839814   61.928760    1.087596   C11   C12   C6
ICOOR_INTERNAL   F1  179.479249   60.507024    1.339698   C12   C10   C11
ICOOR_INTERNAL   H7  179.890648   58.508816    1.084455   C10   C5    C12
ICOOR_INTERNAL   C13   90.076839   60.101978    1.393592   C8    C4    C1
ICOOR_INTERNAL   C16 -179.996924   60.108661    1.395105   C13   C8    C4
ICOOR_INTERNAL   C15    0.002292   60.005592    1.395096   C16   C13   C8
ICOOR_INTERNAL   C17    0.000449   59.982924    1.393731   C15   C16   C13
ICOOR_INTERNAL   C14    0.002395   59.967385    1.394952   C17   C15   C16
ICOOR_INTERNAL   H10  179.984574   60.820114    1.087182   C14   C17   C15
ICOOR_INTERNAL   H12  179.914903   59.259908    1.087540   C17   C15   C14
ICOOR_INTERNAL   C18  179.995067   60.017449    1.490982   C15   C16   C17
ICOOR_INTERNAL   N3  -90.008994   67.647806    1.441424   C18   C15   C16
ICOOR_INTERNAL   C19  179.557268   68.480178    1.462116   N3    C18   C15
ICOOR_INTERNAL   H16  -60.889480   68.832569    1.094491   C19   N3    C18
ICOOR_INTERNAL   H17 -118.595374   70.617562    1.094420   C19   N3    H16
ICOOR_INTERNAL   H18 -119.275213   68.567878    1.094949   C19   N3    H17
ICOOR_INTERNAL   H15  120.598875   70.108159    1.020400   N3    C18   C19
ICOOR_INTERNAL   H13  122.413879   71.220154    1.098268   C18   C15   N3
ICOOR_INTERNAL   H14  114.674094   70.965289    1.098284   C18   C15   H13
ICOOR_INTERNAL   H11  179.836710   60.753206    1.087319   C16   C13   C15
```

ICOOR_INTERNAL    H9   179.901536   59.042950    1.087435   C13   C8    C16

**Content of constratins file (constraints_helix.cst)**
AtomPair N1 1X OD1 131A HARMONIC 3.0 0.3
AtomPair N2 1X OE1 54A HARMONIC 3.0 0.3
AtomPair N3 1X OE1 22A HARMONIC 3.0 0.3

**Content of netcharge file (helix_netcharge.charge)**
DESIRED_CHARGE -5
PENALTIES_CHARGE_RANGE -10 -1
PENALTIES 10 0 0 0 0 0 0 0 0 10
BEFORE_FUNCTION QUADRATIC
AFTER_FUNCTION QUADRATIC

**Content of composition file (helix_composition.comp)**
PENALTY_DEFINITION
TYPE THR
DELTA_START 0
DELTA_END 1
PENALTIES 0 100
ABSOLUTE 12
BEFORE_FUNCTION CONSTANT
AFTER_FUNCTION QUADRATIC
END_PENALTY_DEFINITION

PENALTY_DEFINITION
TYPE GLY
DELTA_START 0
DELTA_END 1
PENALTIES 0 100
ABSOLUTE 4
BEFORE_FUNCTION CONSTANT
AFTER_FUNCTION QUADRATIC
END_PENALTY_DEFINITION

PENALTY_DEFINITION
TYPE SER
DELTA_START 0
DELTA_END 1
PENALTIES 0 100
ABSOLUTE 11
BEFORE_FUNCTION CONSTANT
AFTER_FUNCTION QUADRATIC
END_PENALTY_DEFINITION

PENALTY_DEFINITION

TYPE ASN
DELTA_START 0
DELTA_END 1
PENALTIES 0 100
ABSOLUTE 10
BEFORE_FUNCTION CONSTANT
AFTER_FUNCTION QUADRATIC
END_PENALTY_DEFINITION

PENALTY_DEFINITION
TYPE ARG
DELTA_START 0
DELTA_END 1
PENALTIES 0 100
ABSOLUTE 12
BEFORE_FUNCTION CONSTANT
AFTER_FUNCTION QUADRATIC
END_PENALTY_DEFINITION

PENALTY_DEFINITION
TYPE ALA
DELTA_START 0
DELTA_END 1
PENALTIES 0 100
ABSOLUTE 21
BEFORE_FUNCTION CONSTANT
AFTER_FUNCTION QUADRATIC
END_PENALTY_DEFINITION

PENALTY_DEFINITION
TYPE TRP
DELTA_START -1
DELTA_END 1
PENALTIES 100 0 100
ABSOLUTE 1
BEFORE_FUNCTION CONSTANT
AFTER_FUNCTION QUADRATIC
END_PENALTY_DEFINITION

PENALTY_DEFINITION
TYPE LYS
DELTA_START 0
DELTA_END 1
PENALTIES 0 100
ABSOLUTE 13
BEFORE_FUNCTION CONSTANT

AFTER_FUNCTION QUADRATIC
END_PENALTY_DEFINITION

PENALTY_DEFINITION
TYPE GLN
DELTA_START 0
DELTA_END 1
PENALTIES 0 100
ABSOLUTE 12
BEFORE_FUNCTION CONSTANT
AFTER_FUNCTION QUADRATIC
END_PENALTY_DEFINITION

PENALTY_DEFINITION
TYPE TYR
DELTA_START 0
DELTA_END 1
PENALTIES 0 100
ABSOLUTE 8
BEFORE_FUNCTION CONSTANT
AFTER_FUNCTION QUADRATIC
END_PENALTY_DEFINITION

PENALTY_DEFINITION
TYPE GLU
DELTA_START 0
DELTA_END 1
PENALTIES 0 100
ABSOLUTE 15
BEFORE_FUNCTION CONSTANT
AFTER_FUNCTION QUADRATIC
END_PENALTY_DEFINITION

PENALTY_DEFINITION
TYPE ASP
DELTA_START 0
DELTA_END 1
PENALTIES 0 100
ABSOLUTE 11
BEFORE_FUNCTION CONSTANT
AFTER_FUNCTION QUADRATIC
END_PENALTY_DEFINITION

PENALTY_DEFINITION
TYPE VAL
DELTA_START 0

DELTA_END 1
PENALTIES 0 100
ABSOLUTE 15
BEFORE_FUNCTION CONSTANT
AFTER_FUNCTION QUADRATIC
END_PENALTY_DEFINITION
